# Rewiring DNA repair with PARP-based chemical inducers of proximity

**DOI:** 10.1101/2025.07.26.666954

**Authors:** Bryce da Camara, Eric M. Bilotta, Erin F. Broderick, Ashwini Premashankar, Paige A. Barta, Trever R. Carter, Lauren M. Hargis, Daniel Durocher, Michael A. Erb

## Abstract

Chemical inducers of proximity (CIPs) can elicit durable—and often neomorphic—biological effects through the formation of a ternary complex, even at low equilibrium occupancy of their targets. This “event-driven” pharmacology is exemplified by CIPs that promote targeted protein degradation, but other applications remain underexplored. We developed a generalizable strategy to discover event-driven CIPs by tracking the cellular effects of heterobifunctional small molecules alongside quantitative measures of intracellular target engagement. Using this approach, we discovered PCIP-1, which inhibits DNA repair by recruiting BET proteins to PARP2. Unlike conventional PARP inhibitors, PCIP-1 activity is observed at low equilibrium occupancy of PARP1/2 and without inhibition of PARP-catalyzed PARylation, yet it retains synthetic lethality in cancer cells with homologous recombination deficiencies. *PARP1* knockout, which confers resistance to conventional PARP drugs, increases sensitivity to PCIP-1, offering a potential new mechanism to overcome clinical resistance. Through these studies, we demonstrate that DNA repair can be rewired by CIPs and introduce a new form of event-driven pharmacology.

## Introduction

Conventional classes of small-molecule drugs and chemical probes function through “occupancy-driven” mechanisms of action, typically requiring high equilibrium engagement of their targets to elicit a biological effect. In contrast, chemical inducers of proximity (CIPs) can show “event-driven” mechanisms of action, in which a biological effect is elicited through ternary complex formation, despite low equilibrium target engagement.^1–4^ This phenomenon is well-demonstrated by CIPs that can induce targeted protein degradation (TPD) at sub-stoichiometric concentrations by inducing transient interactions with components of the ubiquitin-proteasome system.^3,5^ Through their distinct mechanisms of action, CIPs can often produce gain-of-function and neomorphic biological effects that are inaccessible to conventional small molecules. This was first shown using genetically encoded systems that enabled chemically induced proximity,^1,2^ with index examples demonstrating that proximity pharmacology can control the activation of cell surface receptors, alter subcellular localization, and create neomorphic transcriptional regulatory complexes.^6–11^

Heterobifunctional small molecules enable the rational design of CIPs that control diverse biological processes.^1,2^ Recent examples have shown that heterobifunctional small molecules can control ubiquitination,^5,12^ phosphorylation,^13–15^ acetylation,^16,17^ O-GlcNAcylation,^18–20^ methylation,^21^ endocytosis,^22–24^ RNA hydrolysis,^25,26^ transcription,^27,28^ and subcellular localization,^29,30^ among other effects.^31^ However, given the matrix of interactions that could, in principle, be promoted between any two targets, the potential outcomes of proximity pharmacology are likely far more expansive than currently appreciated.

Due in part to the difficulty of predicting and measuring the outcomes of induced proximity beyond TPD, other applications of proximity pharmacology remain underexplored. However, we reasoned that the ability of CIPs to elicit event-driven pharmacological effects despite low fractional engagement of their targets could enable a generalizable strategy to discover new classes of heterobifunctional CIPs. Using pre-existing ligands with well-established cellular targets to synthesize new classes of heterobifunctional small molecules, we hypothesized that quantitative measurements of cellular target engagement could be tracked alongside diverse readouts of biological activity to identify new CIPs that show event-driven pharmacological behavior.

We selected the poly(ADP-ribose) polymerase (PARP) family of DNA repair proteins for an initial proof of concept, as DNA repair pathways have yet to be the subject of CIP-induced rewiring. PARP1/2 inhibitors are approved by the FDA to target homologous recombination (HR)-deficient cancers, a paradigmatic example of synthetic lethality caused by the increased sensitivity of these tumors to the inhibition of PARylation and the trapping of PARP1 onto sites of DNA damage.^32–35^ We synthesized a series of heterobifunctional small molecules attaching one such inhibitor, rucaparib, to JQ1, a chemical probe that binds to the bromodomains of the BET family transcriptional co-activators (BRD2/3/4).^36,37^ Prior research has demonstrated that both small molecules can be used to synthesize heterobifunctional compounds,^38–40^ and since the bromodomains of BET proteins function primarily to mediate protein localization, JQ1 can be used to recruit BRD4 to new locations in the cell while preserving its native protein-protein interactions (PPIs) and functions.^27,28,41,42^ We reasoned that JQ1 could therefore be used to synthesize PARP-based CIPs (PCIPs) that would recruit functioning BET family proteins to PARP1/2, potentially altering DNA repair pathways. However, the potential consequences of inducing spatial proximity between PARP and BET proteins was not obvious. Thus, we used cell viability assays as a coarse-grained readout of biological activity, which we tracked alongside measurements of intracellular target engagement to identify compounds that were active at low equilibrium occupancy of their targets.

Through these studies, we identified PCIP-1, which shows potent anti-proliferative activity, despite low fractional engagement of PARP and BET proteins. Unlike conventional PARP inhibitors, which inhibit PARP1/2-catalyzed PARylation, PCIP-1 inhibits DNA repair by recruiting BET proteins to PARP2, resulting in non-overlapping mechanisms of resistance but preserving synthetic lethality with (HR)-deficiencies. In fact, *PARP1* knockout cells, which are resistant to PARP inhibition, show increased sensitivity to PCIP-1, suggesting that PCIPs may represent a new modality to target tumors with acquired resistance to PARP inhibition. Altogether, our study provides a generalizable framework to expand the scope of proximity pharmacology, demonstrates the ability to rewire DNA repair pathways with CIPs, and establishes a new mechanism to target HR-deficient tumors through synthetic lethality.

## Results

### A strategy to identify event-driven CIP activity

Candidate PCIPs were designed from JQ1 and rucaparib, attaching variable linkers at sites on each molecule informed by previously disclosed PROTACs (**Fig. 1a**).^39,40^ Without a clear expectation for the biological effects that could be provoked by the chemically induced proximity of PARP and BET proteins, we began by profiling their effects on cellular viability as a granular readout of diverse biological activities. However, because PARP inhibitors and BET bromodomain inhibitors can elicit anti-proliferative effects independently, we sought a strategy to differentiate between proximity-induced effects and occupancy-driven inhibition of either target. We hypothesized that any biological effects resulting from low fractional occupancy might reflect event-driven CIP activity. To test this, we developed a workflow to quantitatively compare the anti-proliferative effects of each PCIP relative to their intracellular target engagement profiles.

**Fig. 1.**
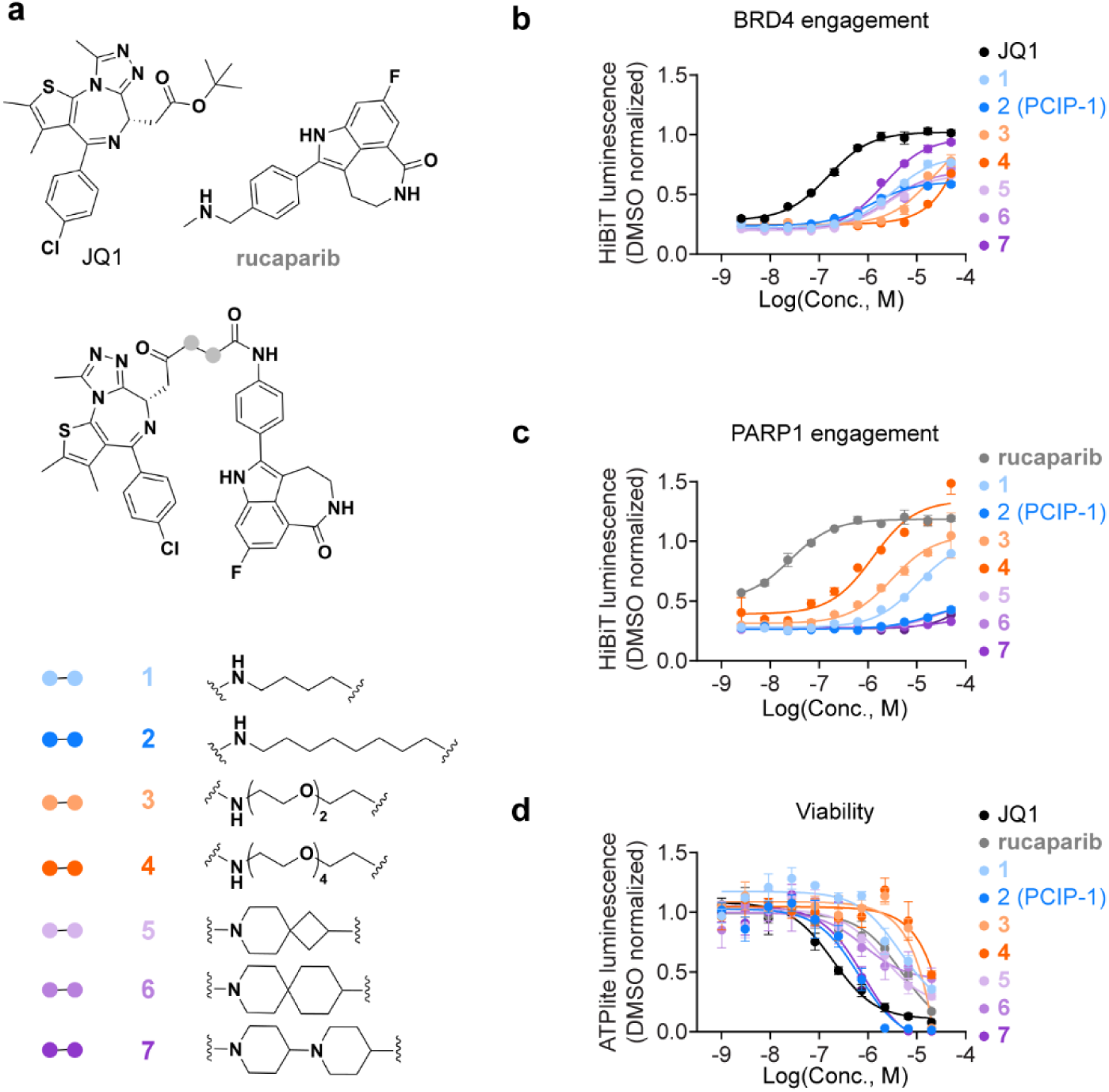
Discovery of PCIP-1. **a**, Chemical structures of candidate PCIPs. **b,** BRD4 target engagement measured by competitive displacement of dBET6. Jurkat-BRD4-HiBit cells are treated with test compounds for 2 h and then for 45 min with 500 nM dBET6 (HiBiT luminescence normalized to DMSO control with no dBET6, *n* = 3, mean ± s.e.m.). **c,** PARP1 target engagement measured by competitive displacement of SK-575. Jurkat-PARP1-HiBit cells are treated with test compounds for 2 h and then for 4 h with 250 nM SK-575 (HiBiT luminescence normalized to DMSO control with no SK-575, *n* = 3, mean ± s.e.m.). **d,** Jurkat viability after 72 h compound treatment (ATPlite luminescence normalized to DMSO, *n* = 3, mean ± s.e.m.).

We used competitive PROTAC displacement assays to measure BRD4 and PARP1 target engagement in living cells.^43,44^ In these assays, cells were pretreated with the candidate PCIPs and then exposed to a fixed concentration of a PROTAC, such that target engagement by the PCIP can be measured by the blockade of PROTAC-induced degradation. These measurements were enabled by fusing HiBiT to the carboxy (C)-termini of PARP1 and BRD4 using CRISPR/Cas9-based endogenous genome editing, allowing for the highly sensitive detection of protein abundance through a luminescence-based protein complementation assay.^45,46^ BRD4 and PARP1 HiBiT reporters were potently degraded by the PROTACs, dBET6 and SK-575,^47,48^ respectively, which was blocked by increasing concentrations of JQ1 or rucaparib, altogether validating these assays for making quantitative measurements of intracellular target engagement (**Fig. 1b,c**).

Next, we profiled all PCIP candidates and their parent inhibitors for BRD4 target engagement, PARP1 target engagement, and cell viability effects in the Jurkat T-cell acute lymphoblastic leukemia (T-ALL) cell line (**Fig. 1b-d**). In these assays, JQ1 and rucaparib produced anti-proliferative effects at high fractional occupancy of BRD4 and PARP1, respectively, consistent with their occupancy-driven mechanisms of action. In contrast, several PCIP candidates showed potent anti-proliferative effects despite weak engagement of BRD4 and PARP1. The most profound effect was seen for **2** (PCIP-1), which inhibits cell viability at similar concentrations as JQ1 (IC_50_: JQ1 = 193 nM, PCIP-1 = 662 nM) but requires much higher concentrations to engage BRD4 in cells (EC_50_: JQ1 = 97 nM, PCIP-1 = 26 µM) (**Fig. 1b,d**). In fact, PCIP-1 failed to saturate BRD4 even at the highest concentration tested, 50 µM (**Fig. 1b**), altogether ruling out the possibility that its anti-proliferative effects can be explained by occupancy-driven inhibition of BET bromodomain proteins. Likewise, despite failing to show any detectable engagement of PARP1, PCIP-1 inhibited the viability of Jurkat cells more potently than rucaparib (**Fig. 1c,d**), demonstrating that it does not act through an occupancy-driven mechanism of PARP inhibition either.

Initially, we attributed the weak engagement of BRD4 by several PCIPs to poor cell penetration, as these compounds maintained relatively strong engagement of purified recombinant BRD4 bromodomain 1 (BD1) *in vitro* **(Extended Data Fig. 1a**). However, the relative rank in target engagement potencies among the candidate PCIPs differed between BRD4 and PARP1, suggesting a more complex explanation. For example, **4**, is among the worst at engaging BRD4 but the best at engaging PARP1. Likewise, while **7** is the most potent engager of BRD4 in cells, it is among the worst for PARP1. The source of these differences is not immediately clear but might plausibly be related to the ability of ternary complex formation to stabilize ligand binding through cooperativity.

### PCIP-1 mediates ternary complex formation with PARP and BET proteins

To assess if the activity of PCIP-1 is potentially mediated by the formation of a ternary complex, we synthesized the enantiomer of PCIP-1, which incorporates a stereochemical inversion within the JQ1 moiety that disrupts binding to BET bromodomains (**Fig. 2a**).^36^ As expected, this control compound, *ent*-PCIP-1 (**8**), is unable to engage BRD4 in cells and *in vitro* and engages PARP1 to the same degree as PCIP-1 (**Fig. 2b,c, Extended Data Fig 1a.**). It also showed no impact on Jurkat viability (**Fig. 2d**), indicating that the activity of PCIP-1 is dependent on its ability to co-opt BET proteins. These data suggested that PCIP-1 might be able to induce the formation of a ternary complex at low fractional engagement of PARP and BET proteins, eliciting a loss of cell viability through an event-driven mechanism of action (MoA).

**Fig. 2.**
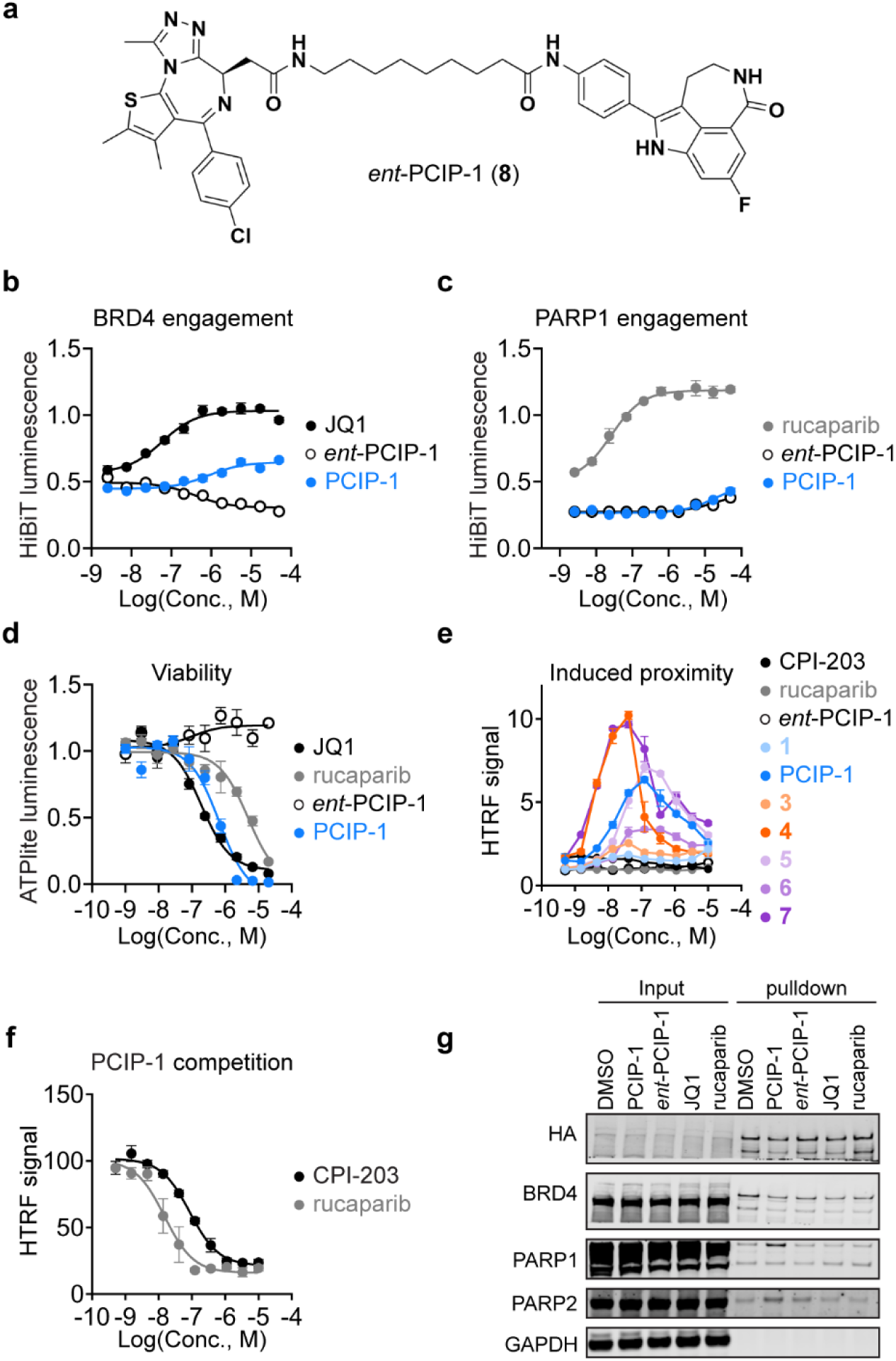
PCIP-1 induces ternary complex formation. **a**, Chemical structure of *ent*-PCIP-1. **b,** BRD4 target engagement measured by displacement of dBET6 (DMSO-normalized HiBiT luminescence, *n* = 3, mean ± s.e.m.). **c,** PARP1 target engagement measured by displacement of SK-575 (DMSO-normalized HiBiT luminescence, *n* = 3, mean ± s.e.m.). **d,** Jurkat viability after 72-h compound treatment (ATPlite luminescence normalized to DMSO, *n* = 3, mean ± s.e.m.). Data in Fig. 1d is taken from the same experiment (PCIP-1, rucaparib, and JQ1 are repeated in both panels). **e,** Compound-induced FRET between 6xHis-BRD4 bromodomain 1 and biotin-PARP1 catalytic domain (DMSO-normalized, *n* = 4, mean ± s.e.m.). **f,** Inhibition of HTRF signal (normalized by percent activity remaining) induced by 500 nM PCIP-1 (*n* = 4, mean ± s.e.m.). **g,** Co-IP of BRD4-2xHA following treatment of 22Rv1 cells with 1 µM of each indicated compound for 4 h.

To assess its potential for ternary complex formation, we developed a homogenous time-resolved (TR)-FRET (HTRF) assay for the catalytic domain of PARP1 and the first bromodomain of BRD4 (**Fig. 2e**). PCIP-1 induced a concentration-dependent increase in HTRF signal with a characteristic “hook effect” that is commonly observed in ternary complex equilibria (**Fig. 2e**).^49^ PCIP-1-induced FRET was suppressed in a dose-dependent manner by the addition of rucaparib or the BET bromodomain inhibitor, CPI-203, indicating that the HTRF assay reports on an authentic induced proximity effect (**Fig. 2f**). In contrast to PCIP-1, *ent*-PCIP-1 was inactive across all concentrations tested (**Fig. 2e**), consistent with the inability to bind BRD4. To determine whether a ternary complex is formed in cells, we performed a co-immunoprecipitation (co-IP) experiment, which revealed an increased association between BRD4 and PARP1/2 in the presence of PCIP-1 but not in the presence of *ent*-PCIP-1, rucaparib, or JQ1 (**Fig. 2g**). Based on these data, we concluded that PCIP-1 is capable of inducing proximity between BET and PARP proteins, despite its weak equilibrium occupancy of these targets in cells.

### PCIP-1 inhibits the repair of DNA damage

To gain a preliminary understanding of the biological processes underlying PCIP-1-induced anti-proliferative effects, we sought to determine whether PCIP-1 impacts the function of PARP1/2. To begin, we repeated the Jurkat cell viability assays in the presence of the DNA alkylating compound, methyl methanesulfonate (MMS), which is known to sensitize cancer cells to PARP1/2 inhibition.^50^ We found that the activities of PCIP-1 and rucaparib are improved in the presence of MMS, whereas JQ1 activity is unchanged, likely suggesting that PCIP-1 interferes with DNA damage repair (**Fig. 3a**). Notably, MMS does not change PCIP-1 engagement of BRD4 or PARP1 in cells, indicating that its enhanced activity does not arise from changes in equilibrium occupancy **(Extended Data Fig. 1b,c)**. By immunoblot analysis of the DNA damage marker, γH2AX, we confirmed that PCIP-1 inhibits the repair of MMS-induced DNA damage (**Fig. 3b**).^51,52^ As expected, this was also observed for rucaparib, but not for JQ1 or *ent*-PCIP-1, altogether indicating that PCIP-1 interferes with PARP1/2-dependent DNA repair pathways through a bifunctional mechanism of action. Cell cycle analysis further supported an impact on DNA repair by demonstrating that PCIP-1 induces a strong S-phase arrest, which is comparable to the effects of rucaparib but not observed for its associated control compounds (**Fig. 3c**). Additionally, we observed markers of apoptosis, including dose-responsive and time-dependent increases in the levels of cleaved PARP1/2 and cleaved Caspase 3, both of which tracked with the accumulation of γH2AX (**Fig. 3d, Extended Data Fig. 1d,e**). However, whereas rucaparib potently inhibits PARP-catalyzed PARylation, PCIP-1 showed minimal effects on global PARylation levels (**Fig. 3b, Extended Data Fig. 1d**), differentiating its cellular mechanism of action from conventional PARP inhibitors and further confirming that it does not function through the occupancy-based inhibition of PARP proteins.

**Fig. 3.**
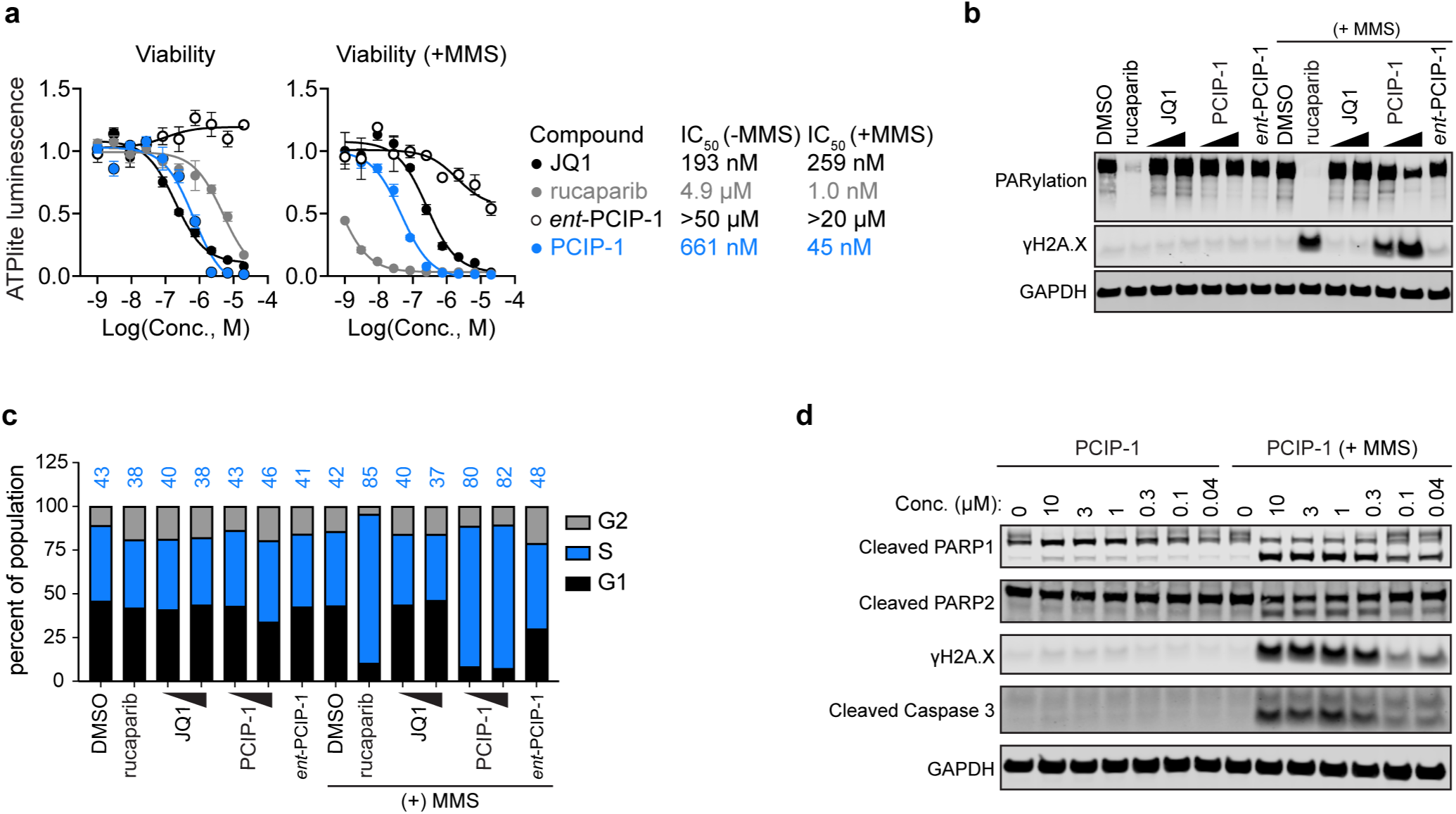
PCIP-1 inhibits DNA damage repair. **a**, Jurkat viability after 72 h compound treatment with and without 25 µM MMS (ATPlite luminescence normalized to DMSO, *n* = 3, mean ± s.e.m.). Data in Fig. 1d. is taken from the same experiment (PCIP-1, rucaparib, *ent-*PCIP-1, and JQ1 are repeated in both panels). **b,** Immunoblot following 24 h treatment of Jurkat cells. PCIP-1 and JQ1 were used at 100 nM and 1 µM. All other compounds were used at 1 µM, with and without 25 µM MMS. **c,** Cell cycle analysis of Jurkat cells treated with indicated drugs for 24 h, JQ1 and PCIP-1 treated at 100 nM and 1 µM, all other compounds at 1 µM, with and without 25 µM MMS. **d,** Immunoblot analysis of Jurkat cells treated with PCIP-1 for 24 h, with and without 25 µM MMS.

### PCIP-1 activity is mediated by the recruitment of BET proteins to PARP2

In total, our data suggested that PCIP-1 inhibits DNA repair through an event-driven pharmacology. To formally assess the requirement for PARP1/2 engagement in this MoA, we performed competitive growth experiments in which Cas9-expressing cells are transduced with a bicistronic vector encoding an sgRNA (single-guide RNA) for targeted gene disruption and EGFP (enhanced green fluorescent protein) for tracking the competitive fitness of sgRNA-positive cells relative to untransduced cells in the same population.^53^ Conventional PARP inhibitors function through a dominant-negative MoA that involves the trapping of PARP1 onto sites of DNA damage such that *PARP1* knockout confers resistance to PARP inhibitors.^50^ As expected, treatment with the PARP inhibitor talazoparib selected for Jurkat cells transduced with each of the 3 distinct sgRNAs targeting *PARP1* in our competitive growth assays (**Fig. 4a, Extended Data Fig. 2a**). In contrast, PCIP-1 had the opposite effect, rapidly eliminating cells expressing *PARP1*-targeted sgRNAs from the population. This selective pressure was not exerted by JQ1 or *ent*-PCIP-1, suggesting that PCIP-1 functions through a mechanism distinct from both PARP inhibitors and BET bromodomain inhibitors.

**Fig. 4.**
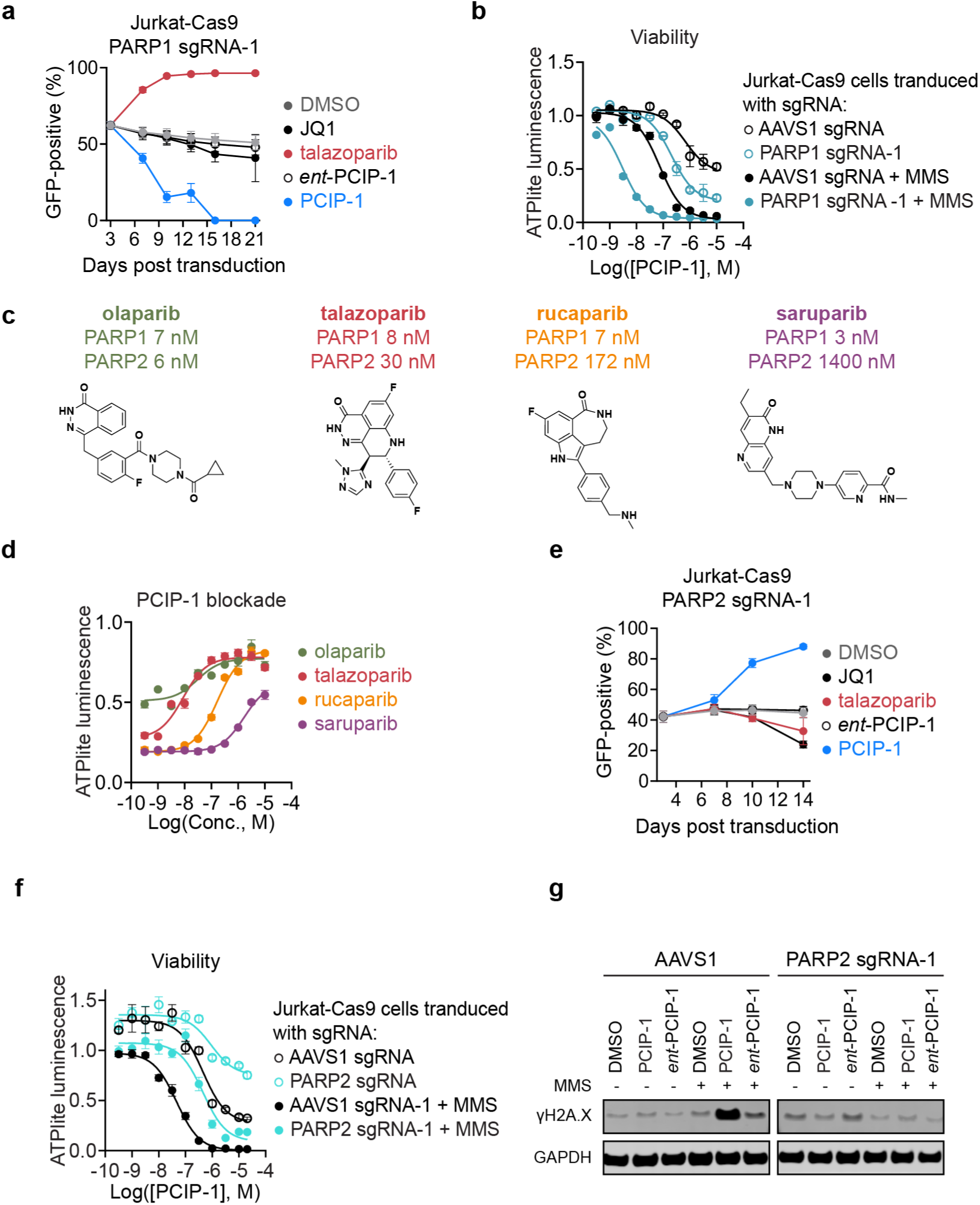
PCIP-1 anti-proliferative effects are mediated by PARP2. **a**, Competitive growth assay testing the effect of *PARP1*-sgRNA-1 on the response of Jurkat-Cas9 cells to talazoparib (1 µM), PCIP-1 (1 µM), and *ent*-PCIP-1 (1 µM), or JQ1 (250 nM). Proportion of GFP-positive cells over time was measured by flow cytometry (*n* = 3, mean ± s.e.m.). **b,** Viability assay (72 h) in Jurkat-Cas9 cells with and without 25 µM MMS (ATPlite luminescence normalized to DMSO, *n* = 3, mean ± s.e.m.). **c)** Chemical structures of PARP1/2 inhibitors and their corresponding IC_50_ values for PARP1/2 selectivity. **d,** Viability assay (72 h) in Jurkat-Cas9 cells with and without 25 µM MMS, cell cultures treated with 1 µM PCIP-1 (ATPlite luminescence normalized to DMSO, *n* = 3, mean ± s.e.m.). **e,** Competitive growth assay testing the effect of *PARP2*-sgRNA-1 on the response of Jurkat-Cas9 cells to 1 µM talazoparib, PCIP-1, *ent*-PCIP-1, and JQ1. Proportion of GFP-positive cells over time was measured by flow cytometry (*n* = 3, mean ± s.e.m.). **f,** Viability assay (72 h) in Jurkat-Cas9 cells with and without 25 µM MMS (ATPlite luminescence normalized to DMSO, *n* = 3, mean ± s.e.m.). **g,** Immunoblot analysis of Jurkat-CAS9 cells after 100 nM compound treatment after 24h, with and without 25 µM MMS.

To enable dose-response assays, we isolated a population of *PARP1* knockout cells by selecting sgRNA-positive cells to homogeneity with talazoparib (1 µM). Loss of *PARP1* expression in these cells was validated by immunoblot analysis and by their resistance to multiple distinct PARP inhibitors **(Extended Data Fig. 2b-f)**.

Consistent with the previous competitive growth experiments, cell viability assays revealed that PCIP-1 potency is improved by more than 10-fold in these *PARP1* loss-of-function cells compared to control cells harboring an sgRNA targeting the *AAVS1* safe-harbor locus (**Fig. 4b, Extended Data Fig. 2g)**. These data, which revealed that *PARP1* knockout cells are more sensitive to PCIP-1—despite being resistant to conventional PARP inhibitors—further highlight the differentiated mechanism of action by which PCIP-1 inhibits cell viability.

Based on the divergent effects of PCIP-1 and conventional PARP inhibitors in response to *PARP1* knockout, we hypothesized that PCIP-1 might function through PARP2 instead of PARP1. To test this hypothesis, we evaluated whether saturating the PARP2 ligand-binding site with excess talazoparib could block the anti-proliferative effects of PCIP-1 in *PARP1* knockout cells. Indeed, co-treating with talazoparib (1 µM) fully blocked the anti-proliferative effects of PCIP-1 in each of the 3 distinct *PARP1*-deficient Jurkat populations **(Extended Data Fig. 3a)**. In total, we tested 4 PARP inhibitors—olaparib, talazoparib, rucaparib, and saruparib—for their ability to block the effects of PCIP-1. These 4 structurally distinct ligands share single-digit-nanomolar potency for PARP1, but their potencies for PARP2 vary dramatically, ranging from 6 nM to 1.4 µM (**Fig. 4c**).^54^ All four compounds blocked the effects of PCIP-1 in a dose-responsive manner and at concentrations that tracked with their respective PARP2 potencies (**Fig. 4c,d**), indicating that PCIP-1 functions through PARP2.

To validate these findings further, we sought to establish *PARP1* loss-of-function cells through an alternative experimental procedure, using electroporation to introduce a CRISPR/Cas9 ribonucleoprotein (RNP) complex targeting the *PARP1* gene into wild-type Jurkat cells (using the sgRNA-1 sequence previously used in competitive growth assays). We treated the electroporated cells for 14 days with 1 µM talazoparib to select for insertion-deletion alterations (indels) causing *PARP1* loss-of-function, which was confirmed by TIDE (tracking of indels by decomposition) and was functionally validated by resistance to 4 distinct PARP inhibitors **(Extended Data Fig. 4a,b)**.^55^ Consistent with previous results, these cells are more than 10-fold sensitized to PCIP-1 compared to wild-type Jurkat cells **(Extended Data Fig. 4c)**, and the anti-proliferative effects of PCIP-1 can be blocked by co-treatment with 1 µM talazoparib **(Extended Data Fig. 4d)**. Once again, we observed that the blockade of PCIP-1-induced cell viability effects by each of the four PARP inhibitors corresponded with their relative PARP2 potencies **(Extended Data Fig. 4e)**.

These data indicated that PCIP-1 functions through the recruitment of BET proteins specifically to PARP2. To test this orthogonally, we performed competitive growth assays evaluating whether *PARP2* loss-of-function can confer resistance to PCIP-1. Indeed, PCIP-1 rapidly selected for Jurkat-Cas9 cells transduced with sgRNAs targeting *PARP2*, which was not observed with JQ1, talazoparib, or *ent*-PCIP-1 (**Fig. 4e**). This experiment was performed with 3 additional sgRNAs targeting *PARP2*, producing the same results in each instance **(Extended Data Fig. 5a)**. After completion of the competitive growth assay, Jurkat-Cas9 cells that had been transduced with *PARP2*-targeting sgRNAs and selected with PCIP-1 were expanded and confirmed to harbor *PARP2* knockout by immunoblot analysis **(Extended Data Fig. 5b)**. These were then used for dose-response assays, which demonstrated decreased sensitivity to the anti-proliferative effects of PCIP-1 (**Fig. 4f, Extended Data Fig. 5c,d**). Notably, the resistance of *PARP2* knockout cells to PCIP-1 tracked with a decrease in the ability of PCIP-1 to inhibit the repair of DNA damage (**Fig. 4g, Extended Data Fig. 5e**). Finally, we recapitulated these results in Jurkat cells electroporated with CRISPR/Cas9 RNPs targeting the *PARP2* gene using an sgRNA distinct from the 4 used for competitive growth assays **(Extended Data Fig. 6a,b)**.

Altogether, these data demonstrate that PCIP-1 elicits toxicity through the recruitment of BET proteins to PARP2, as PCIP-1 induced toxicity can be rescued *via* genetic depletion of *PARP2* or chemical blockade of its ligand-binding site. However, since we had not previously measured PARP2 engagement—only PARP1—it was possible that PCIP-1 might induce a ternary complex that drives occupancy-based inhibition of PARP2 rather than an event-driven pharmacology. This would be conceptually similar to rapamycin, FK506, cyclosporin A, and other drugs that inhibit enzymes by forming a ternary complex with cyclophilin or immunophilin proteins at high equilibrium occupancy.^56–60^ We tested PARP2 engagement using a cell-based NanoBRET assay that reports on the competitive displacement of a fluorescent tracer from the PARP2 ligand-binding site, finding that PCIP-1 is unable to stably engage PARP2 in cells **(Extended Data Fig. 6c)**. Thus, we can conclude that the recruitment of BET proteins to PARP2 at low equilibrium occupancy produces an as-yet undefined but durable event-driven effect that results in the inhibition of DNA damage repair.

### Synthetic lethality between PCIP-1 and HR deficiency

Since PCIP-1 inhibits the repair of DNA damage through a mechanism that is differentiated from conventional PARP inhibitors, we sought to determine if it remained synthetically lethal with genetic alterations that cause HR deficiencies. To test this, we characterized its effects in DLD1, HCT116, and RPE1-TP53^KO^ cell lines with and without knockout of *BRCA1 or BRCA2*, which are commonly used to test the effects of PARP inhibitors and other agents exploiting synthetic lethality with HR deficiencies.^61–66^ In DLD1 cells, rucaparib and PCIP-1 show little effect on the viability of parental cells. However, we found that *BRCA2^KO^* cells are highly sensitive to PCIP-1, showing more than 100-fold sensitization compared to parental cells (**Fig. 5a**). These *BRCA2^KO^* cells are also highly sensitive to rucaparib, but its effects were less potent than PCIP-1. The effects of JQ1 were unaltered by *BRCA2* knockout and *ent*-PCIP-1 did not impact the viability of parental or *BRCA2^KO^* cells. We observed similar effects to varying degrees in HCT116 cells and RPE1-TP53^KO^ cells (**Fig. 5b,c**), further confirming the synthetic lethality between PCIP-1 and HR-deficient tumors. The RPE1 model was previously used to discover a mechanism of resistance to PARP inhibitors involving the restoration of HR through the knockout of *53BP1*.^67,68^ As expected, *53BP1* knockout successfully reversed rucaparib-induced cytotoxicity, but it only partially reduced the anti-proliferative effects of PCIP-1, further reinforcing that PCIP-1 acts through a differentiated mechanism of action compared to conventional PARP inhibitors. These results suggest PCIPs may provide an opportunity to overcome resistance to PARP inhibition that occurs through the restoration of HR.^69,70^ Combined with our observation that *PARP1* knockout cells are hypersensitive to PCIP-1, these data suggest that PCIPs may be able to address two of the predominant pathways for acquired resistance to PARPi therapy.

**Fig. 5.**
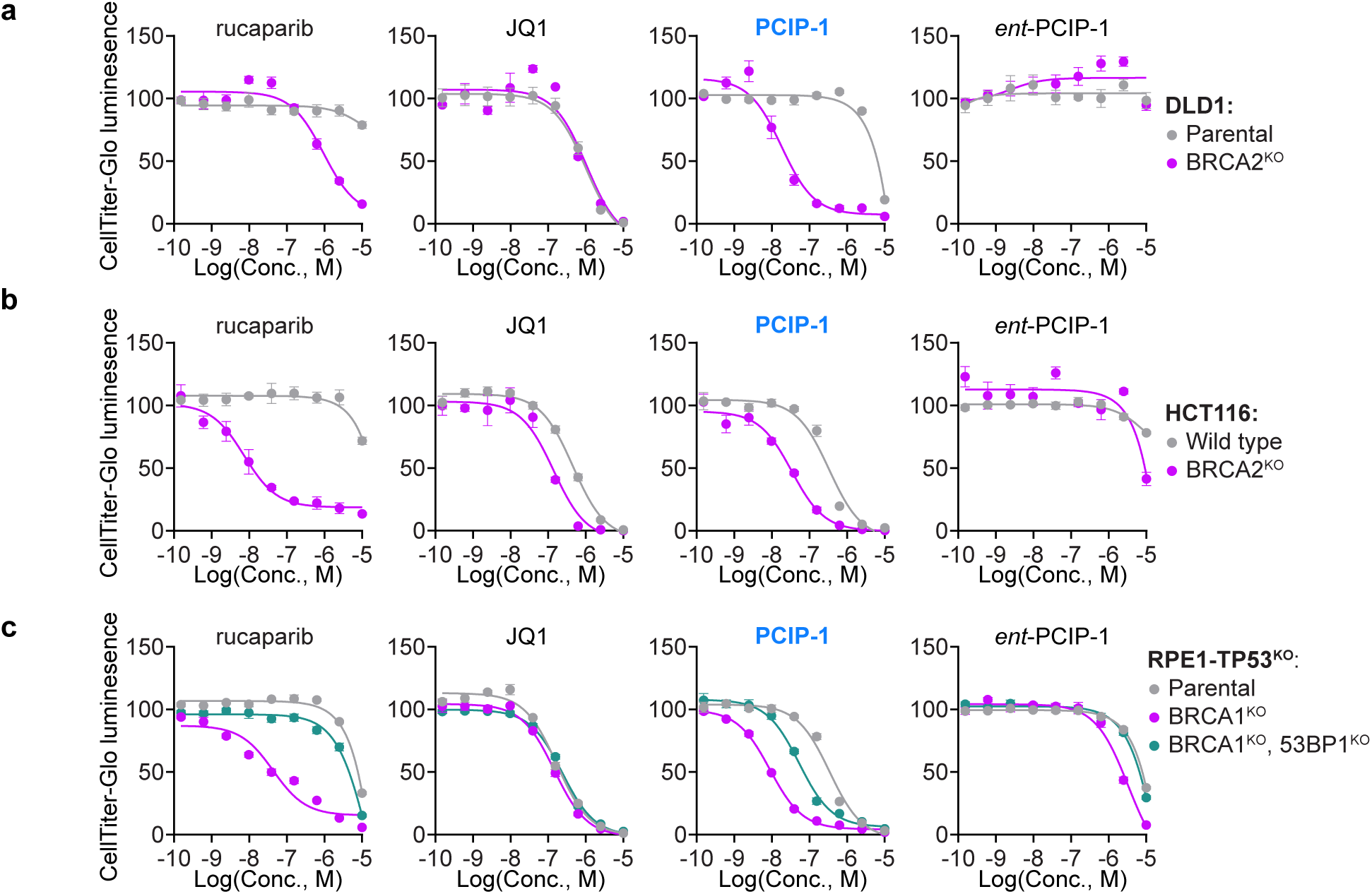
PCIP-1 is synthetically lethal in BRCA mutated cancers. **a**, DLD1 viability after 6 day compound treatment (CellTiter-Glo luminescence normalized to DMSO, *n* = 3, mean ± s.e.m.). **b,** HCT116 viability after 6 day compound treatment (CellTiter-Glo luminescence normalized to DMSO, *n* = 3, mean ± s.e.m.). **c,** RPE1-hTERT-TP53^KO^ viability after 6 day compound treatment (CellTiter-Glo luminescence normalized to DMSO, *n* = 3, mean ± s.e.m.).

## Discussion

The modular construction of heterobifunctional small molecules has enabled the rational discovery of CIPs that can rewire cellular interactions.^1,2^ TPD remains the most advanced application of proximity pharmacology, but it is now joined by a growing list of other CIPs.^5,31^ While proximity-induced changes in post-translational modifications (PTMs) can be relatively straightforward to measure, at least in principle, other potential outcomes of induced proximity can be comparatively difficult to anticipate. The number of unanticipated pharmacological mechanisms that have been revealed by serendipitously discovered CIPs—e.g. steric inhibition of enzyme activity by cyclophilin/immunophilin recruitment,^56–59^ induced degradation by polymerization of BCL6,^71,72^ and destruction of the nuclear pore complex by TRIM21 ligands^73–75^—highlights the exceptionally diverse cellular effects that can be achieved by induced proximity. It is therefore unsurprising that new classes of CIPs have been relatively difficult to discover, particularly those that do not directly regulate a measurable PTM.

Here, we discovered a new class of CIPs that inhibit the repair of DNA damage by recruiting BET proteins to PARP2. We have yet to determine how the recruitment of BET proteins to PARP2 causes a durable inhibition of DNA damage repair. However, given the exceptionally weak engagement of PARP2 by PCIP-1, we presume that it must trigger a biochemical reaction that is capable of multiple turnovers and is either slowly reversible or non-reversible. This could, for example, include the activation of aberrant transcriptional activity at sites of DNA damage or the PARylation of a protein brought into close proximity of PARP2. Alternatively, it might be possible for BET-associated proteins, like the kinase P-TEFb, to install post-translational modifications onto PARP2 or other proteins in its proximity. Indeed, the transient recruitment of BET proteins may lead to a durable trapping of PARP2 onto sites of DNA damage, which is known to contribute to the anti-proliferative effects of PARP1/2 inhibitors.^76^

To better understand these possibilities, we expect it will be important to determine which individual BET proteins (BRD2, BRD3, or BRD4) can mediate PCIP-1 activity, as well as which domains and functions within those proteins are essential. Likewise, it will be necessary to determine why PCIP-1 inhibits DNA damage repair through PARP2 and not PARP1. In total, our data suggests that PCIP-1 forms a productive ternary complex with PARP2 but not with PARP1. Since PARP1 is far more abundant than PARP2, this would explain the increased sensitivity in *PARP1*-knockout cells. Whether future PCIP analogs, including those constructed from structurally divergent PARP inhibitors, will continue to function through PARP2 is unclear. Understanding the determinants of PCIP-1 activity would greatly benefit future optimization. We wonder whether PARP-BET molecular glues— monovalent compounds that cooperatively stabilize protein-protein interactions—can be discovered in the future.

We have recently developed a high-throughput chemistry (HTC)-based approach to prospectively convert pre-existing ligands into molecular glue degraders, which may facilitate the analogous discovery of molecular glue PCIPs in the future.^77^

While PARP proteins have been successfully targeted by PROTAC degraders,^40,47^ DNA repair processes have not previously—to our knowledge—been rewired through the design of other heterobifunctional CIPs. In demonstrating synthetic lethality with HR deficiencies, PCIP-1 reflects favorably on the potential for this class of CIPs to be explored as a new therapeutic modality. The increased sensitivity of *PARP1* knockout cells to PCIP-1 suggests PCIPs might afford an attractive mechanism to overcome or circumvent a common form of PARPi drug resistance.^69,70^ Furthermore, these findings suggest that other DNA repair proteins should be explored for proximity pharmacology, potentially making further use, for example, of the agents being developed for Polθ in HR-deficient tumors or WRN helicase in MSI-high tumors.^64,66,78–87^ Ultimately, our findings highlight the underexplored potential of CIPs to modulate cellular biology in unanticipated ways, providing a generalizable framework to expand the scope of proximity pharmacology.

## Acknowledgements

This work was supported by the following funding sources the Baxter Foundation Young Investigator Award and the National Cancer Institute (R01CA280720) (M.A.E.); a postdoctoral fellowship from the National Center for Advancing Translational Science (T32TR004396) (B.D.C.); and a philanthropic gift, Dani Reiss Innovation Fund for Healthy Ageing (D.D.). The following Scripps Research facilities were instrumental in supporting this work: the Flow Cytometry Core Facility, the Automated Synthesis Facility, the Center for Metabolomics and Mass Spectrometry, and the Nuclear Magnetic Resonance Facility. We also gratefully acknowledge the MacRae lab for generously sharing instrumentation that enabled protein purification.

---

**Extended Data Fig. 1.**
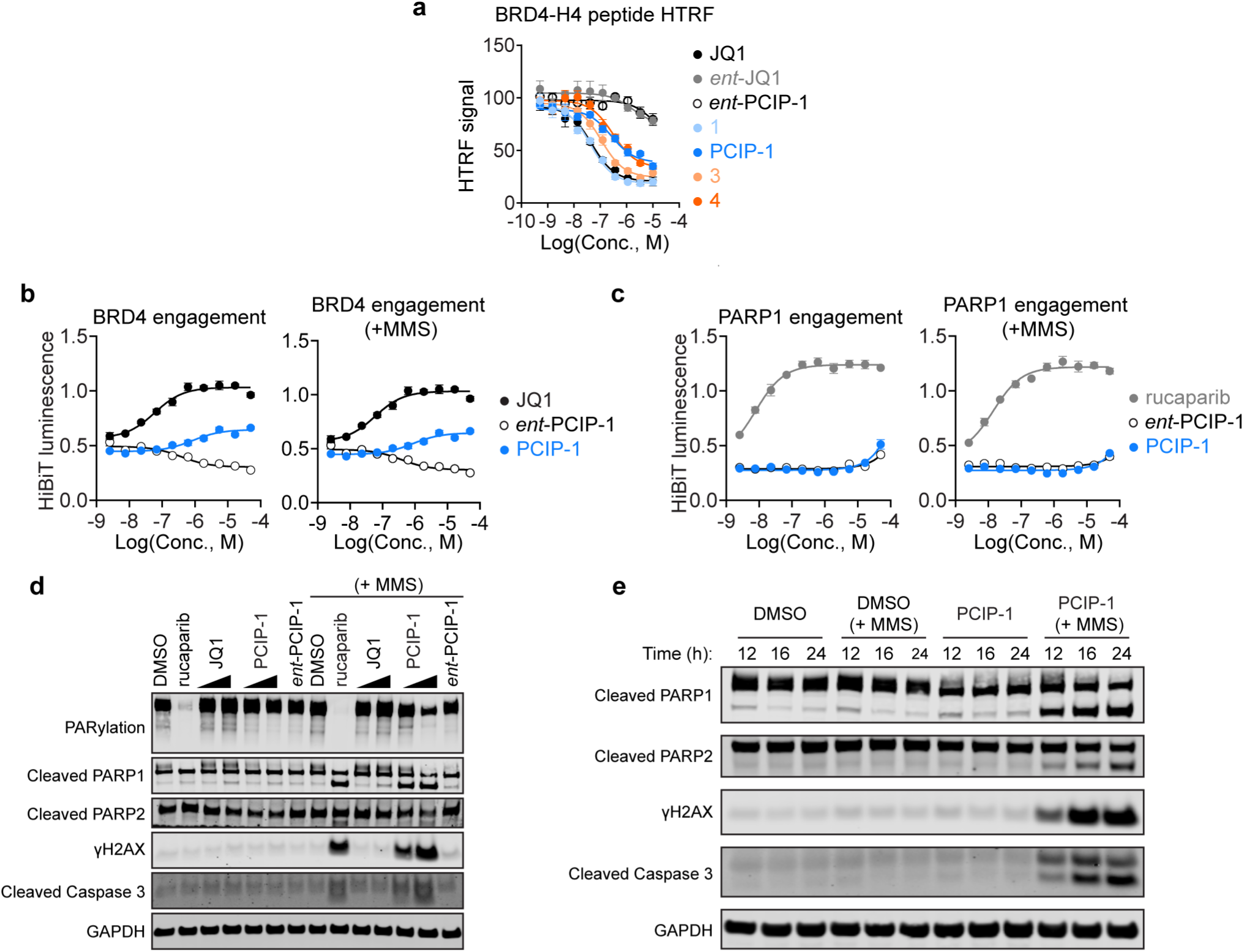
Characterization of PCIP candidates. **a,** Competitive BRD4–H4 peptide HTRF assay measuring inhibition of BRD4 association with an acetylated histone peptide (normalized by percent activity remaining) (*n* = 4, mean ± s.e.m.). **b,** BRD4 target engagement measured by displacement of dBET6 with and without 25 µM MMS (DMSO-normalized HiBiT luminescence, *n* = 3, mean ± s.e.m.). Data in Fig. 2b is taken from the same experiment (PCIP-1, *ent*-PCIP-1, and JQ1 are repeated in both panels). **c,** PARP1 target engagement measured by displacement of SK-575 with and without 25 µM MMS (DMSO-normalized HiBiT luminescence, *n* = 3, mean ± s.e.m.). **d,** Immunoblot following 24 h treatment of Jurkat cells. PCIP-1 and JQ1 were used at 100 nM and 1 µM. All other compounds were used at 1 µM, with and without 25 µM MMS . Data in Fig. 3b is taken from the same experiment (blots for γH2AX, PARylation, and GAPDH are repeated in both panels and stem from a singular experiment). **e,** Immunoblot following time-course treatment of Jurkat cells using 1 µM PCIP-1, with and without 25 µM MMS.

**Extended Data Fig. 2.**
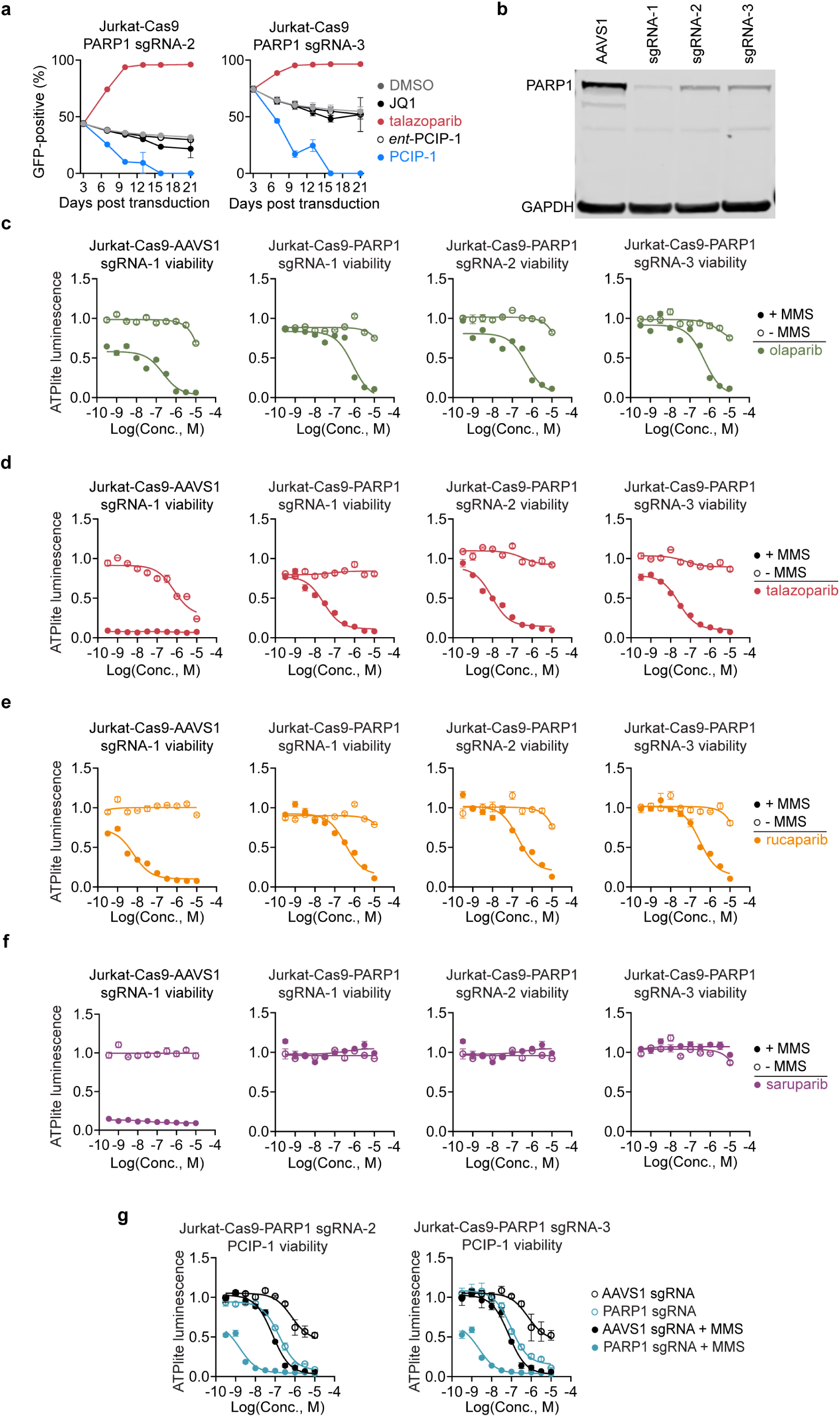
Effects of PARP1 depletion in Jurkat-Cas9 cells. **a**, Competitive growth assay testing the effect of *PARP1*-sgRNA-2-3 on the response of Jurkat-Cas9 cells to talazoparib (1 µM), PCIP-1 (1 µM), and *ent*-PCIP-1 (1 µM), or JQ1 (250 nM). Proportion of GFP-positive cells over time was measured by flow cytometry (*n* = 3, mean ± s.e.m.). **b,** Immunoblot analysis of Jurkat-Cas9 cells transduced with safe harbor sgRNA *AAVS1* and 2 sgRNAs targeting the *PARP1* gene, demonstrating successful depletion of *PARP1* function. **c,** Viability assay (72 h) in Jurkat-Cas9 cells with and without 25 µM MMS (ATPlite luminescence normalized to DMSO, *n* = 3, mean ± s.e.m.). **d,** same as **c** but using talazoparib in dose-response. **e,** same as **c** but using rucaparib in dose-response. **f,** same as **c** but using saruparib in dose-response. **g,** Viability assay (72 h) in Jurkat-Cas9 cells transduced with sgRNA-2,3 targeting *PARP1* gene, with and without 25 µM MMS (ATPlite luminescence normalized to DMSO, *n* = 3, mean ± s.e.m.).

**Extended Data Fig. 3.**
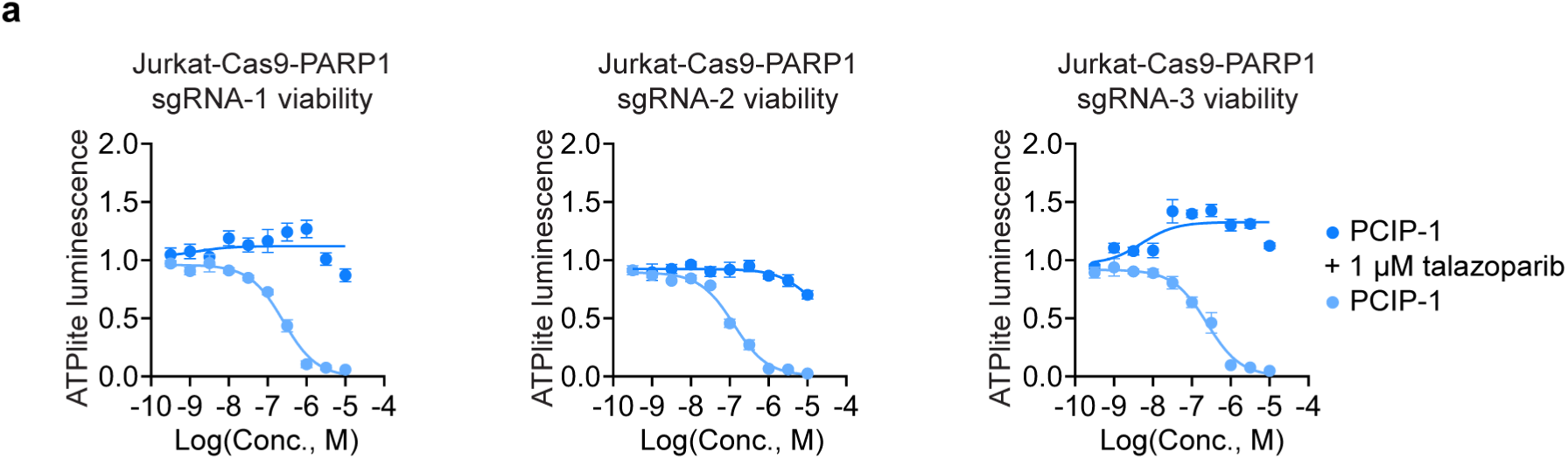
Viability effects of PCIP-1 in Jurkat-Cas9 cells transduced with safe harbor sgRNA *AAVS1* and sgRNAs targeting *PARP1*. **a**, Viability assay (72 h) in Jurkat-Cas9 cells transduced with sgRNA-1-3 targeting *PARP1* gene (ATPlite luminescence normalized to DMSO, *n* = 3, mean ± s.e.m.).

**Extended Data Fig. 4.**
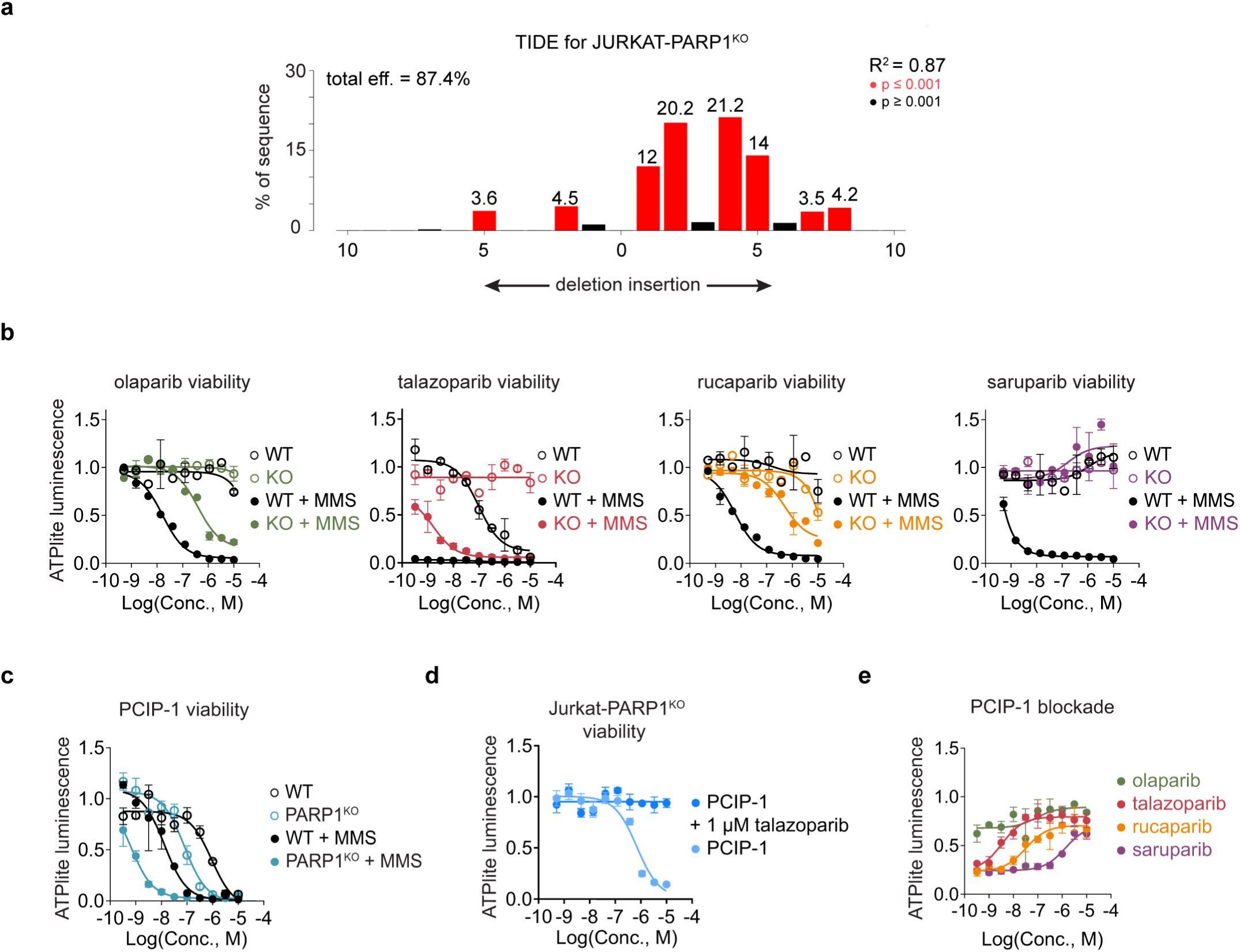
Validation of *PARP1* knockout effects. **a**, TIDE of CRISPR-Cas9 editing, targeting *PARP1* in Jurkat cells. Genomic data was collected after 2 weeks of selection with 1 µM talazoparib which began 7 days after transfection. Target locus was analyzed 500 bp flanking the cut site and was PCR-amplified and Sanger-sequenced. Indel distribution was analyzed with the TIDE webtool (https://tide.nki.nl/). Total editing efficiency was 87.4% (R² = 0.87). Red bars indicate indels with p < 0.001; black bars indicate non-significant indels (p ≥ 0.001). Most indels were centered around ±1–5 bp, consistent with efficient NHEJ repair. **b,** 72 h viability assay of Jurkat and Jurkat-PARP1^KO^ cells treated with indicated PARP1/2 inhibitors with and without 25 µM MMS (ATPlite luminescence normalized to DMSO, *n* = 2, mean ± s.e.m.). **c,** 72 h viability of Jurkat and Jurkat-PARP1^KO^ cells, with and without 25 µM MMS (ATPlite luminescence normalized to DMSO, *n* = 2, mean ± s.e.m.). **d,** 72 h viability assay of Jurkat-PARP1^KO^ (ATPlite luminescence normalized to DMSO, *n* = 5, mean ± s.e.m.). **e,** 72 h viability assay of Jurkat-PARP1^KO^ cells treated with 500 nM PCIP-1 (ATPlite luminescence normalized to DMSO, *n* = 2, mean ± s.e.m.).

**Extended Data Fig. 5.**
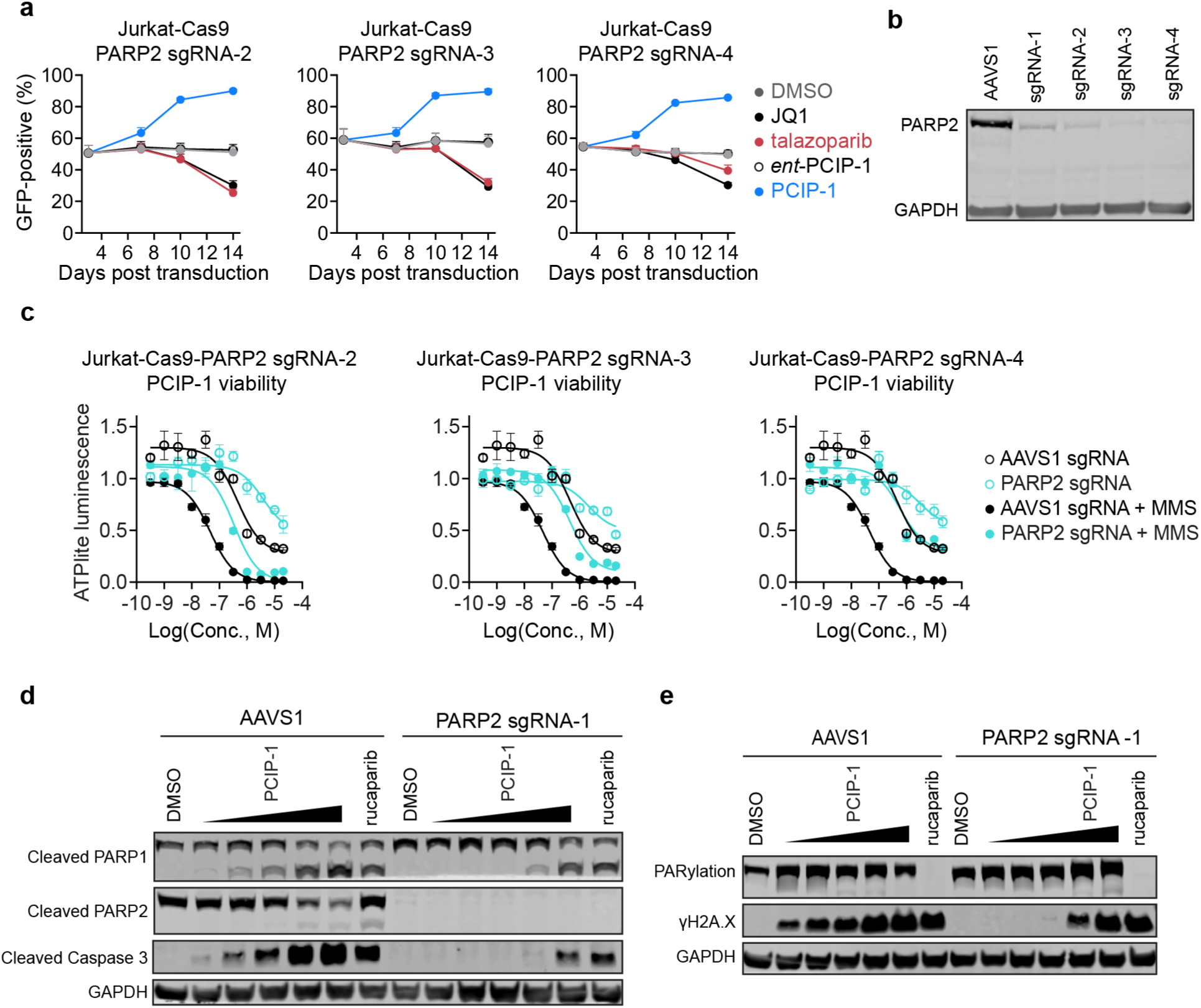
*PARP2* knockout confers resistance to PCIP-1 but not conventional PARP inhibitors. **a**, Competitive growth assay testing the effect of *PARP2*-sgRNA-2-4 on the response of Jurkat-Cas9 cells to 1 µM talazoparib, PCIP-1, *ent*-PCIP-1, and JQ1. Proportion of GFP-positive cells over time was measured by flow cytometry (*n* = 3, mean ± s.e.m.). **b,** Immunoblot analysis of Jurkat-Cas9 cells transduced with safe harbor sgRNA *AAVS1* and 4 sgRNA’s targeting the *PARP2* gene, demonstrating successful disruption of PARP2 function. **c,** Viability assay (72 h) in Jurkat-Cas9 cells with and without 25 µM MMS (ATPlite luminescence normalized to DMSO, *n* = 3, mean ± s.e.m.). **d,** Immunoblot analysis of Jurkat-Cas9 cells treated with PCIP-1 at 50 nM, 100 nM, 250 nM, 500 nM, and 1 µM, cells were transduced with safe harbor sgRNA *AAVS1* and sgRNA-1 targeting *PARP2* demonstrating rescue of apoptotic markers, co-treated with 25 µM MMS. **e,** Immunoblot analysis of Jurkat-Cas9 cells treated with PCIP-1 at 50 nM, 100 nM, 250 nM, 500 nM, and 1 µM, cells were transduced with safe harbor sgRNA *AAVS1* and sgRNA-1 targeting *PARP2* sgRNA 1 demonstrating DNA damage rescue, co-treated with 25 µM MMS.

**Extended Data Fig. 6.**
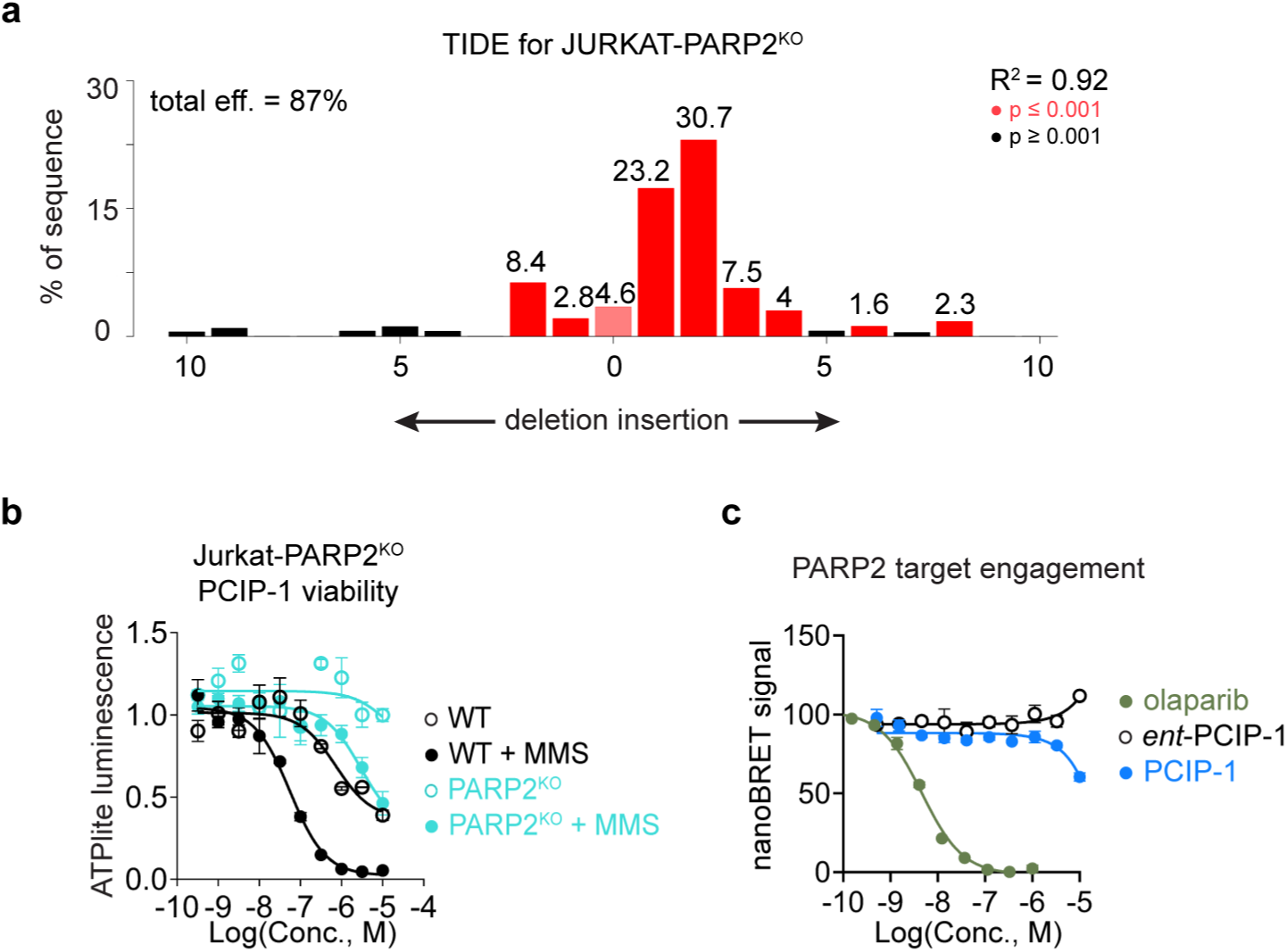
Validation of *PARP2* knockout effects. **a**, TIDE of CRISPR-Cas9 editing targeting *PARP2* in Jurkat cells. Genomic data was collected after 10 days of selection with 500 nM PCIP-1 which began 7 days after transfection. Target locus was analyzed 500 bp flanking the cut site and was PCR-amplified and Sanger-sequenced. Indel distribution was analyzed with the TIDE webtool (https://tide.nki.nl/). Total editing efficiency was 87% (R² = 0.92). Red bars indicate indels with p < 0.001; black bars indicate non-significant indels (p ≥ 0.001). Most indels were centered around ±1–5 bp, consistent with efficient NHEJ repair. **b,** 72 h viability of Jurkat and Jurkat-PARP2^KO^ cells, with and without 25 µM MMS (ATPlite luminescence normalized to DMSO, *n* = 3, mean ± s.e.m.). **c,** NanoBRET intracellular target engagement *PARP2* assay performed by Reaction Biology (fluorescent signal normalized to DMSO, *n* = 2, mean ± s.e.m.).

## References

1. Stanton, B. Z., Chory, E. J. & Crabtree, G. R. Chemically induced proximity in biology and medicine. Science 359, (2018).

2. Gerry, C. J. & Schreiber, S. L. Unifying principles of bifunctional, proximity-inducing small molecules. Nat. Chem. Biol. 16, 369–378 (2020).

3. Bondeson, D. P. et al. Catalytic in vivo protein knockdown by small-molecule PROTACs. Nat. Chem. Biol. 11, 611–617 (2015).

4. Lai, A. C. & Crews, C. M. Induced protein degradation: an emerging drug discovery paradigm. Nat. Rev. Drug Discov. 16, 101–114 (2016).

5. Li, K. & Crews, C. M. PROTACs: past, present and future. Chem. Soc. Rev. 51, 5214–5236 (2022).

6. Spencer, D. M., Wandless, T. J., Schreiber, S. L. & Crabtree, G. R. Controlling signal transduction with synthetic ligands. Science 262, 1019–1024 (1993).

7. Belshaw, P. J., Ho, S. N., Crabtree, G. R. & Schreiber, S. L. Controlling protein association and subcellular localization with a synthetic ligand that induces heterodimerization of proteins. Proc. Natl. Acad. Sci. U. S. A. 93, 4604–4607 (1996).

8. Ho, S. N., Biggar, S. R., Spencer, D. M., Schreiber, S. L. & Crabtree, G. R. Dimeric ligands define a role for transcriptional activation domains in reinitiation. Nature 382, 822–826 (1996).

9. Rivera, V. M. et al. A humanized system for pharmacologic control of gene expression. Nat. Med. 2, 1028– 1032 (1996).

10. Amara, J. F. et al. A versatile synthetic dimerizer for the regulation of protein-protein interactions. Proc. Natl. Acad. Sci. U. S. A. 94, 10618–10623 (1997).

11. Clackson, T. et al. Redesigning an FKBP-ligand interface to generate chemical dimerizers with novel specificity. Proc. Natl. Acad. Sci. U. S. A. 95, 10437–10442 (1998).

12. Henning, N. J. et al. Deubiquitinase-targeting chimeras for targeted protein stabilization. Nat. Chem. Biol. 18, 412–421 (2022).

13. Yamazoe, S. et al. Heterobifunctional molecules induce dephosphorylation of kinases-A proof of concept study. J. Med. Chem. 63, 2807–2813 (2020).

14. Siriwardena, S. U. et al. Phosphorylation-inducing chimeric small molecules. J. Am. Chem. Soc. 142, 14052–14057 (2020).

15. Chen, P.-H. et al. Modulation of phosphoprotein activity by phosphorylation targeting chimeras (PhosTACs). ACS Chem. Biol. 16, 2808–2815 (2021).

16. Wang, W. W. et al. Targeted protein acetylation in cells using heterobifunctional molecules. J. Am. Chem. Soc. 143, 16700–16708 (2021).

17. Kabir, M. et al. Acetylation Targeting Chimera enables acetylation of the tumor suppressor p53. J. Am. Chem. Soc. 145, 14932–14944 (2023).

18. Ramirez, D. H. et al. Engineering a proximity-directed O-GlcNAc transferase for selective protein O-GlcNAcylation in cells. ACS Chem. Biol. 15, 1059–1066 (2020).

19. Ge, Y. et al. Target protein deglycosylation in living cells by a nanobody-fused split O-GlcNAcase. Nat. Chem. Biol. 17, 593–600 (2021).

20. Ma, B. et al. Targeted protein O-GlcNAcylation using bifunctional small molecules. J. Am. Chem. Soc. 146, 9779–9789 (2024).

21. Seabrook, L. J. et al. Methylarginine targeting chimeras for lysosomal degradation of intracellular proteins. Nat. Chem. Biol. 20, 1566–1576 (2024).

22. Banik, S. M. et al. Lysosome-targeting chimaeras for degradation of extracellular proteins. Nature 584, 291–297 (2020).

23. Huang, B. et al. Designed endocytosis-inducing proteins degrade targets and amplify signals. Nature 638, 796–804 (2025).

24. Zhang, D. et al. Transferrin receptor targeting chimeras for membrane protein degradation. Nature 638, 787–795 (2025).

25. Tong, Y. et al. Programming inactive RNA-binding small molecules into bioactive degraders. Nature 618, 169–179 (2023).

26. Costales, M. G., Matsumoto, Y., Velagapudi, S. P. & Disney, M. D. Small molecule targeted recruitment of a nuclease to RNA. J. Am. Chem. Soc. 140, 6741–6744 (2018).

27. Erwin, G. S. et al. Synthetic transcription elongation factors license transcription across repressive chromatin. Science 358, 1617–1622 (2017).

28. Gourisankar, S. et al. Rewiring cancer drivers to activate apoptosis. Nature 620, 417–425 (2023).

29. Gibson, W. J. et al. Bifunctional small molecules that induce nuclear localization and targeted transcriptional regulation. J. Am. Chem. Soc. 145, 26028–26037 (2023).

30. Ng, C. S. C., Liu, A., Cui, B. & Banik, S. M. Targeted protein relocalization via protein transport coupling. Nature 633, 941–951 (2024).

31. Nalawansha, D. A., Mangano, K., den Besten, W. & Potts, P. R. TAC-tics for leveraging proximity biology in drug discovery. Chembiochem 25, e202300712 (2024).

32. Farmer, H. et al. Targeting the DNA repair defect in BRCA mutant cells as a therapeutic strategy. Nature 434, 917–921 (2005).

33. Bryant, H. E. et al. Specific killing of BRCA2-deficient tumours with inhibitors of poly(ADP-ribose) polymerase. Nature 434, 913–917 (2005).

34. Ray Chaudhuri, A. & Nussenzweig, A. The multifaceted roles of PARP1 in DNA repair and chromatin remodelling. Nat. Rev. Mol. Cell Biol. 18, 610–621 (2017).

35. Fong, P. C. et al. Inhibition of poly(ADP-ribose) polymerase in tumors from BRCA mutation carriers. N. Engl. J. Med. 361, 123–134 (2009).

36. Filippakopoulos, P. et al. Selective inhibition of BET bromodomains. Nature 468, 1067–1073 (2010).

37. Canan Koch, S. S., et al. Novel tricyclic poly(ADP-ribose) polymerase-1 inhibitors with potent anticancer chemopotentiating activity: design, synthesis, and X-ray cocrystal structure. J. Med. Chem. 45, 4961–4974 (2002).

38. Anders, L. et al. Genome-wide localization of small molecules. Nat. Biotechnol. 32, 92–96 (2014).

39. Winter, G. E. et al. Phthalimide conjugation as a strategy for in vivo target protein degradation. Science 348, 1376–1381 (2015).

40. Wang, S. et al. Uncoupling of PARP1 trapping and inhibition using selective PARP1 degradation. Nat. Chem. Biol. 15, 1223–1231 (2019).

41. Liszczak, G. P. et al. Genomic targeting of epigenetic probes using a chemically tailored Cas9 system. Proc. Natl. Acad. Sci. U. S. A. 114, 681–686 (2017).

42. Chiarella, A. M. et al. Dose-dependent activation of gene expression is achieved using CRISPR and small molecules that recruit endogenous chromatin machinery. Nat. Biotechnol. 38, 50–55 (2020).

43. Donovan, K. A. et al. Mapping the degradable kinome provides a resource for expedited degrader development. Cell 183, 1714–1731.e10 (2020).

44. Gechijian, L. N. et al. Functional TRIM24 degrader via conjugation of ineffectual bromodomain and VHL ligands. Nat. Chem. Biol. 14, 405–412 (2018).

45. Riching, K. M. et al. Quantitative Live-Cell Kinetic Degradation and Mechanistic Profiling of PROTAC Mode of Action. ACS Chem. Biol. 13, 2758–2770 (2018).

46. Schwinn, M. K. et al. CRISPR-mediated tagging of endogenous proteins with a luminescent peptide. ACS Chem. Biol. 13, 467–474 (2018).

47. Cao, C. et al. Discovery of SK-575 as a Highly Potent and Efficacious Proteolysis-Targeting Chimera Degrader of PARP1 for Treating Cancers. J. Med. Chem. 63, 11012–11033 (2020).

48. Winter, G. E. et al. BET bromodomain proteins function as master transcription elongation factors independent of CDK9 recruitment. Mol. Cell 67, 5–18.e19 (2017).

49. Douglass, E. F., Jr, Miller, C. J., Sparer, G., Shapiro, H. & Spiegel, D. A. A comprehensive mathematical model for three-body binding equilibria. J. Am. Chem. Soc. 135, 6092–6099 (2013).

50. Murai, J. et al. Trapping of PARP1 and PARP2 by Clinical PARP Inhibitors. Cancer Res. 72, 5588–5599 (2012).

51. Mah, L.-J., El-Osta, A. & Karagiannis, T. C. γH2AX: a sensitive molecular marker of DNA damage and repair. Leukemia 24, 679–686 (2010).

52. Murai, J. et al. Stereospecific PARP trapping by BMN 673 and comparison with olaparib and rucaparib. Mol. Cancer Ther. 13, 433–443 (2014).

53. Shi, J. et al. Discovery of cancer drug targets by CRISPR-Cas9 screening of protein domains. Nat. Biotechnol. 33, 661–667 (2015).

54. Johannes, J. W. et al. Discovery of 5-{4-[(7-Ethyl-6-oxo-5,6-dihydro-1,5-naphthyridin-3-yl)methyl]piperazin-1-yl}-N-methylpyridine-2-carboxamide (AZD5305): A PARP1–DNA Trapper with High Selectivity for PARP1 over PARP2 and Other PARPs. J. Med. Chem. 64, 14498–14512 (2021).

55. Brinkman, E. K., Chen, T., Amendola, M. & van Steensel, B. Easy quantitative assessment of genome editing by sequence trace decomposition. Nucleic Acids Res. 42, e168 (2014).

56. Schreiber, S. L. & Crabtree, G. R. The mechanism of action of cyclosporin A and FK506. Immunol. Today 13, 136–142 (1992).

57. Yang, H. et al. mTOR kinase structure, mechanism and regulation. Nature 497, 217–223 (2013).

58. Sabatini, D. M., Erdjument-Bromage, H., Lui, M., Tempst, P. & Snyder, S. H. RAFT1: a mammalian protein that binds to FKBP12 in a rapamycin-dependent fashion and is homologous to yeast TORs. Cell 78, 35– 43 (1994).

59. Sabers, C. J. et al. Isolation of a protein target of the FKBP12-rapamycin complex in mammalian cells. J. Biol. Chem. 270, 815–822 (1995).

60. Schulze, C. J. et al. Chemical remodeling of a cellular chaperone to target the active state of mutant KRAS. Science 381, 794–799 (2023).

61. Noordermeer, S. M. et al. The shieldin complex mediates 53BP1-dependent DNA repair. Nature 560, 117– 121 (2018).

62. Zimmermann, M. et al. CRISPR screens identify genomic ribonucleotides as a source of PARP-trapping lesions. Nature 559, 285–289 (2018).

63. Hewitt, G. et al. Defective ALC1 nucleosome remodeling confers PARPi sensitization and synthetic lethality with HRD. Mol. Cell 81, 767–783.e11 (2021).

64. Zatreanu, D. et al. Polθ inhibitors elicit BRCA-gene synthetic lethality and target PARP inhibitor resistance. Nat. Commun. 12, 3636 (2021).

65. Illuzzi, G. et al. Preclinical characterization of AZD5305, A next-generation, highly selective PARP1 inhibitor and trapper. *Clin. Cancer Res.* **28**, 4724–4736 (2022).

66. Fried, W. et al. Discovery of a small-molecule inhibitor that traps Polθ on DNA and synergizes with PARP inhibitors. Nat. Commun. 15, 2862 (2024).

67. Bouwman, P. et al. 53BP1 loss rescues BRCA1 deficiency and is associated with triple-negative and BRCA-mutated breast cancers. Nature structural & molecular biology vol. 17 688–695 (2010).

68. Bunting, S. F. et al. 53BP1 inhibits homologous recombination in Brca1-deficient cells by blocking resection of DNA breaks. Cell vol. 141 243–254 (2010).

69. Gogola, E., Rottenberg, S. & Jonkers, J. Resistance to PARP inhibitors: Lessons from preclinical models of BRCA-associated cancer. Annu. Rev. Cancer Biol. 3, 235–254 (2019).

70. Noordermeer, S. M. & van Attikum, H. PARP inhibitor resistance: A tug-of-war in BRCA-mutated cells. Trends Cell Biol. 29, 820–834 (2019).

71. Kerres, N. et al. Chemically induced degradation of the oncogenic transcription factor BCL6. Cell Rep. 20, 2860–2875 (2017).

72. Słabicki, M. et al. Small-molecule-induced polymerization triggers degradation of BCL6. Nature 588, 164– 168 (2020).

73. Lu, P. et al. Selective degradation of multimeric proteins by TRIM21-based molecular glue and PROTAC degraders. Cell 187, 7126–7142.e20 (2024).

74. Li, X. et al. Chemically induced Nuclear Pore Complex Protein degradation via TRIM21. ACS Chem. Biol. 20, 1020–1028 (2025).

75. Cheng, Y. et al. TRIM21-NUP98 interface accommodates structurally diverse molecular glue degraders. ACS Chem. Biol. 20, 953–959 (2025).

76. Blessing, C. et al. The oncogenic helicase ALC1 regulates PARP inhibitor potency by trapping PARP2 at DNA breaks. Mol. Cell 80, 862–875.e6 (2020).

77. Shaum, J. B. et al. High-throughput diversification of protein-ligand surfaces to discover chemical inducers of proximity. bioRxiv 2024.09.30.615685 (2024) doi:10.1101/2024.09.30.615685.

78. Chan, E. M. et al. WRN helicase is a synthetic lethal target in microsatellite unstable cancers. Nature 568, 551–556 (2019).

79. Kategaya, L., Perumal, S. K., Hager, J. H. & Belmont, L. D. Werner syndrome helicase is required for the survival of cancer cells with microsatellite instability. iScience 13, 488–497 (2019).

80. van Wietmarschen, N. et al. Repeat expansions confer WRN dependence in microsatellite-unstable cancers. Nature 586, 292–298 (2020).

81. Ferretti, S. et al. Discovery of WRN inhibitor HRO761 with synthetic lethality in MSI cancers. Nature 629, 443–449 (2024).

82. Rodríguez Pérez, F., et al. WRN inhibition leads to its chromatin-associated degradation via the PIAS4-RNF4-p97/VCP axis. Nat. Commun. 15, 6059 (2024).

83. Baltgalvis, K. A. et al. Chemoproteomic discovery of a covalent allosteric inhibitor of WRN helicase. Nature 629, 435–442 (2024).

84. Ito, F. et al. Structural basis for a Polθ helicase small-molecule inhibitor revealed by cryo-EM. Nat. Commun. 15, 7003 (2024).

85. Mateos-Gomez, P. A. et al. Mammalian polymerase θ promotes alternative NHEJ and suppresses recombination. Nature 518, 254–257 (2015).

86. Ceccaldi, R. et al. Homologous-recombination-deficient tumours are dependent on Polθ-mediated repair. Nature 518, 258–262 (2015).

87. Bubenik, M. et al. Identification of RP-6685, an orally bioavailable compound that inhibits the DNA polymerase activity of Polθ. J. Med. Chem. 65, 13198–13215 (2022).

