## Supplementary material for "Rewiring DNA repair with PARP-based chemical inducers of proximity": Methods and Supporting Information

### General information, materials, and methods

**Analytical data**: $^{1}H$, $^{13}C$, and $^{19}F$ spectra were recorded on the following NMR spectrometers: Bruker AV NEO 400 MHz equipped with a 5 mm BBFO probe, Bruker AVIII HD 600 MHz equipped with a 5 mm CPQCI and 1.7 mm CPTCI CryoProbe, Bruker AVIII HD 600 MHz MNR equipped with a 5 mm CPDCH cryoprobe, or JEOL JNM-ECZ400R 400 MHz equipped with a 5 mm H/F/X royal probe. The spectrometers were automatically tuned and matched to the correct operating frequencies. Proton ($^{1}H$), carbon ($^{13}C$), and fluorine ($^{19}F$) chemical shifts are reported in parts per million ($\delta$) with respect to tetramethylsilane (TMS, $\delta$ = 0) and referenced internally with respect to the protio solvent impurity or hexafluorobenzene (HFB) for $^{19}F$ spectra (HFB, $\delta$ = -164.9). Multiplicities are abbreviated: singlet, s; doublet, d; triplet, t; quartet, q; doublet of doublet, dd; doublet of doublet of doublets, ddd; multiplet, m. Deuterated NMR solvents were obtained from Cambridge Isotope Laboratories, Inc., Andover, MA, and used without purification. Spectra were digitally processed (phase and baseline corrections, integration, peak analysis) using MestreNova. Unless otherwise noted all spectra were obtained at 298 K and all $^{13}C$ spectra are {$^{1}H$}.

**Ultra performance liquid chromatography**: Ultra performance liquid chromatography (UPLC) data was collected using a Waters I-Class LC with diode array and QDa mass spectrometer.

**High-resolution mass spectrometry:** HRMS (ESI-TOF) was run as direct injection and analyzed on an Agilent 6230 TOF LC/MS System.

**Purification chromatography:** Normal and reverse phase chromatography were run using Teledyne ISCO Combiflash NextGen 300+ auto columns. Column size and conditions are compound specific and are listed as such. Unless otherwise explicitly stated, compounds purified *via* reverse phase chromatography are isolated as the TFA salt.

**Reagents and solvents:** Reagents and solvents were purchased from commercial sources and used as received. All solvents were purchased as anhydrous Sure/Seal™.

**Reactions**: Reactions were performed as written and monitored by ultra-performance liquid chromatography (UPLC).

### Genetic modification of cell lines

**Jurkat-BRD4-HiBit and Jurkat-PARP1-HiBit cell line creation**: All components necessary were ordered from Integrated DNA Technologies. A Neon Transfection System (ThermoFisher) was used to electroporate cells employing the 1 μL tip kit. Briefly, sgRNA complexes are synthesized by mixing equal volumes of crRNA (2 μL, 160 μM in nuclease free water, aatcttttctgagcgcacct for BRD4-HiBit and taagacctccctgtggtaat for PARP1-HiBit) and tracrRNA (2 μL, 160 μM) and are incubated at 37 °C for 30 min before Cas9 (2.7 μL, 10 μg/μL, Alt-R™ S.p. Cas9 Nuclease V3, catalog number 1081058) and duplex buffer (1.3 μL, 30 mM HEPES, pH 7.5; 100 mM potassium acetate in nuclease free water) are added. This suspension is incubated at 37 °C for an additional 15 min to form the active RNP complex. The RNP solution (1 μL) is added to ssODN (2 μM in nuclease free water, catgaatttccagagtgatctattgtcaatatttgaagaaaatcttttcggtggcggtggctcgggcggtggtgggtcgggtggcggcggatctgt gagcggctggcggctgttcaagaagattagctgacctaggtggcttctgactttgattttctggcaaaacattgactttccata for BRD4-HiBit and atctgaagtatctgctgaaactgaaattcaattttaagacctccctgtggggaagcggagtaagcggctggcggctgttcaagaagattagcta aaattgggagaggtagccgagtcacacccggtggctctggtatgaattcac for PARP1-HiBit). The resulting mixture is electroporated (1700 pulse voltage, 20 millisecond, and 1 pulse for Jurkats) containing 200,000 cells in electroporation buffer before the cells are deposited into antibiotic free media. The cells are allowed to rest in a 37 °C incubator for 24 h before additional antibiotic containing media is added. Replicates with the highest overall signal were selected and expanded.

**22Rv1FKBP12$^{F36V}$-2xHA-BRD4**: BRD4 (CCDS 12328.1 from NCBI) was inserted into the gateway compatible lentiviral destination plasmid pLEX_305-N-dTAG 22Rv1 (Addgene #91797) using gateway cloning. The final lentiviral FKBP12$^{F36V}$-2xHA-BRD4 plasmid was confirmed by whole plasmid sequencing. 22Rv1 cells were purchased from ATCC (CRL-2505). Lenti-X 293T (DMEM supplemented with 10% FBS and Gibco Antibiotic-Antimycotic) cells were purchased from Takara for lentivirus production. All cell lines were tested negative for mycoplasma infections regularly. Lentiviral packaging plasmids pMD2.G (a gift from Didier Trono, Addgene plasmid #12259; http://n2t.net/addgene:12259; RRID:Addgene_12259), psPAX2 (a gift from Didier Trono, Addgene plasmid #12260; http://n2t.net/addgene:12260; RRID:Addgene_12260), and the lentiviral expression plasmid were co-transfected to Lenti-X 293T cells to produce corresponding lentivirus. Supernatants with viral particles were harvested at 48 and 72 hours after transfection, filtered with 0.44 μm membrane and concentrated by 50-fold with Lenti-X Concentrator (Takara, #631232). All cells were transduced by spinoculation at 800 g for 1 hour at room temperature supplemented with 8 μg/mL polybrene. After at least 72 hours post transduction, cells were selected with 2 μg/mL puromycin for at least 1 week, and expression was confirmed by immunoblot.

**Jurkat-PARP1$^{KO}$ and PARP2$^{KO}$ *via* Transfection and TIDE Analysis:** PARP1$^{KO}$ cells and PARP2$^{KO}$ cells were generated *via* electroporation using the Neon Electroporation System protocol described in "**Jurkat-BRD4-HiBit and Jurkat-PARP1-HiBit Cell Line Creation"** by omitting the procedural step of mixing RNP and ssODN. Instead, RNP is directly mixed into cells suspended in electroporation buffer before electroporation. The sgRNA sequences used were as follows: PARP1 (cgatgcctattactgcactg) and PARP2 (cgatgcctattactgcactg). PARP1$^{KO}$ cells were selected *via* continual treatment of 1 μM talazoparib over 14 days, beginning 7 days after transfection. PARP2$^{KO}$ cells were selected *via* continual treatment of 500 nM PCIP-1 (compound **2**) for 10 days, beginning 7 days after transfection. Following selection, gDNA was extracted using the DNeasy Blood and Tissue Kit (Qiagen, Cat # 69504). Amplicons were generated for sequencing using CloneAmp HiFi PCR (Takara, Cat. # 639298) and custom DNA oligos were designed for PARP1 (IDT, 5'-cttgctctagagtgccagg-3', 5'-ggaggtattttgcgttgagaat-3') and PARP2 (IDT, 5'-gccccacttggtaggacttc-3', 5'-tttctaggtcacggggctct-3'). PCR purification was done using the QIAquick PCR Purification Kit (Qiagen, Cat # 28104). Purified product was then sent for Sanger sequencing *via* GeneWiz. Knockout percentage was measured by tracking of indels by decomposition (TIDE) sequencing and visualized online (https://tide.nki.nl/.).

**Jurkat-Cas9:** Lenti X cells were co-transfected with BFP Cas9 plasmid pFUGW Cas9-BFP (Addgene #127396) and packaging plasmids psPAX2 and pMD2.G. Lentivirus was collected and concentrated with Lenti X concentrator (Takara). Jurkat cells at passage number 13 were transduced with 1X BFP Cas9 Lentivirus. Cells were sorted into three BFP expression level populations. The highest BFP expression level population was expanded for transduction with Brunello guide sgRNA lentivirus for future gene knockout studies.

**Jurkat-Cas9-PARP1$^{KO}$:** Jurkat-Ca9 cells were transduced with PARP1 targeting sgRNAs-1-3: cgatgcctattactgcactg, taccgatcaccgtacccaca, and agctaggcatgattgaccgc that were cloned into an LRG lentiviral vector with EGFP tags. Lenti X cells were co-transfected with PARP1 sgRNA plasmids and packaging plasmids psPAX2 and pMD2.G. Lentivirus was collected and concentrated with Lenti X concentrator (Takara).

**Jurkat-Cas9-PARP2$^{KO}$:** Jurkat-Ca9 cells were transduced with PARP2 targeting sgRNAs-1-4: catgcaatgaattctacacc, aataccaagaaagccccact, gggggcgcaaggcacaatgt, and ttgttcaggcaatctcaaca that were cloned into an LRG lentiviral vector with EGFP tags. Lenti X cells were co-transfected with PARP1 sgRNA plasmids and packaging plasmids psPAX2 and pMD2.G. Lentivirus was collected and concentrated with Lenti X concentrator (Takara).

**Cellular or biochemical assays**

**Graphing and $EC_{50}$ and $IC_{50}$ calculations:** All assay graphs were plotted using Graphpad Prism 10 and fitted using Log(inhibitor) vs. response (three parameters) which automatically calculated $EC_{50}$ and $IC_{50}$ value. For datasets where drug treatments did not result in a complete response magnitude the graphs were modelled using Log(inhibitor) vs. response Variable slope (4 parameters), where the data was constrained to the lowest and highest values found in the dataset, to ensure accurate calculation of $EC_{50}$ and $IC_{50}$ values.

**BRD4 target engagement assay:** Cells were brought to 1,000,000 cells/mL, and 18,000 cells (18 µL) were seeded into white 384-well polystyrene microplates (Greiner Bio-One) that had been seeded with indicated drugs by transferring from a source plate using an 655 Echo Acoustic Liquid Handler Instrument (Beckman). A 0.5 µM working solution of dBET6 was prepared by diluting 4 µL of a 10 mM DMSO stock into 8 mL of RPMI-1640 medium supplemented with 10% heat-inactivated fetal bovine serum (FBS) and 5% Antibiotic-Antimycotic (Thermo Fisher, #15240062). A corresponding DMSO-only solution in supplemented media was prepared in parallel. After 2 hours of compound incubation, 2 µL of the 0.5 µM dBET6 solution was added to wells that were pre-seeded with compound, while 2 µL of DMSO-only media was added to wells pre-seeded with DMSO. After a further 45-minute incubation at 37 °C, 20 µL of Nano-Glo® HiBiT Lytic Detection Reagent (Promega, N3050) was added to each well. Plates were centrifuged at 200 × g for 1 minute, incubated for 5 minutes at room temperature, and luminescence was measured using a Clariostar Plus microplate reader (BMG Labtech). Luminescence values from dBET6-treated wells were normalized to DMSO-only controls to visualize the rescue of the degradation effect. The assay was performed with three technical replicates per condition. Data points are presented as the mean of these replicates, and error bars represent the standard error of the mean (s.e.m.).

**PARP1 target engagement assay:** Cells were brought to 1,000,000 cells/mL, and 18,000 cells (18 µL) were seeded into white 384-well polystyrene microplates (Greiner Bio-One) that had been seeded with indicated drugs by transferring from a source plate using an 655 Echo Acoustic Liquid Handler Instrument (Beckman). A 0.23 µM working solution of SK-575 was prepared by diluting 2 µL of a 10 mM DMSO stock into 8 mL of RPMI-1640 medium supplemented with 10% heat-inactivated fetal bovine serum (FBS) and 5% Antibiotic-Antimycotic (Thermo Fisher, #15240062). A corresponding DMSO-only solution in supplemented media was prepared in parallel. After 2 hours of compound incubation, 2 µL of the 0.5 µM SK-575 solution was added to wells that were pre-seeded with compound, while 2 µL of DMSO-only media was added to wells pre-seeded with DMSO. After a further 4 hours incubation at 37 °C, 20 µL of Nano-Glo® HiBiT Lytic Detection Reagent (Promega, N3050) was added to each well. Plates were centrifuged at 200 × g for 1 minute, incubated for 5 minutes at room temperature, and luminescence was measured using a Clariostar Plus microplate reader (BMG Labtech). Luminescence values from SK-575-treated wells were normalized to DMSO-only controls to visualize the rescue of the degradation effect.The assay was performed with three technical replicates per condition. Data points are presented as the mean of these replicates, and error bars represent the standard error of the mean (s.e.m.).

**Viability assay for Jurkat, Jurkat-PARP1$^{KO}$ and Jurkat-PARP2$^{KO}$ cells:** Cells were diluted to a concentration of 20,000 cells/mL, and 1,000 cells (50 µL) were seeded into white 384-well polystyrene microplates (Greiner Bio-One). Compounds were pre-deposited into the plates using an 655 Echo acoustic liquid handler (Beckman) to transfer from a source plate. Viability studies involving methyl methanesulfonate (MMS) were performed by pretreating cells with MMS from a 25 mM stock solution prepared in RPMI-1640 supplemented media. Cells in 50 µL of culture media at 20,000 cells/mL were seeded into the compound-containing plates and incubated for 72 hours at 37 °C. Cell viability was assessed using ATPlite luminescence detection reagent (Revvity), which was diluted 1:4 using MilliQ water. Following incubation, 25 µL of ATPlite reagent, diluted 1:4 with MilliQ water, was added to wells. Plates were centrifuged at 200 × g for 1 minute, incubated at room temperature for 5 minutes, and luminescence was measured using a Clariostar Plus microplate reader (BMG Labtech). In all cases, the mean of technical replicates was plotted, with error bars representing the standard error of the mean (s.e.m.). Values were determined by normalizing treated wells to DMSO wells.

**Jurkat-Cas9 PARP1 and PARP2 Knockout Competitive Growth Assays:** Jurkat cells expressing Cas9 with a BFP reporter tag were sorted and high expression level populations were collected. Three Brunello sgRNA guides targeting PARP1 and four sgRNA guides targeting PARP2 were cloned into lentiviral expression vectors and Lentivirus was prepared and collected as previously described (Takara). JURKAT Cas9 cells were transduced with a titration of virus containing PARP1 or PARP2 guides, with AAVS1 sgRNA as a negative control, and RPS19 sgRNA as a positive control to verify Cas9 activity. On day 3 post transduction, GFP expression was quantified *via* flow cytometry (Novocyte) and populations with approximately 50% GFP expression were expanded and selected using 250 nM JQ1, 1 μM PCIP-1, 1 μM *ent*-PCIP-1, or 1 μM talazoparib for $PARP1^{KO}$ or 1 μM JQ1, PCIP-1, *ent*-PCIP-1, or talazoparib for $PARP2^{KO}$. These doses were selected from approximate $IC_{50}$ values in Jurkat cells treated for 72 h *via* viability assays. Cells were collected for GFP analysis in triplicate and sub-cultured every 3-5 days for up to 21 days post transduction. Relative % GFP from averages of triplicates are plotted over time with error bars showing standard error the mean (s.e.m.). Conditions with fewer than 5% viable cells were excluded from the plots. After 21 days, triplicate wells of 1 μM talazoparib selected $PARP1^{KO}$ JURKAT cells were combined and expanded in the presence of 1 μM talazoparib to maintain selective pressure. As a control, AAVS1's triplicates were pooled and expanded. After 14 days of selection with PCIP-1, PARP2 $^{KO}$ cells were cultured in the absence of PCIP-1 for 14 days and expanded for anti-proliferation assays.

**Viability assay for Jurkat-Cas9-$PARP1^{KO}$ and Jurkat-Cas9-$PARP2^{KO}$ cells:** JURKAT-Cas9 cells selected for $PARP1^{KO}$ with 1 μM talazoparib were cultured in absence of talazoparib for 72 hours prior to seeding cells for the anti-proliferation assays. JURKAT-Cas9 cells selected for $PARP2^{KO}$ with 1 μM PCIP-1 were cultured in absence of PCIP-1 for 2 weeks prior to seeding cells for the anti-proliferation assays, in order to recover the viability of the populations. 30 nL of serial half-log diluted compounds were printed into white 384 well plates (Corning REF 353988) using an 655 Echo Acoustic Liquid Handler (Beckman). 3,400 cells in 30 μL of media were plated in triplicate. 15 μL of ATPLite reagent diluted 1:4 with MilliQ water was added to wells after 72 hours of growth and plates were incubated for 10 minutes in the dark at room temperature while shaking. Fluorescence values were measured using the Clariostar Plus microplate reader (BMG Labtech). The average of triplicate values are plotted with error bars representing standard error of mean (s.e.m.).

**PARP2 target engagement assay:** Was carried out by Reaction Biology using the following protocol "HEK293 cells transiently expressing PARP2-NanoLuc Fusion Vector were seeded into the wells of

384-well plates. The cells were pre-treated with PARP Tracer-01 and then treated with reference compound olaparib for 1 hour. The BRET signal was measured on an Envision 2104 Multilabel Reader." (https://www.reactionbiology.com/datasheet/parp2-nano/)

**Homogenous time-resolved FRET (HTRF) assay**: HTRF assays were performed in assay buffer (25 mM HEPES pH 7, 20 mM NaCl, 0.2% Pluronic F-127, and 0.05% BSA). Assays consisted of 1 nM LanthaScreen Eu-anti-His Tag antibody (ThermoFisher, Cat. No, PV5597), and 8.9 nM SureLight allophycocyanin-streptavidin (PerkinElmer, APC-SA, Cat. No. CR130-100). 5 µL was dispensed per well into black 1536-well plates (Corning, Cat. # 9007BC) *via* Multidrop Combi reagent dispenser (Thermo). Compound addition was performed with the 655 Echo Acoustic Liquid Handler (Beckman). Plates were incubated for 2 hours in the dark after compound administration before measurement *via* PHERAstar plate reader (BMG Labtech; simultaneous dual emission; excitation = 337 nm, emission 1 = 620 nm, emission 2 = 665 nm).

For the BRD4_BD1-H4 tetra-acetylated peptide HTRF consisted of 10 nM BRD4_BD1 (BPS Bioscience, Cat. # 31042) and 13.3 nM tetra-acetylated H4 (BioVision Cat. No. 7144−01). HTRF signals (ration of emission 2 to emission 1) from DMSO-treated wells (maximum signal control) and no-peptide control wells (minimum signal control) were used for percent inhibition calculations. For the ternary complex HTRF, 50 nM PARP1_Cat and 25 nM BRD4_BD1 were added simultaneously. HTRF signals were compared to DMSO-treated wells (minimum signal control).

**Viability assays for homologous recombination deficient cell lines:** Cell viability was assessed using the CellTiter-Glo luminescent cell viability assay (Promega) in 96-well opaque white polystyrene microplates (Costar 3917, non-clear bottom). Peripheral wells were filled with media only and excluded from analysis to buffer against edge effects due to evaporation. Each experimental well contained 100 µL of media and cells seeded at the following densities:

- DLD1 and BRCA2$^{ko}$ cells: 1,400 cells/well
- HCT116 and BRCA2$^{ko}$ cells: 900 cells/well
- RPE1-hTERT-Flag-Cas9-TP53$^{ko}$ and derived lines (BRCA1$^{ko}$, BRCA1/53BP$^{ko}$): 800 cells/well

Cells were allowed to adhere and recover before drug treatment. For compound dosing, the media in each well was first aspirated and replaced with 100 µL of fresh culture media. Drug-containing media were prepared at 2X concentration and added at 100 µL per well, resulting in a final volume of 200 µL and a 1X final drug concentration. Eight-point, 4-fold serial dilutions were prepared, with final compound

concentrations ranging from 10 µM down to 0.152 nM, plus a vehicle-only (0 µM) control. Cells were treated with compounds for 6 days. At endpoint, media were aspirated from all wells and replaced with 100 µL of fresh media. An additional 100 µL of fresh media was added to four wells containing no cells to serve as blank controls. Subsequently, 100 µL of CellTiter-Glo reagent was added to each well, followed by a brief incubation according to the manufacturer's instructions.
Luminescence was measured using a BioTek Synergy Neo plate reader (Gen 5.2.09). Three technical replicates per treatment condition. The mean of technical replicates was plotted, with error bars representing the standard error of the mean (s.e.m.).

**Cell culture**

The following cell lines were used in this work: Jurkat, Jurkat-BRD4-HiBit, Jurkat-PARP1-HiBit, 22Rv1-FKBP12$^{F36V}$-2XHA-BRD4 Jurkat-PARP1$^{KO}$, Jurkat-PARP2$^{KO}$, Jurkat-Cas9, Jurkat-Cas9-PARP1$^{KO}$ (sgRNA1-3), Jurkat-Cas9-PARP2$^{KO}$ (sgRNA 1-4), RPE1-hTERT *TP53-KO Flag-Cas9* (RPE1-P53$^{KO}$), RPE1-P53$^{KO}$-BRCA1$^{KO}$, RPE1-P53$^{KO}$-BRCA1$^{KO}$-53BP1$^{KO}$, DLD1, DLD1-BRCA2$^{KO}$, HCT116, and HCT116-BRCA2$^{KO}$.

All Jurkat isogenic cell lines were cultured in RPMI1640 media containing 10% heat-inactivated fetal bovine serum (FBS) and 5% Antibiotic-Antimycotic (Thermofisher, #15240062) and stored at 37°C at 5% $CO_2$. The parental line was a gift from the lab of Professor Michael Bollong's lab at Scripps Research.

22Rv1-FKBP12$^{F36V}$-2XHA-BRD4 cells were cultured in RPMI1640 media containing 10% heat-inactivated fetal bovine serum (FBS) and were supplemented with 5% Antibiotic-Antimycotic (Thermofisher, #15240062) and stored at 37°C at 5% $CO_2$.

All RPE1-hTERT TP53$^{KO}$ isogenic cell lines were cultured in DMEM +1% penicillin/streptomycin and 2 µg/mL blasticidin and stored at 37°C at 5% $CO_2$. The parental cell line RPE1-hTERT was purchased from ATCC and the ensuing genetic alterations are described in.[1]

All DLD1 isogenic cell lines were cultured in RPMI-1640 +1% penicillin/streptomycin +10% FBS and stored 37°C at 5% $CO_2$, DLD1-BRCA2$^{KO}$ storage was supplemented with 3% $O_2$. These isogenic cell lines were purchased from Horizon.

All HCT116 isogenic cell lines were cultured in ATCC-formulated McCoy's 5a medium +1% penicillin/streptomycin +10% FBS and stored 37°C at 5% $CO_2$, HCT116-BRCA2$^{KO}$ storage was supplemented with 3% $O_2$. HCT116 cells were purchased from ATCC and the BRCA2$^{KO}$ was originally purchased from Ximbio.

**Immunoblot analysis**

At time of cell harvest, 3-6 million cells per condition were isolated. The cells were spun down (500 g x 5 min) and then given washed with 1 mL of PBS before being spun down again. Cells were flash frozen in liquid nitrogen before protein isolation. Protein was isolated by 30-minute treatment on ice of 100 µL RIPA lysis buffer (ThermoFisher, Cat # 89900) with 1x Halt protease inhibitor (ThermoFisher, Cat # 78429) and 1:1000 benzonase nuclease (Sigma-Aldrich, Cat # 70746-4). Cleared protein lysates were isolated from cell debris through centrifugation (16000 g x 10 min @ 4°C). Protein concentrations were obtained through BCA assay (ThermoFisher, Cat # 23225). 4x SDS sample buffer and 10% 2-mercaptoethanol were added to each sample and then protein was denatured at 95°C for 10 minutes. 16 µg of each sample was loaded into a well of a 4-12% Bis-Tris gel (ThermoFisher, Cat # NW04127). Protein ladders (BioRad, Cat # 1610375) were added on both sides of the samples. Protein gels were run at 90 volts for 10 minutes and then 134 volts for 42 minutes in MES buffer (Invitrogen, Cat # B000202). Proteins were transferred to nitrocellulose membranes and then blocked in 5% nonfat milk in TBS-T for 1 hour at RT. Primary antibodies were incubated over the membranes overnight at 4°C with light shaking. Following TBS-T washes, secondary infrared antibodies (IRDye) were incubated with light shaking for 1 hour at room temperature. Following final washes, blots were imaged on the Odyssey CLx Images (LI-COR). The following primary antibodies were used for overnight blotting: anti-GAPDH (Sigma-Aldrich, Cat # G8795), anti-PARP1 (CST, Cat # 9542), anti-PAR (CST, Cat # 89190), anti-γH2A.X (CST, Cat # 2595S), anti-cleaved caspase 3 (CST, Cat # 9661), anti-Cleaved Caspase 7 (CST, Cat # 9491).

**Co-immunoprecipitation**

22Rv1FKBP12$^{F36V}$-2xHA-BRD4 cells expressing BRD4_HA_dTAG were treated with compound for 4 hours. Cells were washed with cold PBS and scraped to detach cells from the dish. Following a PBS wash and centrifugation (500 g x 5 min at 4°C), cells were lysed with a buffer containing Cell Lytic M (Sigma-Aldrich, Cat # C2978) with 1x Halt protease inhibitor (ThermoFisher, Cat # 78429) and 1:1000 benzonase nuclease (Sigma-Aldrich, Cat # 70746-4). Pellets were sonicated and placed to rotate at 4°C for 1 hour. Cleared protein lysates were isolated from cell debris through centrifugation (16000 g x 10 min @ 4°C). Protein concentrations were obtained through BCA assay (ThermoFisher, Cat # 23225). Input samples were prepared at 4 mg/mL with 50 µL aliquots removed for Western blotting. Anti-HA magnetic beads (Sigma-Aldrich, Cat # 88836) were washed with Cell Lytic M and 25 µL beads per 1 mL of protein was added to each sample. Beads and protein inputs were incubated with rotation overnight at 4°C. Beads were then collected through magnetization and washed 3 times with TBS-T.

Protein was eluted by adding 40 µL 1X loading buffer (1X SDS loading buffer + 2.5% 2-mercaptoethanol) and heated at 95°C for 10 minutes. Input samples were diluted to 2 mg/mL. 4x SDS sample buffer and 10% 2-mercaptoethanol was then added to input samples and heated at 95°C for 10 minutes. 18 µL of each sample and input was loaded into a well of a 4-12% Bis-Tris gel (ThermoFisher, Cat # NW04127). Protein gels were run at 90 volts for 10 minutes and then 150 volts for 50 minutes in MES buffer (Invitrogen, Cat # B000202). Proteins were transferred to nitrocellulose membranes and then blocked in 5% nonfat milk in TBS-T for 1 hour at RT. Primary antibodies were incubated over the membranes overnight at 4°C with light shaking. Following TBS-T washes, secondary infrared antibodies (IRDye) were incubated with light shaking for 1 hour at RT. Following final washes, blots were imaged on the Odyssey CLx Images (LI-COR). The following primary antibodies were used for overnight blotting: anti-GAPDH (Sigma-Aldrich, Cat # G8795), anti-PARP1 (CST, Cat # 9542), anti-BRD4 (Bethyl, Cat # BL-151-6F11), and anti-HA (CST, Cat # 3724).

**Cell cycle analysis**

100,000 cells were seeded per well in a 96-well plate and treated with compounds as a 1:1000 dilution of DMSO stocks. Following incubation, plates were spun at 500 g for 5 minutes. Following two wash steps with PBS, cells were fixed overnight with cold 70% ethanol at 4°C overnight. Cells were then centrifuged at 1000 g for 5 minutes and washed twice with PBS. Cells were stained in the dark at room temperature in 100 µL PBS with 0.5 µg/mL RNAse A (Thermo, Cat. # EN0531) and 50 µg/mL propidium iodide (VWR, Cat. # 102876-812). Propidium iodide quantification was performed *via* flow cytometry on a Novocyte 3000 (Agilent).

**Protein production**

The catalytic domain of PARP1 (amino acids 661-1014) was recombinantly expressed and purified in BL21 (DE3) cells (New England Biolabs). The protein was cloned into a pET28(a)+ expression vector with an N-terminal 6xHis tag and a C-terminal AviTag. BL21 (DE3) cells were co-transformed with the AviTag-containing construct and a biotin ligase expression vector, pBirAcm (Avidity). Bacteria was induced at an OD600 of 0.6 at 16°C by 1 mM isopropyl-β-D-thiogalactopyranoside (IPTG) (Fisher Scientific, Cat. # 50-213-380). Following a 20-hour incubation, the culture was harvested *via* centrifugation. The cell pellet was resuspended in lysis buffer (50mM sodium phosphate (pH 7.4), 500 mM NaCl, 10mM imidazole, 1x Halt Protease Inhibitor Cocktail (Thermo Scientific, Cat. # 78429) and passed three times through a microfluidizer (Microfluidics). Following centrifugation at 16,000 g for 20 minutes at 4°C, the supernatant was incubated with TALON metal affinity resin (Fisher Scientific, Cat. # NC9306569) for 1 hour at 4°C. The resin was then placed over an elution column where it was washed 2x with lysis buffer before elution *via* 1mL fractions of elution buffer (50 mM sodium phosphate (pH 7.4), 500 mM NaCl, 300 mM imidazole, 1x Halt Protease Inhibitor Cocktail. Fractions containing protein underwent buffer exchange to size exclusion buffer (20mM Tris-HCl (pH 7.5) in a 10K desalting column (Amnicon Ultracel). Purified protein was then further purified through a HiLoad Superdex column on an AKTA pure size exclusion system (Cytiva).

**Synthesis**

Scheme S1. Synthetic route to parent ligands.

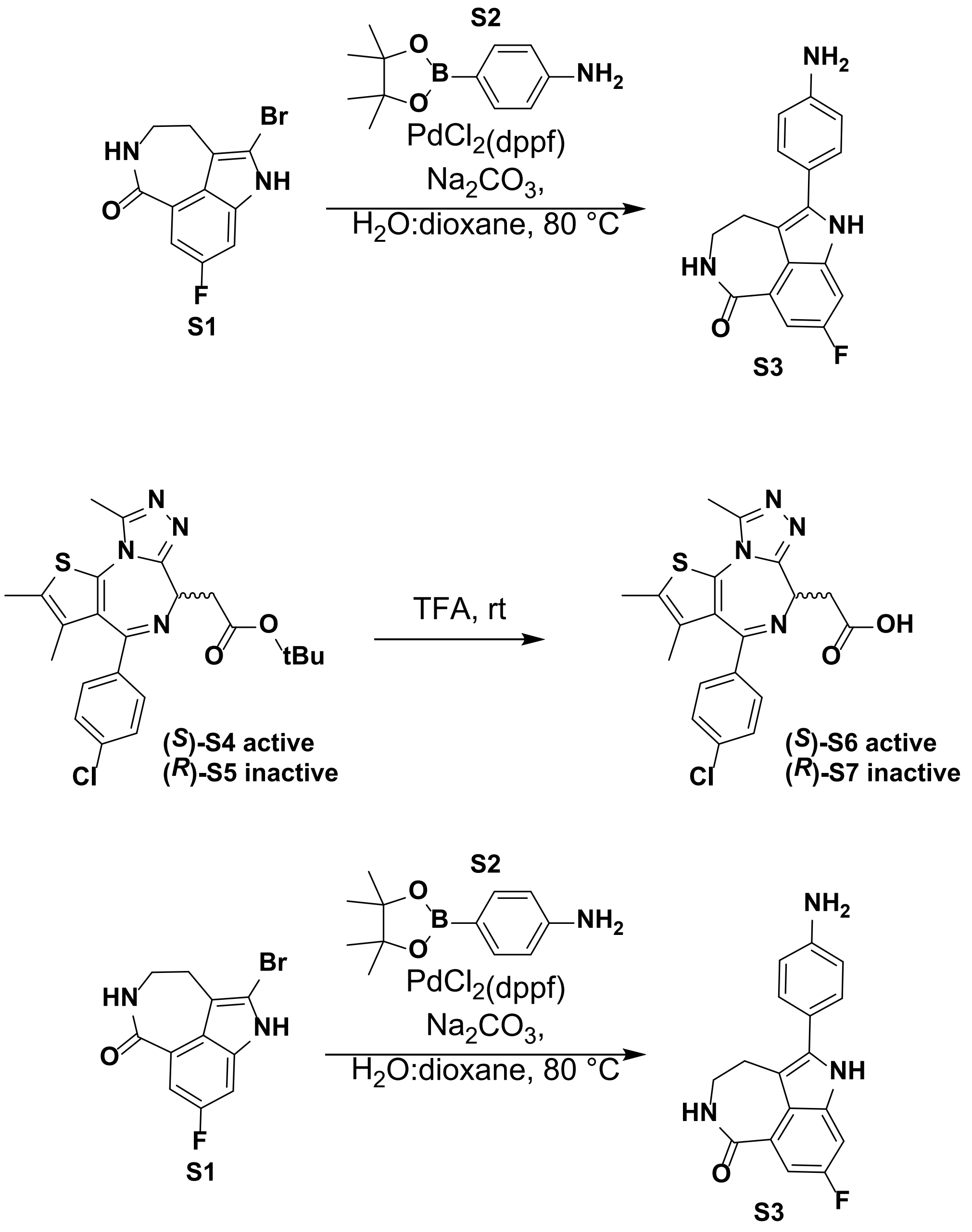

Synthesis of 5-(4-aminophenyl)-8-fluoro-2,3,4,6-tetrahydro-1H-azepino[5,4,3-cd]indol-1-one (**S3**): A 25 mL round bottom flask was equipped with a magnetic stir bar and charged with 5-bromo-8-fluoro-2,3,4,6-tetrahydro-1H-azepino[5,4,3-cd]indol-1-one (**S1**) (500 mg, 1.7 mmol), 4-(4,4,5,5-tetramethyl-1,3,2-dioxaborolan-2-yl)aniline (**S2**) (464 mg, 2.1 mmol), and sodium carbonate (350 mg, 3.5 mmol) before the flask was sealed with a rubber septum. The solid was suspended in 8 mL of dioxane and 500 µL of water which was subsequently degassed using a ballon of nitrogen and vent needle over the course of twenty minutes. Ten minutes into the degassing procedure 1,1′-Bis(diphenylphosphino)ferrocene]dichloropalladium(II) (65 mg, 0.08 mmol) was added portion wise. After degassing for an additional twenty minutes the needles were removed from the flask septum and the reaction was heated using an oil bath to 80 °C and heated for 16 h. Once the reaction had reached completion the mixture was allowed to cool to room temperature before being transferred to a 1 L beaker by washing and diluting 800 mL of ethyl acetate. The organic layer was sequentially washed in a 1 L separatory funnel with water (3 × 5 mL) and saturated aqueous sodium bicarbonate (3 × 5 mL), followed by a brine wash (3 × 5 mL), and then dried over magnesium sulfate. The dried organic solution was flushed through a basic alumina plug and the volatiles were removed using rotary evaporation. This residue was dissolved in N,N-dimethylformamide (1.0 mL) and dry loaded onto celite before being packed into a plastic cartridge which was directly loaded onto a 42-gram RediSep C18 reversed phase column equipped to a Combiflash NextGen 300+ auto column. The crude reaction mixture was purified using an acidic gradient (0.1% trifluoracetic acid) ranging from 10-100% water/acetonitrile. The gradient began during the first fraction and concluded after the 25-minute run. Clean fractions were determined using UPLC technologies and subsequently lyophilized to yield a beige powder (210 mg, 40%). Spectra is consistent with primary literature.[2]

$^1$H NMR (400 MHz, DMSO-$d_6$) δ 11.46 (s, 1H), 8.20 (t, *J* = 5.8 Hz, 1H), 7.45 – 7.35 (m, 3H), 7.26 (dd, *J* = 9.2, 2.4 Hz, 1H), 6.86 (d, *J* = 8.1 Hz, 2H), 3.44 – 3.34 (m, 2H), 3.04 – 2.95 (m, 2H).

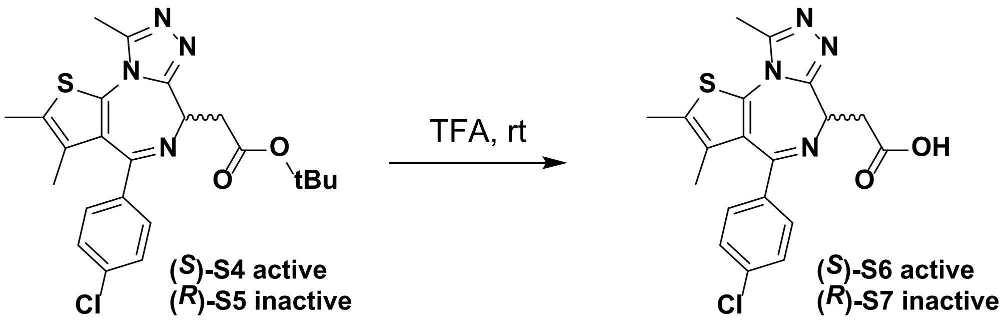

Synthesis of 4 (*S*)-2-(4-(4-chlorophenyl)-2,3,9-trimethyl-6H-thieno[3,2-f][1,2,4]triazolo[4,3-a][1,4]diazepin-6-yl)acetic acid **((*S*)-S6)**: A 20 mL dram vial was equipped with a magnetic stir bar and charged with tert-butyl tert-butyl (*S*)-2-(4-(4-chlorophenyl)-2,3,9-trimethyl-6H-thieno[3,2-f][1,2,4]triazolo[4,3-a][1,4]diazepin-6-yl)acetate **((*S*)-S4)**: (500 mg, 1.1 mmol) and was then dissolved in trifluoroacetic acid (3 mL), before being stirred at room temperature. The reaction was monitored using UPLC and following completion after two hours the trifluoroacetic acid was removed using reduced pressure. This residue was dissolved in N,N-dimethylformamide (0.5 mL) and loaded directly onto a 42-gram RediSep C18 reversed phase column equipped to a Combiflash NextGen 300+ auto column. The crude reaction mixture was purified using an acidic gradient (0.1% trifluoracetic acid) ranging from 10-100% water/acetonitrile. The gradient began during the first fraction and concluded after the 25-minute run. Clean fractions were determined using UPLC technologies and subsequently lyophilized to yield a yellow powder (401 mg, 91%).

$^1$H NMR (600 MHz, DMSO-$d_6$) δ 7.56 – 7.52 (m, 2H), 7.48 (d, *J* = 8.2 Hz, 2H), 4.50 (t, *J* = 7.0 Hz, 1H), 3.50 – 3.43 (m, 1H), 3.40 – 3.33 (m, 1H), 2.65 (d, *J* = 1.3 Hz, 3H), 2.54 (p, *J* = 1.8 Hz, 2H), 2.45 (s, 3H), 2.12 (d, *J* = 1.3 Hz, 1H), 1.67 (s, 3H).

Synthesis of 4 (*R*)-2-(4-(4-chlorophenyl)-2,3,9-trimethyl-6H-thieno[3,2-f][1,2,4]triazolo[4,3-a][1,4]diazepin-6-yl)acetic acid **((*R*)-S7)**: The procedure and outcomes are identical to the synthesis of **((*S*)-S6)**.

Scheme S2. Representative routes to rucaparib-JQ1 based heterobifunctional molecules.

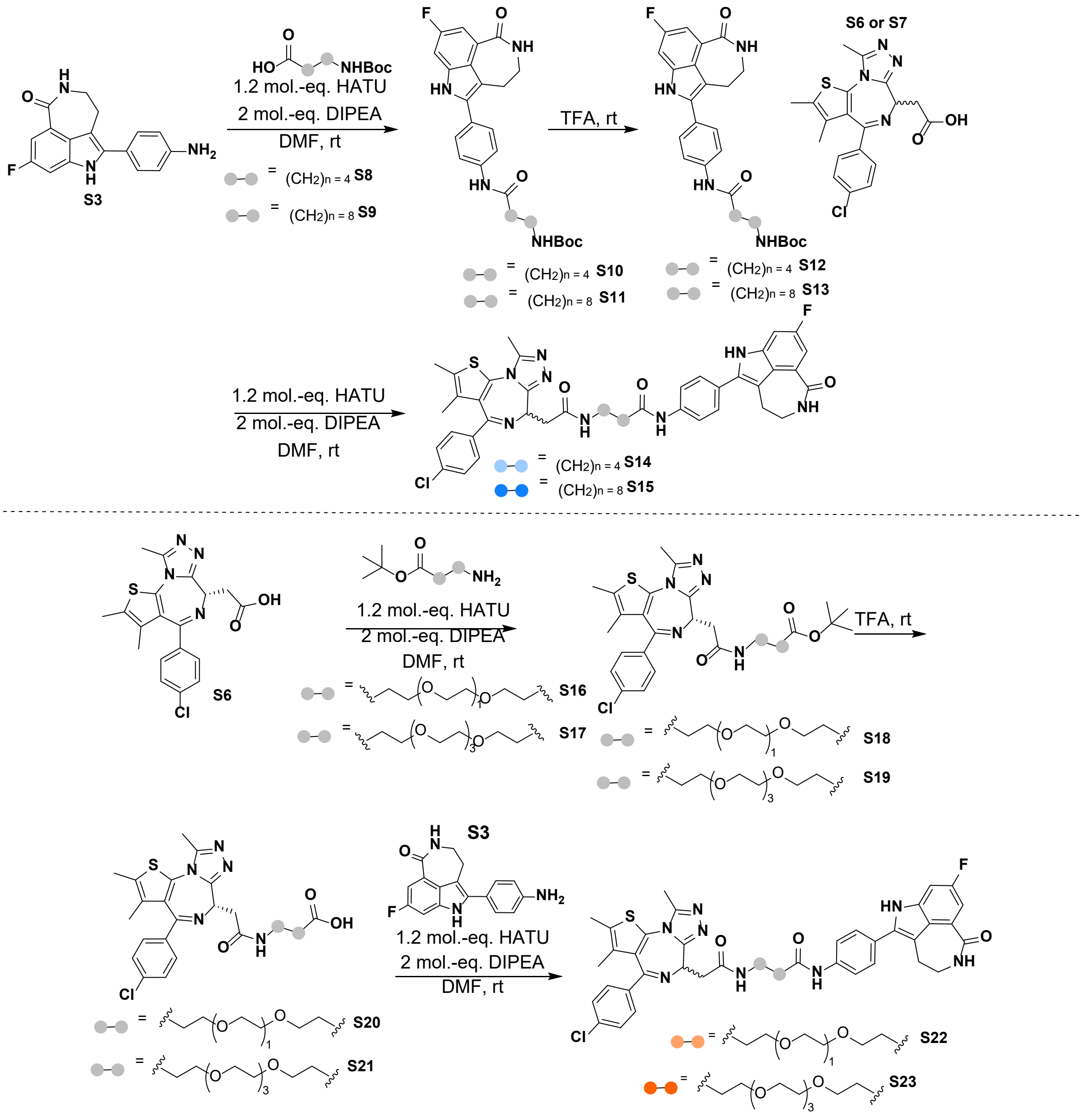

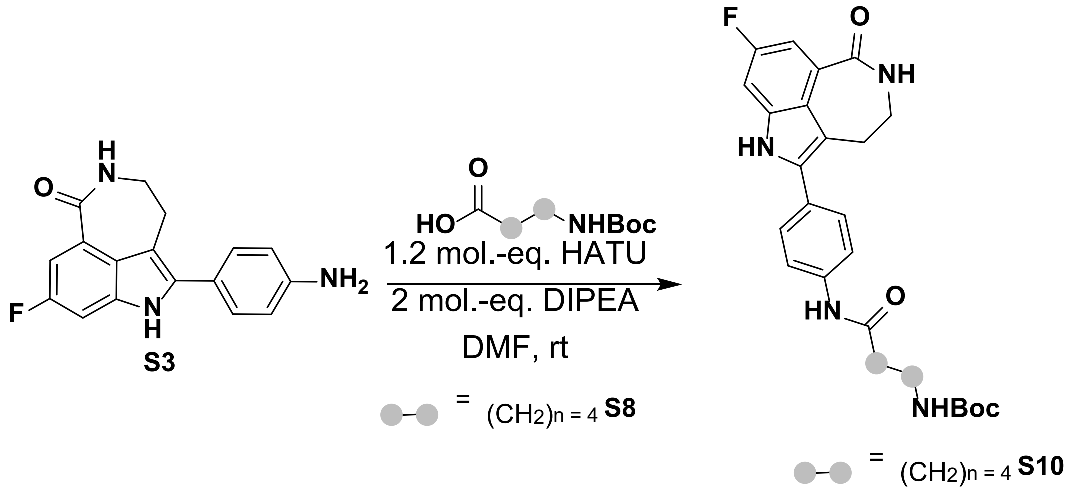

Synthesis of tert-butyl (5-((4-(8-fluoro-1-oxo-2,3,4,6-tetrahydro-1H-azepino[5,4,3-cd]indol-5-yl)phenyl)amino)-5-oxopentyl)carbamate (**S10**): A 3.0 mL dram vial equipped with a stir bar was charged with 5-((tert-butoxycarbonyl)amino)pentanoic acid (**S8**) (122 mg, 0.51 mmol), HATU (232 mg, 0.61 mmol), N,N-dimethylformamide (1.0 mL), and N,N-diisopropylethylamine (177 µL, 1.0 mmol). The solution was allowed to stir for 10 minutes before the portion wise addition of 5-(4-aminophenyl)-8-fluoro-2,3,4,6-tetrahydro-1H-azepino[5,4,3-cd]indol-1-one (**S3**) (150 mg, 0.51 mmol). Stirring continued until UPLC analysis demonstrated complete consumption of the starting materials. The reaction reached completion after five and a half hours of stirring at room temperature. The crude reaction solution was directly loaded onto a 12-gram RediSep C18 reversed phase column equipped to a Combiflash NextGen 300+ auto column. The crude reaction mixture was purified using an acidic gradient (0.1% trifluoracetic acid) ranging from 10-100% water/acetonitrile. The gradient began during the first fraction and concluded after the 25-minute run. Clean fractions were determined using UPLC technologies and subsequently lyophilized to yield a white powder (111 mg, 44%).

$^1$H NMR (600 MHz, DMSO-$d_6$) δ 11.63 (s, 1H), 10.05 (s, 1H), 8.25 (t, *J* = 5.9 Hz, 1H), 7.75 (d, *J* = 8.3 Hz, 2H), 7.57 (d, *J* = 8.3 Hz, 2H), 7.41 (dd, *J* = 11.0, 2.5 Hz, 1H), 7.30 (dd, *J* = 9.1, 2.5 Hz, 1H), 6.83 (t, *J* = 5.7 Hz, 1H), 3.39 (s, 2H), 3.03 (t, *J* = 4.9 Hz, 2H), 2.94 (q, *J* = 6.6 Hz, 2H), 2.34 (t, *J* = 7.4 Hz, 2H), 1.58 (p, *J* = 7.6 Hz, 2H), 1.42 (q, *J* = 7.6 Hz, 2H), 1.38 (s, 9H).

$^{13}$C NMR (151 MHz, DMSO-$d_6$) δ 170.74, 167.86, 158.44, 156.89, 154.99, 150.55, 138.31, 136.01, 134.74, 127.66, 125.61, 125.02, 122.69, 118.45, 110.52, 108.71, 99.96, 76.75, 41.26, 35.50, 28.56, 28.17, 27.67, 21.86.

$^{19}F$ NMR (376 MHz, DMSO-$d_6$, referenced to $C_6F_6$): δ -123.91.

HRMS (ESI-TOF) m/z: $[M+H]^+$ calculated for $C_{27}H_{32}FN_4O_4$: 495.2403, found: 495.2409.

UPLC-MS, $ESI^+$, *m/z* 495.31 $[M+H]^+$

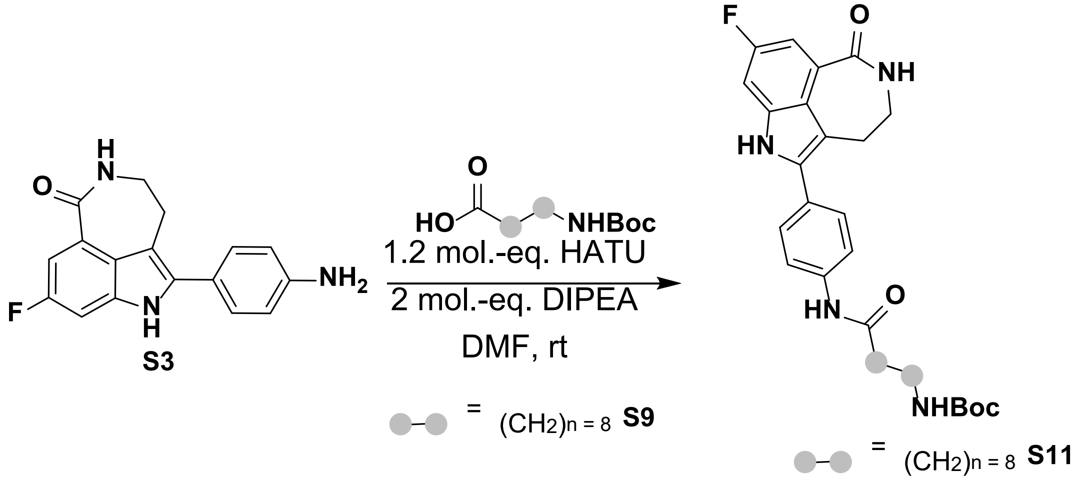

Synthesis of tert-butyl (9-((4-(8-fluoro-1-oxo-2,3,4,6-tetrahydro-1H-azepino[5,4,3-cd]indol-5-yl)phenyl)amino)-9-oxononyl)carbamate (**S11**): A 3.0 mL dram vial equipped with a stir bar was charged with 9-((tert-butoxycarbonyl)amino)nonanoic acid (**S9**) (110 mg, 0.51 mmol), HATU (232 mg, 0.61 mmol), N,N-dimethylformamide (1.0 mL), and N,N-diisopropylethylamine (177 µL, 1.0 mmol). The solution was allowed to stir for 10 minutes before the portion wise addition of 5-(4-aminophenyl)-8-fluoro-2,3,4,6-tetrahydro-1H-azepino[5,4,3-cd]indol-1-one (**S3**) (150 mg, 0.51 mmol). Stirring continued until UPLC analysis demonstrated complete consumption of the starting materials. The reaction reached completion after five and a half hours of stirring at room temperature. The crude reaction solution was directly loaded onto a 12-gram RediSep C18 reversed phase column equipped to a Combiflash NextGen 300+ auto column. The crude reaction mixture was purified using an acidic gradient (0.1% trifluoracetic acid) ranging from 10-100% water/acetonitrile. The gradient began during the first fraction and concluded after the 25-minute run. Clean fractions were determined using UPLC technologies and subsequently lyophilized to yield a white powder (154 mg, 55%).

$^1H$ NMR (400 MHz, DMSO-$d_6$) δ 11.63 (s, 1H), 10.05 (s, 1H), 8.26 (t, *J* = 5.7 Hz, 1H), 7.75 (d, *J* = 8.4 Hz, 2H), 7.57 (d, *J* = 8.5 Hz, 2H), 7.41 (dd, *J* = 11.0, 2.5 Hz, 1H), 7.31 (dd, *J* = 9.1, 2.5 Hz, 1H), 6.78 (s, 1H), 3.39 (s, 2H), 3.03 (s, 2H), 2.89 (q, *J* = 6.6 Hz, 2H), 2.33 (t, *J* = 7.4 Hz, 2H), 1.60 (s, 2H), 1.37 (s, 9H), 1.27 (br, 9H).

$^{13}C$ NMR (151 MHz, DMSO-$d_6$) δ 172.06, 169.77, 159.64, 158.08, 156.12, 138.98, 136.89, 135.58, 128.38, 126.82, 125.34, 123.63, 119.54, 111.17, 110.16, 109.99, 101.00, 100.83, 42.58, 42.44, 37.16, 37.11, 29.96, 29.32, 29.20, 29.13, 28.63, 26.78, 25.60.

$^{19}F$ NMR (376 MHz, DMSO-$d_6$, referenced to $C_6F_6$): δ -123.9.

HRMS (ESI-TOF) m/z: $[M+H]^+$ calculated for $C_{31}H_{40}FN_4O_4$: 551.3029, found: 551.3028.

UPLC-MS, $ESI^+$, *m/z* 551.47 $[M+H]^+$

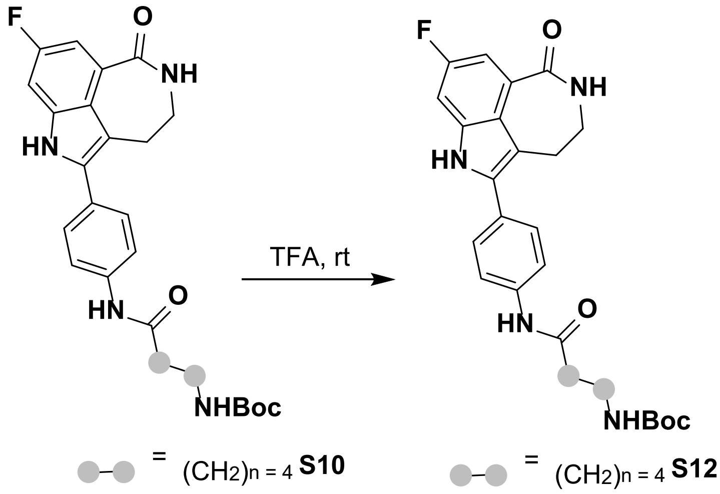

= $(CH_2)_{n=4}$ **S10** = $(CH_2)_{n=4}$ **S12**

Synthesis of 5-amino-N-(4-(8-fluoro-1-oxo-2,3,4,6-tetrahydro-1H-azepino[5,4,3-cd]indol-5-yl)phenyl)pentanamide (**S12**): A 3.0 mL dram vial equipped with a stir bar was charged with tert-butyl (5-((4-(8-fluoro-1-oxo-2,3,4,6-tetrahydro-1H-azepino[5,4,3-cd]indol-5-yl)phenyl)amino)-5-oxopentyl)carbamate (**S10**) (45 mg, 91 µmol) and trifluoracetic acid (2.0 mL). The solution was allowed to stir at room temperature for 15.5 h, reaction completion was determined using UPLC, and before the solvent was removed using reduced pressure. The crude reaction solution was dissolved in N,N-dimethylformamide (1.0 mL) and directly loaded onto a 12-gram RediSep C18 reversed phase column equipped to a Combiflash NextGen 300+ auto column. The crude reaction mixture was purified using an acidic gradient (0.1% trifluoracetic acid) ranging from 10-100% water/acetonitrile. The gradient began during the first fraction and concluded after the 25-minute run. Clean fractions were determined using UPLC technologies and subsequently lyophilized to yield a white powder (25 mg, 70%).

$^{1}$H NMR (600 MHz, DMSO-$d_6$) δ 11.66 (s, 1H), 10.15 (s, 1H), 8.26 (t, $J$ = 5.8 Hz, 1H), 7.76 (d, $J$ = 8.7 Hz, 2H), 7.58 (d, $J$ = 8.5 Hz, 2H), 7.42 (dd, $J$ = 11.0, 2.5 Hz, 1H), 7.32 (dd, $J$ = 9.0, 2.5 Hz, 1H), 3.40 (s, 2H), 3.03 (br, 2H), 2.83 (br, 2H), 2.39 (t, $J$ = 7.1 Hz, 2H), 1.74 – 1.54 (m, 4H).

$^{13}$C NMR (151 MHz, DMSO-$d_6$) δ 171.51, 168.95, 159.53, 157.97, 139.30, 137.18, 135.77, 128.76, 126.79, 123.75, 119.54, 111.62, 109.97, 101.06, 42.33, 39.17, 36.17, 29.26, 27.09, 22.41.

$^{19}$F NMR (376 MHz, DMSO-$d_6$, referenced to $C_6F_6$): δ -123.83.

HRMS (ESI-TOF) m/z: $[M+H]^+$ calculated for $C_{22}H_{24}FN_4O_2$: 395.1878, found: 395.1890.

UPLC-MS, $ESI^+$, *m/z* 395.31 $[M+H]^+$

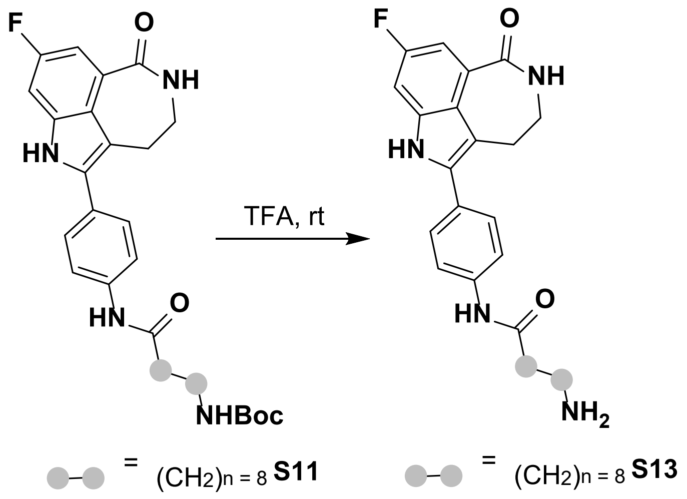

Synthesis of 9-amino-N-(4-(8-fluoro-1-oxo-2,3,4,6-tetrahydro-1H-azepino[5,4,3-cd]indol-5-yl)phenyl)nonanamide (**S13**): A 3.0 mL dram vial equipped with a stir bar was charged with tert-butyl (5-((4-(8-fluoro-1-oxo-2,3,4,6-tetrahydro-1H-azepino[5,4,3-cd]indol-5-yl)phenyl)amino)-5-oxopentyl)carbamate (**S11**) (134 mg, 243 μmol) and trifluoracetic acid (2.0 mL). The solution was allowed to stir at room temperature for 15.5 h, reaction completion was determined using UPLC, and before the solvent was removed using reduced pressure. The crude reaction solution was dissolved in N,N-dimethylformamide (1.0 mL) and directly loaded onto a 12-gram RediSep C18 reversed phase column equipped to a Combiflash NextGen 300+ auto column. The crude reaction mixture was purified using an acidic gradient (0.1% trifluoracetic acid) ranging from 10-100% water/acetonitrile. The gradient began during the first fraction and concluded after the 25-minute run. Clean fractions were determined using UPLC technologies and subsequently lyophilized to yield a white powder (79 mg, 72%).

$^{1}H$ NMR (600 MHz, DMSO-$d_6$) δ 11.65 (s, 1H), 10.11 (s, 1H), 8.19 (t, $J$ = 5.8 Hz, 1H), 7.70 (d, $J$ = 8.7 Hz, 2H), 7.50 (d, $J$ = 8.7 Hz, 2H), 7.34 (dd, $J$ = 11.0, 2.5 Hz, 1H), 7.25 (dd, $J$ = 9.1, 2.5 Hz, 1H), 3.32 (s, 2H), 2.96 (s, 2H), 2.64 (t, $J$ = 7.5 Hz, 2H), 2.28 (t, $J$ = 7.4 Hz, 2H), 1.54 (t, $J$ = 7.2 Hz, 2H), 1.44 (t, $J$ = 7.5 Hz, 2H), 1.23 (s, 10H).

$^{13}C$ NMR (151 MHz, DMSO-$d_6$) δ 171.98, 168.97, 159.49, 157.94, 139.46, 137.20, 135.84, 128.71, 126.64, 126.00, 123.76, 119.52, 111.53, 109.92, 100.89, 42.34, 36.89, 29.26, 29.14, 29.05, 28.98, 28.60, 26.38, 25.55.

$^{19}F$ NMR (376 MHz, DMSO-$d_6$, referenced to $C_6F_6$): δ -123.96.

HRMS (ESI-TOF) m/z: $[M+H]^+$ calculated for $C_{26}H_{32}FN_4O_2$: 451.2504, found: 451.2509.

UPLC-MS, $ESI^+$, $m/z$ 451.37 $[M+H]^+$

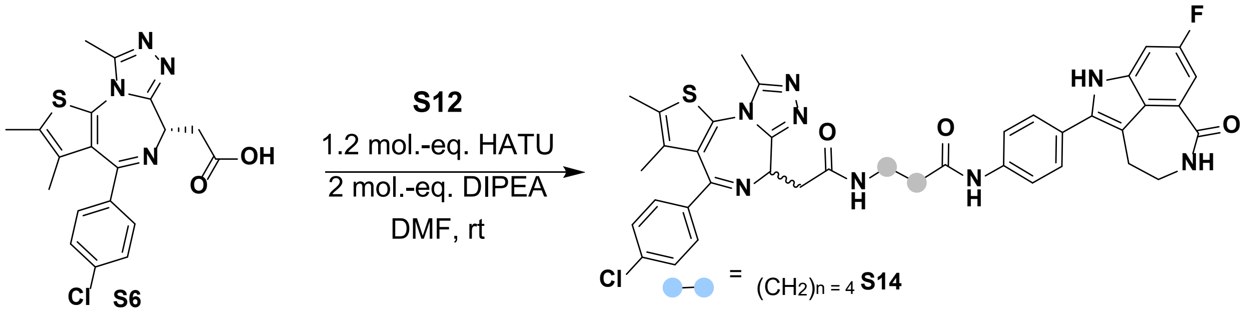

Synthesis of ($S$)-5-(2-(4-(4-chlorophenyl)-2,3,9-trimethyl-6H-thieno[3,2-f][1,2,4]triazolo[4,3-a][1,4]diazepin-6-yl)acetamido)-N-(4-(8-fluoro-1-oxo-2,3,4,6-tetrahydro-1H-azepino[5,4,3-cd]indol-5-yl)phenyl)pentanamide (**S14**, compound **1**): A 3.0 mL dram vial equipped with a stir bar was charged with ($S$)-4-(4-chlorophenyl)-2,3,9-trimethyl-6H-thieno[3,2-f][1,2,4]triazolo[4,3-a][1,4]diazepine-6-carboxylic acid (**S6**) (19 mg, 0.05 mmol), HATU (22 mg, 0.06 mmol), N,N-dimethylformamide (0.5 mL), and N,N-diisopropylethylamine (27 µL, 0.15 mmol). The solution was allowed to stir for 10 minutes before the portion wise addition of 5-amino-N-(4-(8-fluoro-1-oxo-2,3,4,6-tetrahydro-1H-azepino[5,4,3-cd]indol-5-yl)phenyl)pentanamide (**S12**) (19 mg, 0.05 mmol). Stirring continued until UPLC analysis demonstrated complete consumption of the starting materials. The reaction reached completion after 10 minutes of stirring at room temperature and was allowed to continue stirring for an additional 20 minutes. The crude reaction solution was directly loaded onto a 12-gram RediSep C18 reversed phase column equipped to a Combiflash NextGen 300+ auto column. The crude reaction mixture was purified using an acidic gradient (0.1% trifluoracetic acid) ranging from 10-100% water/acetonitrile. The gradient began during the first fraction and concluded after the 25-minute run. Clean fractions were determined using UPLC technologies and subsequently lyophilized to yield a yellow powder (8 mg, 20%).

$^{1}$H NMR (600 MHz, DMSO-$d_6$) δ 11.63 (s, 1H), 10.08 (s, 1H), 8.25 (q, *J* = 5.7 Hz, 2H), 7.76 (d, *J* = 8.9 Hz, 2H), 7.56 (d, *J* = 8.9 Hz, 2H), 7.48 (d, *J* = 8.9 Hz, 2H), 7.42 (s, 3H), 7.31 (dd, *J* = 9.1, 2.4 Hz, 1H), 4.52 (dd, *J* = 8.6, 5.7 Hz, 1H), 3.39 (s, 2H), 3.31 – 3.15 (m, 4H), 3.13 – 3.08 (m, 1H), 3.03 (s, 2H), 2.60 (s, 3H), 2.43 – 2.36 (m, 5H), 1.66 (td, 2H), 1.61 (s, 3H), 1.52 (p, *J* = 7.2 Hz, 2H).

$^{13}$C NMR (151 MHz, DMSO-$d_6$) δ 170.69, 168.75, 167.84, 162.50, 158.41, 156.86, 154.44, 149.30, 138.29, 136.04, 134.63, 131.60, 130.17, 129.51, 129.22, 128.95, 127.85, 127.62, 125.59, 124.93, 122.67, 118.41, 115.76, 113.83, 110.49, 108.86, 99.94, 53.23, 41.23, 37.65, 36.98, 35.49, 28.25, 28.16, 21.93, 13.42, 12.04, 10.68.

$^{19}$F NMR (376 MHz, DMSO-$d_6$, referenced to $C_6F_6$): δ -123.87.

HRMS (ESI-TOF) m/z: $[M+H]^+$ calculated for $C_{41}H_{39}ClFN_8O_3S$: 777.2533, found: 777.2533

UPLC-MS, ESI$^+$, *m/z* 777.49 $[M]^+$

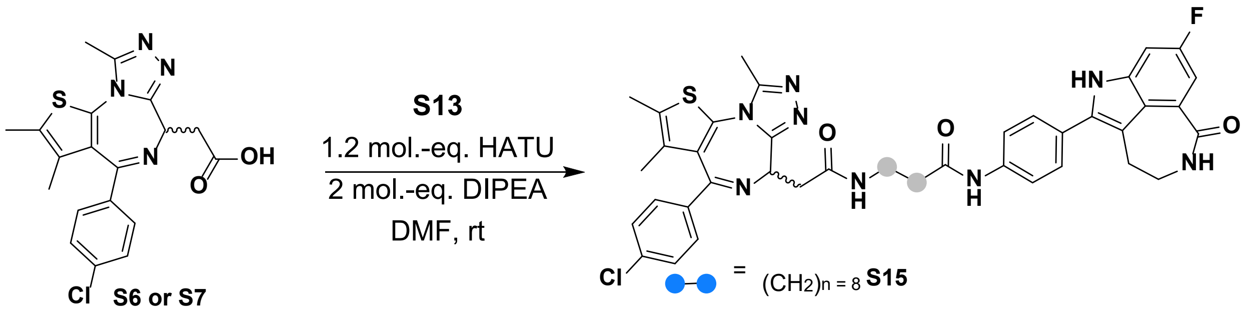

Synthesis of (*S* or *R*)-9-(2-(4-(4-chlorophenyl)-2,3,9-trimethyl-6H-thieno[3,2-f][1,2,4]triazolo[4,3-a][1,4]diazepin-6-yl)acetamido)-N-(4-(8-fluoro-1-oxo-2,3,4,6-tetrahydro-1H-azepino[5,4,3-cd]indol-5-

yl)phenyl)nonanamide (**S15,** PCIP-1): A 3.0 mL dram vial equipped with a stir bar was charged with (*S*)-4-(4-chlorophenyl)-2,3,9-trimethyl-6H-thieno[3,2-f][1,2,4]triazolo[4,3-a][1,4]diazepine-6-carboxylic acid (**S6 or S7**) (8.6 mg, 0.02 mmol), HATU (10 mg, 0.03 mmol), N,N-dimethylformamide (0.5 mL), and N,N-diisopropylethylamine (12 μL, 0.07 mmol). The solution was allowed to stir for 10 minutes before the portion wise addition of 9-amino-N-(4-(8-fluoro-1-oxo-2,3,4,6-tetrahydro-1H-azepino[5,4,3-cd]indol-5-yl)phenyl)nonanamide (**S13**) (10 mg, 0.02 mmol). Stirring continued until UPLC analysis demonstrated complete consumption of the starting materials. The reaction reached completion after 10 minutes of stirring at room temperature and was allowed to continue stirring for an additional 20 minutes. The crude reaction solution was directly loaded onto a 12-gram RediSep C18 reversed phase column equipped to a Combiflash NextGen 300+ auto column. The crude reaction mixture was purified using an acidic gradient (0.1% trifluoracetic acid) ranging from 10-100% water/acetonitrile. The gradient began during the first fraction and concluded after the 25-minute run. Clean fractions were determined using UPLC technologies and subsequently lyophilized to yield an orange powder (16 mg, 86%, when using **S6**). The synthesis of compound **8** (ent-PCIP-1) was carried out under conditions identical to those used for PCIP-1, yielding the same outcomes.

$^{1}$H NMR (600 MHz, DMSO-$d_6$) δ 11.61 (s, 1H), 10.03 (s, 1H), 8.23 (t, *J* = 5.7 Hz, 1H), 8.17 (t, *J* = 5.7 Hz, 1H), 7.75 (d, *J* = 8.7 Hz, 2H), 7.56 (d, *J* = 8.6 Hz, 2H), 7.48 (d, *J* = 8.7 Hz, 2H), 7.42 (m, 3H), 7.30 (d, *J* = 9.1 Hz, 1H), 4.53 (t, *J* = 6.6 Hz, 1H), 3.38 (s, 2H), 3.25 (dd, J = 15.0, 8.3 Hz, 1H), 3.12 (m, 3H), 3.02 (t, J = 4.9 Hz, 2H), 2.60 (s, 3H), 2.40 (s, 3H), 2.32 (t, *J* = 7.4 Hz, 2H), 1.62 (s, 5H), 1.44 (s, 2H), 1.29 (s, 10H).

$^{13}$C NMR (151 MHz, DMSO-$d_6$) δ 171.92, 169.75, 168.94, 163.55, 157.96, 155.56, 150.38, 139.42, 137.10, 135.77, 132.69, 131.30, 130.63, 130.30, 130.06, 128.92, 128.73, 126.65, 126.04, 123.77, 119.51, 116.85, 114.92, 111.59, 101.04, 54.34, 42.33, 38.92, 38.06, 36.94, 29.72, 29.30, 29.24, 29.16, 29.14, 26.85, 25.58, 14.52, 13.14, 11.76.

$^{19}$F NMR (376 MHz, DMSO-$d_6$, referenced to $C_6F_6$): δ -123.91.

HRMS (ESI-TOF) m/z: $[M+H]^+$ calculated for $C_{45}H_{47}ClFN_8O_3S$: 833.3159, found: 833.3157.

UPLC-MS, ESI$^+$, *m/z* 833.57 $[M]^+$

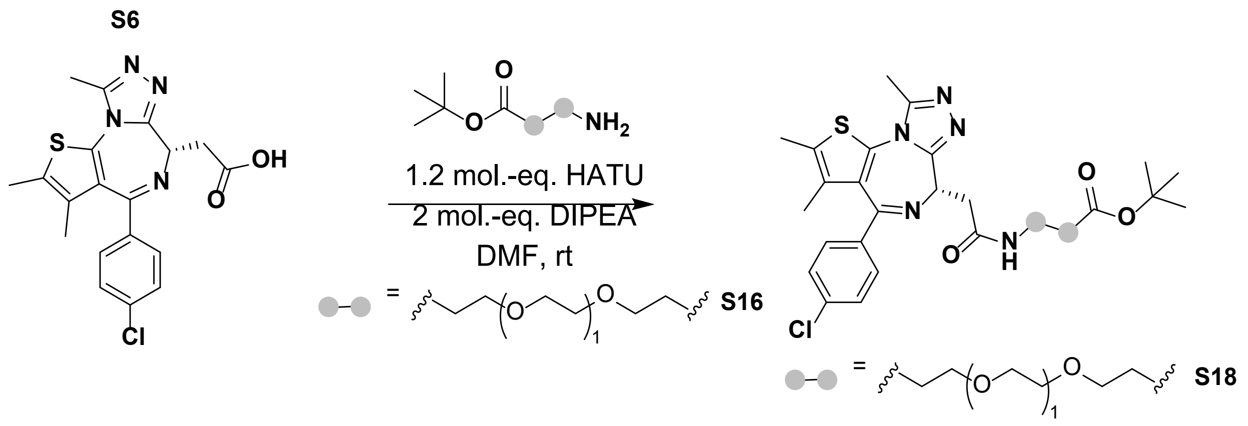

Synthesis of tert-butyl (*S*)-3-(2-(2-(2-(4-(4-chlorophenyl)-2,3,9-trimethyl-6H-thieno[3,2-f][1,2,4]triazolo[4,3-a][1,4]diazepin-6-yl)acetamido)ethoxy)ethoxy)propanoate (**S18**): A 3.0 mL dram vial equipped with a stir bar was charged with (*S*)-4-(4-chlorophenyl)-2,3,9-trimethyl-6H-thieno[3,2-f][1,2,4]triazolo[4,3-a][1,4]diazepine-6-carboxylic acid (**S6**) (200 mg, 0.5 mmol), HATU (228 mg, 0.06 mmol), N,N-dimethylformamide (1.0 mL), and N,N-diisopropylethylamine (174 µL, 1.0 mmol). The solution was allowed to stir for 10 minutes before the portion wise addition of tert-butyl 3-(2-(2-aminoethoxy)ethoxy)propanoate (**S16**) (140 mg, 0.05 mmol). Stirring continued until UPLC analysis demonstrated complete consumption of the starting materials. The reaction reached completion after one hour of stirring at room temperature. The crude reaction solution was directly loaded onto a 12-gram RediSep C18 reversed phase column equipped to a Combiflash NextGen 300+ auto column. The crude reaction mixture was purified using an acidic gradient (0.1% trifluoracetic acid) ranging from 10-100% water/acetonitrile. The gradient began during the first fraction and concluded after the 35-minute run. Clean fractions were determined using UPLC technologies and subsequently lyophilized to yield a colorless oil (35 mg, 81%).

$^{1}$H NMR (600 MHz, DMSO-*d6*) δ 8.29 (s, 1H), 7.49 (d, *J* = 8.9 Hz, 2H), 7.43 (d, *J* = 8.9 Hz, 2H), 4.54 – 4.47 (m, 1H), 3.60 (s, 3H), 3.25 (s, 5H), 2.60 (s, 3H), 2.42 (s, 5H), 1.63 (s, 3H), 1.39 (s, 9H).

$^{13}$C NMR (151 MHz, DMSO-*d6*) δ 170.89, 170.13, 163.49, 155.57, 150.31, 137.21, 135.69, 132.74, 131.19, 130.64, 130.29, 130.02, 128.93, 80.20, 70.08, 69.99, 69.65, 66.70, 54.29, 39.07, 37.95, 36.29, 28.22, 14.54, 13.16, 11.78.

HRMS (ESI-TOF) m/z: [M+H]$^+$ calculated for $C_{30}H_{39}ClN_5O_5S$: 616.2355, found 616.2358.

UPLC-MS, ESI$^+$, $m/z$ 616.45 $[M]^+$

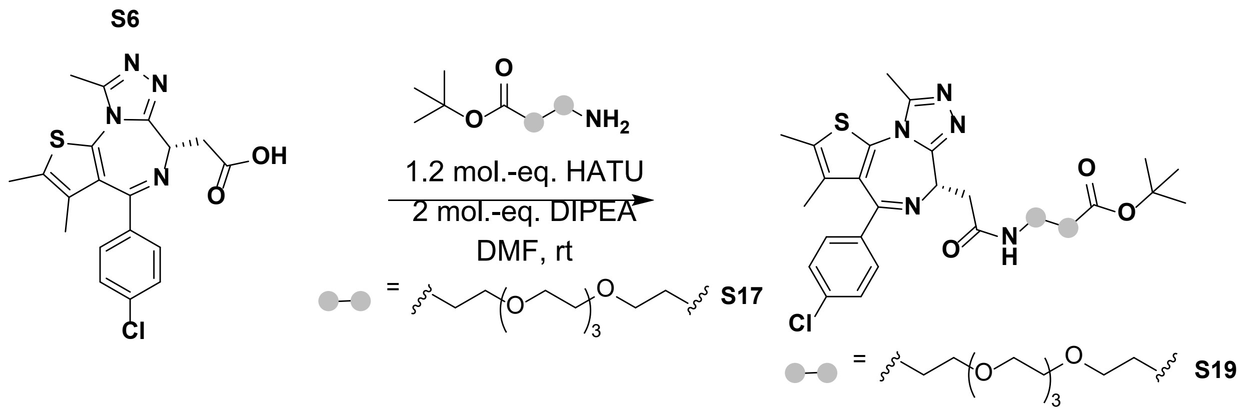

Synthesis of tert-butyl (*S*)-1-(4-(4-chlorophenyl)-2,3,9-trimethyl-6H-thieno[3,2-f][1,2,4]triazolo[4,3-a][1,4]diazepin-6-yl)-2-oxo-6,9,12,15-tetraoxa-3-azaoctadecan-18-oate (**S19**): A 3.0 mL dram vial equipped with a stir bar was charged with (*S*)-4-(4-chlorophenyl)-2,3,9-trimethyl-6H-thieno[3,2-f][1,2,4]triazolo[4,3-a][1,4]diazepine-6-carboxylic acid (**S6**) (200 mg, 0.5 mmol), HATU (228 mg, 0.06 mmol), N,N-dimethylformamide (1.0 mL), and N,N-diisopropylethylamine (174 µL, 1.0 mmol). The solution was allowed to stir for 10 minutes before the portion wise addition of tert-butyl 1-amino-3,6,9,12-tetraoxapentadecan-15-oate (**S17**) (160 mg, 0.05 mmol). Stirring continued until UPLC analysis demonstrated complete consumption of the starting materials. The reaction reached completion after one hour of stirring at room temperature. The crude reaction solution was directly loaded onto a 12-gram RediSep C18 reversed phase column equipped to a Combiflash NextGen 300+ auto column. The crude reaction mixture was purified using an acidic gradient (0.1% trifluoracetic acid) ranging from 10-100% water/acetonitrile. The gradient began during the first fraction and concluded after the 35-minute run. Clean fractions were determined using UPLC technologies and subsequently lyophilized to yield a colorless oil (122 mg, 35%).

$^1$H NMR (400 MHz, $CDCl_3$) δ 7.40 (d, *J* = 8.5 Hz, 2H), 7.32 (d, *J* = 8.5 Hz, 2H), 7.20 (s, 1H), 4.66 (t, *J* = 7.1 Hz, 1H), 3.66 – 3.50 (m, 16H), 3.36 (dd, *J* = 14.7, 7.0 Hz, 1H), 2.65 (s, 3H), 2.48 (t, *J* = 6.4 Hz, 2H), 2.39 (s, 3H), 1.67 (s, 3H), 1.42 (s, 9H).

$^{13}$C NMR (151 MHz, $CDCl_3$) δ 171.00, 170.65, 163.81, 155.70, 149.81, 136.72, 136.67, 132.16, 130.93, 130.74, 130.50, 129.88, 128.69, 80.53, 70.54, 70.51, 70.44, 70.40, 70.30, 69.90, 66.87, 54.39, 39.40, 39.02, 36.19, 28.09, 14.43, 13.11, 11.86.

HRMS (ESI-TOF) m/z: $[M+H]^+$ calculated for HRMS (ESI-TOF) m/z: $[M+H]^+$ calculated for $C_{34}H_{47}ClN_5O_7S$: 704.2880, found 704.2883.

UPLC-MS, $ESI^+$, *m/z* 704.52 $[M]^+$

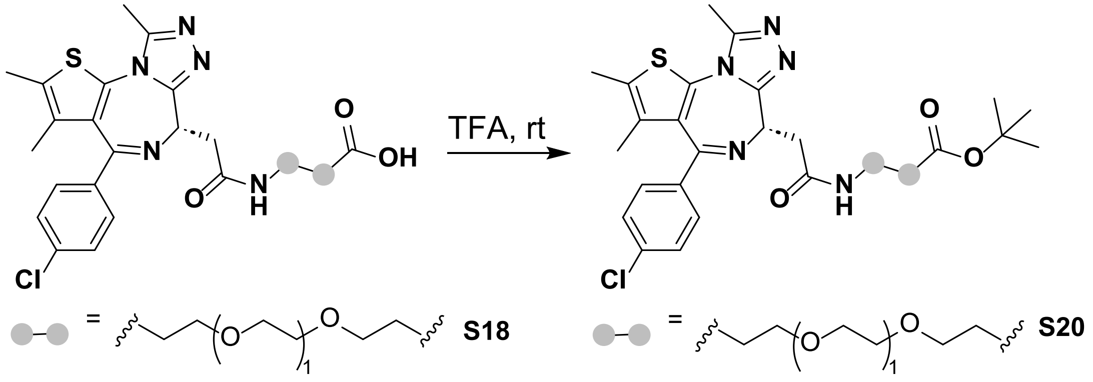

Synthesis of (*S*)-3-(2-(2-(2-(4-(4-chlorophenyl)-2,3,9-trimethyl-6H-thieno[3,2-f][1,2,4]triazolo[4,3-a][1,4]diazepin-6-yl)acetamido)ethoxy)ethoxy)propanoic acid (**S20**)**:** A 3.0 mL dram vial equipped with a stir bar was charged tert-butyl 3-(2-(2-(2-((6*S*)-4-(4-chlorophenyl)-2,3,9-trimethyl-3a,10a-dihydro-6H-thieno[3,2-f][1,2,4]triazolo[4,3-a][1,4]diazepin-6-yl)acetamido)ethoxy)ethoxy)propanoate (**S18**) (95 mg, 0.15 mmol) and trifluoracetic acid (2.0 mL). The solution was allowed to stir at room temperature for 4h, reaction completion was determined using UPLC, before removing the solvent using reduced pressure. The crude reaction solution was dissolved in N,N-dimethylformamide (1.0 mL) and directly loaded onto a 12-gram RediSep C18 reversed phase column equipped to a Combiflash NextGen 300+ auto column. The crude reaction mixture was purified using an acidic gradient (0.1% trifluoracetic acid) ranging from 10-100% water/acetonitrile. The gradient began during the first fraction and concluded after the 35-minute run. Clean fractions were determined using UPLC technologies and subsequently lyophilized to yield a colorless oil (79 mg, 92%).

$^1H$ NMR (600 MHz, DMSO-$d_6$) δ 8.29 (t, $J$ = 5.8 Hz, 1H), 7.49 (d, $J$ = 8.6 Hz, 2H), 7.43 (d, $J$ = 8.5 Hz, 2H), 4.52 (dd, $J$ = 8.3, 5.9 Hz, 1H), 3.61 (t, $J$ = 6.4 Hz, 2H), 3.45 (t, $J$ = 5.9 Hz, 2H), 3.33 – 3.19 (m, 4H), 2.61 (s, 3H), 2.45 (t, $J$ = 6.4 Hz, 2H), 2.42 (s, 3H), 1.63 (s, 3H).

$^{13}C$ NMR (151 MHz, DMSO-$d_6$) δ 172.06, 169.01, 162.52, 154.46, 149.35, 136.05, 134.69, 131.62, 130.24, 129.61, 129.27, 129.00, 127.87, 68.98, 68.92, 68.60, 65.64, 53.17, 38.01, 36.80, 34.13, 13.47, 12.10, 10.70.

HRMS (ESI-TOF) m/z: $[M+H]^+$ calculated for $C_{26}H_{31}ClN_5O_5S$: 616.2355, found 616.2358.

UPLC-MS, $ESI^+$, *m/z* 560.35 $[M]^+$

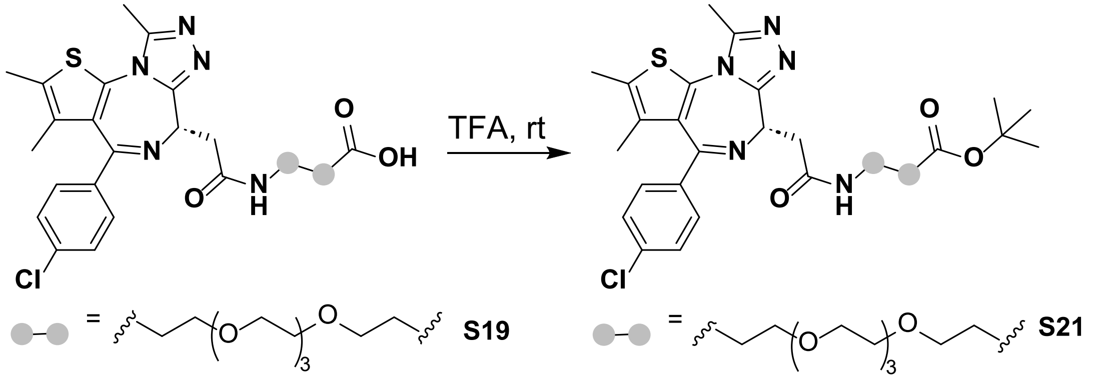

Synthesis of (*S*)-1-(4-(4-chlorophenyl)-2,3,9-trimethyl-6H-thieno[3,2-f][1,2,4]triazolo[4,3-a][1,4]diazepin-6-yl)-2-oxo-6,9,12,15-tetraoxa-3-azaoctadecan-18-oic acid (**S21**): A 3.0 mL dram vial equipped with a stir bar was charged tert-butyl 3-(2-(2-(2-((6*S*)-4-(4-chlorophenyl)-2,3,9-trimethyl-3a,10a-dihydro-6H-thieno[3,2-f][1,2,4]triazolo[4,3-a][1,4]diazepin-6-yl)acetamido)ethoxy)ethoxy)propanoate (**S19**) (113 mg, 0.16 mmol) and trifluoracetic acid (2.0 mL). The solution was allowed to stir at room temperature for 4h, reaction completion was determined using UPLC, before removing the solvent using reduced pressure. The crude reaction solution was dissolved in N,N-dimethylformamide (1.0 mL) and directly loaded onto a 12-gram RediSep C18 reversed phase column equipped to a Combiflash NextGen 300+ auto column. The crude reaction mixture was purified using an acidic gradient (0.1% trifluoracetic acid) ranging from 10-100% water/acetonitrile. The gradient began during the first fraction and concluded after the 35-minute run. Clean fractions were determined using UPLC technologies and subsequently lyophilized to yield a colorless oil (79 mg, 92%).

$^1$H NMR (600 MHz, DMSO-$d_6$) δ 8.23 (t, *J* = 5.7 Hz, 1H), 7.42 (d, *J* = 8.7 Hz, 2H), 7.36 (d, *J* = 8.6 Hz, 2H), 4.44 (dd, *J* = 8.2, 5.9 Hz, 1H), 3.52 (t, *J* = 6.4 Hz, 2H), 3.49 – 3.35 (m, 16H), 3.24 – 3.12 (m, 4H), 2.53 (s, 3H), 2.38 – 2.33 (m, 5H), 1.56 (s, 3H).

$^{13}C$ NMR (151 MHz, DMSO-$d_6$) δ 173.11, 170.11, 163.53, 155.55, 150.35, 137.18, 135.72, 132.73, 131.24, 130.65, 130.31, 130.04, 128.94, 70.25, 70.21, 70.16, 70.09, 69.69, 66.69, 54.27, 39.09, 37.92, 35.19, 14.54, 13.16, 11.78.

HRMS (ESI-TOF) m/z: $[M+H]^+$ calculated for $C_{30}H_{39}ClN_5O_7S$: 648.2254, found 648.2246.

UPLC-MS, $ESI^+$, *m/z* 648.42 $[M]^+$

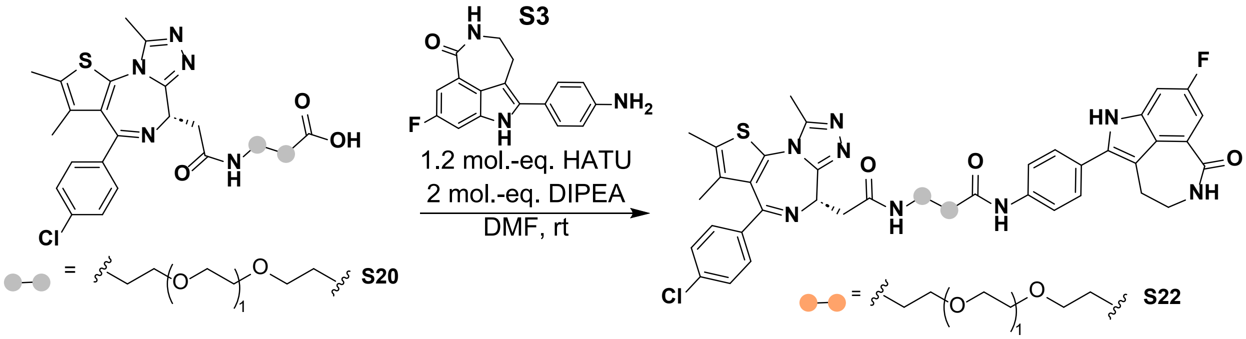

Synthesis of (*S*)-1-(2-(4-(4-chlorophenyl)-2,3,9-trimethyl-6H-thieno[3,2-f][1,2,4]triazolo[4,3-a][1,4]diazepin-6-yl)acetamido)-N-(4-(8-fluoro-1-oxo-2,3,4,6-tetrahydro-1H-azepino[5,4,3-cd]indol-5-yl)phenyl)-3,6,9,12-tetraoxapentadecan-15-amide (**S22**, compound **3**) A 3.0 mL dram vial equipped with a stir bar was charged with 3-(2-(2-((6*S*)-4-(4-chlorophenyl)-2,3,9-trimethyl-3a,5,6,10a-tetrahydro-4H-thieno[3,2-f][1,2,4]triazolo[4,3-a]azepine-6-carboxamido)ethoxy)ethoxy)propanoic acid (**S20**) (51 mg, 0.10 mmol), HATU (42 mg, 0.12 mmol), N,N-dimethylformamide (1.0 mL), and N,N-diisopropylethylamine (52 µL, 0.30 mmol). The solution was allowed to stir for 10 minutes before the portion wise addition of 5-(4-aminophenyl)-8-fluoro-2,3,4,6-tetrahydro-1H-azepino[5,4,3-cd]indol-1-one (**S3**) (27 mg, 0.10 mmol). Stirring continued until UPLC analysis demonstrated complete consumption of the starting materials. The reaction reached completion after three and a half hours of stirring at room temperature. The crude reaction solution was directly loaded onto a 12-gram RediSep C18 reversed phase column equipped to a Combiflash NextGen 300+ auto column. The crude reaction mixture was purified using an acidic gradient (0.1% trifluoracetic acid) ranging from 10-100% water/acetonitrile. The gradient began during the first fraction and concluded after the 35-minute run. Clean fractions were determined using UPLC technologies and subsequently lyophilized to yield a yellow powder (25 mg, 32%).

$^1$H NMR (600 MHz, DMSO-$d_6$) δ 11.62 (s, 1H), 10.13 (s, 1H), 8.28 (t, *J* = 5.7 Hz, 1H), 8.24 (t, *J* = 5.8 Hz, 1H), 7.75 (d, *J* = 8.6 Hz, 2H), 7.56 (d, *J* = 8.7 Hz, 2H), 7.49 – 7.45 (m, 2H), 7.44 – 7.39 (m, 3H), 7.30 (dd, *J* = 9.1, 2.4 Hz, 1H), 4.50 (dd, *J* = 8.1, 6.1 Hz, 1H), 3.73 (d, *J* = 6.2 Hz, 2H), 3.55 (t, *J* = 2.5 Hz, 4H), 3.45 (t, *J* = 5.9 Hz, 2H), 3.37 (t, 2H), 3.29 – 3.20 (m, 4H), 3.01 (s, 2H), 2.64 – 2.56 (m, 5H), 2.39 (s, 3H), 1.60 (s, 3H).

$^{13}$C NMR (151 MHz, DMSO-$d_6$) δ 169.06, 168.83, 167.84, 162.41, 154.47, 149.23, 138.16, 136.11, 134.61, 131.65, 130.09, 129.20, 128.92, 127.84, 127.68, 125.73, 124.97, 122.68, 120.54, 118.46, 116.72, 114.78, 110.55, 108.70, 99.95, 69.04, 68.92, 68.60, 66.00, 53.20, 41.24, 38.00, 36.86, 36.62, 28.15, 13.43, 12.06, 10.69.

$^{19}$F NMR (376 MHz, DMSO-$d_6$, referenced to $C_6F_6$): δ -122.06.

HRMS (ESI-TOF) m/z: $[M+H]^+$ calculated for $C_{43}H_{43}ClFN_8O_5S$: 837.2735, found: 837.2738.

UPLC-MS, $ESI^+$, *m/z* 837.54 $[M]^+$

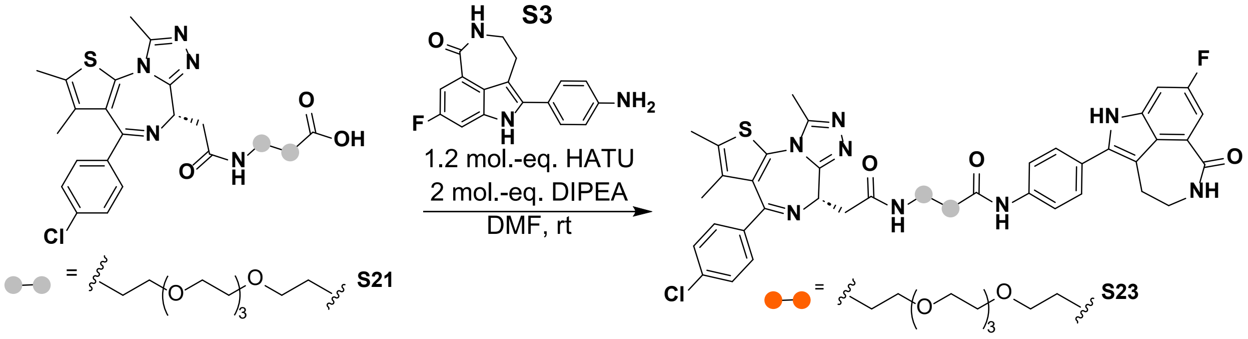

Synthesis of (*S*)-1-(2-(4-(4-chlorophenyl)-2,3,9-trimethyl-6H-thieno[3,2-f][1,2,4]triazolo[4,3-a][1,4]diazepin-6-yl)acetamido)-N-(4-(8-fluoro-1-oxo-2,3,4,6-tetrahydro-1H-azepino[5,4,3-cd]indol-5-yl)phenyl)-3,6,9,12-tetraoxapentadecan-15-amide (**S23**, compound **4**): A 3.0 mL dram vial equipped with a stir bar was charged with 3-(2-(2-((6*S*)-4-(4-chlorophenyl)-2,3,9-trimethyl-3a,5,6,10a-tetrahydro-4H-thieno[3,2-f][1,2,4]triazolo[4,3-a]azepine-6-carboxamido)ethoxy)ethoxy)propanoic acid (**S21**) (66 mg, 0.10 mmol), HATU (46 mg, 0.12 mmol), N,N-dimethylformamide (1.0 mL), and N,N-diisopropylethylamine (57 µL, 0.33 mmol). The solution was allowed to stir for 10 minutes before the portion wise addition of 5-(4-aminophenyl)-8-fluoro-2,3,4,6-tetrahydro-1H-azepino[5,4,3-cd]indol-1-one (**S3**) (30 mg, 0.10 mmol). Stirring continued until UPLC analysis demonstrated complete consumption of the starting materials. The reaction reached completion after three and a half hours of stirring at room temperature. The crude reaction solution was directly loaded onto a 12-gram RediSep C18 reversed phase column equipped to a Combiflash NextGen 300+ auto column. The crude reaction mixture was purified using an acidic gradient (0.1% trifluoracetic acid) ranging from 10-100% water/acetonitrile. The gradient began during the first fraction and concluded after the 35-minute run. Clean fractions were determined using UPLC technologies and subsequently lyophilized to yield a yellow powder (25 mg, 32%).

$^{1}$H NMR (600 MHz, DMSO-$d_6$) δ 11.63 (s, 1H), 10.12 (s, 1H), 8.30 (t, $J$ = 5.7 Hz, 1H), 8.24 (t, $J$ = 5.8 Hz, 1H), 7.75 (d, $J$ = 8.5 Hz, 2H), 7.56 (d, $J$ = 8.5 Hz, 2H), 7.48 (d, $J$ = 8.6 Hz, 2H), 7.44 – 7.39 (m, 3H), 7.30 (dd, $J$ = 9.1, 2.5 Hz, 1H), 4.51 (dd, $J$ = 8.1, 6.0 Hz, 1H), 3.71 (t, $J$ = 6.2 Hz, 2H), 3.62 (dt, $J$ = 10.9, 6.7 Hz, 1H), 3.55 – 3.49 (m, 10H), 3.44 (t, $J$ = 5.9 Hz, 2H), 3.38 (s, 2H), 3.29 – 3.19 (m, 4H), 3.02 (s, 2H), 2.64 – 2.56 (m, 5H), 2.40 (s, 3H), 1.61 (s, 3H).

$^{13}$C NMR (151 MHz, DMSO-$d_6$) δ 169.02, 168.78, 167.82, 162.43, 158.41, 156.86, 154.43, 149.25, 138.16, 136.06, 134.61, 131.61, 130.12, 129.54, 129.19, 128.91, 127.83, 127.64, 125.70, 124.94, 122.65, 118.41, 115.95, 114.01, 110.52, 108.68, 99.93, 69.14, 69.10, 69.05, 68.97, 68.55, 65.97, 53.15, 41.22, 37.98, 36.80, 36.58, 28.14, 20.34, 19.55, 13.41, 12.03, 10.66.

$^{19}$F NMR (376 MHz, DMSO-$d_6$, referenced to $C_6F_6$): δ -123.87.

HRMS (ESI-TOF) m/z: $[M+H]^+$ calculated for $C_{47}H_{51}ClFN_8O_7S$: 925.3269, found: 925.3271.

UPLC-MS, ESI$^+$, *m/z* 925.64 $[M]^+$

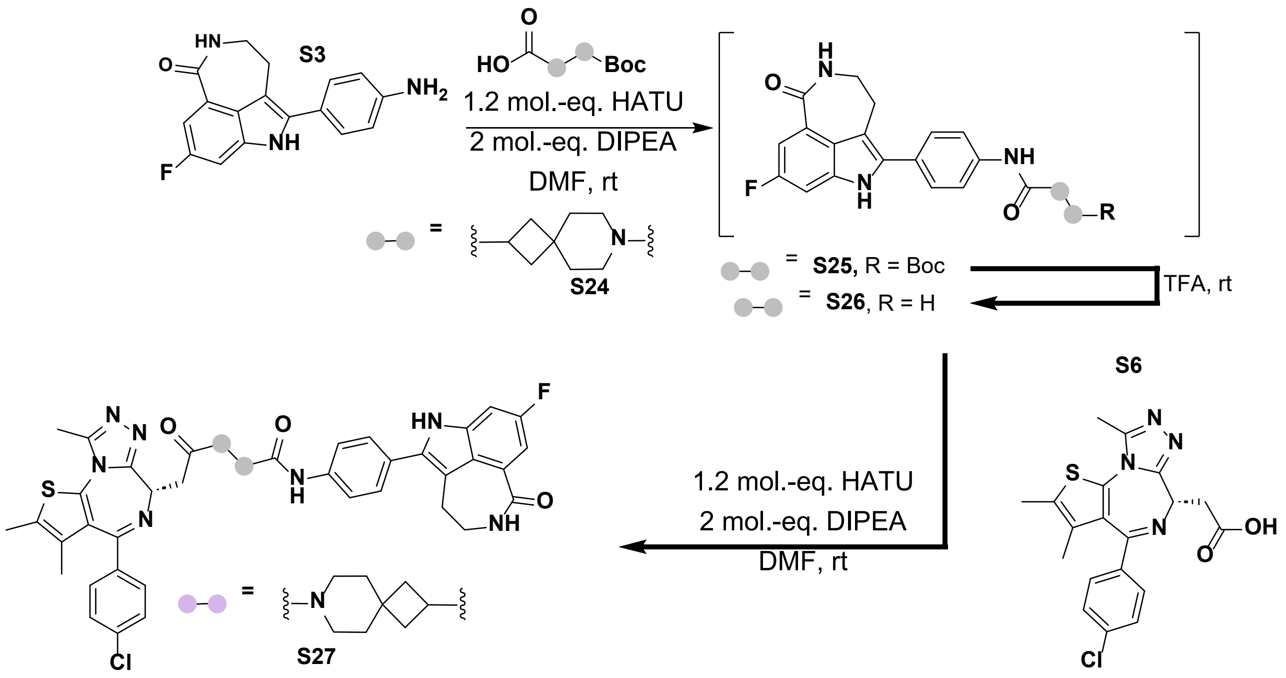

Synthesis of tert-butyl 2-((4-(8-fluoro-1-oxo-2,3,4,6-tetrahydro-1H-azepino[5,4,3-cd]indol-5-yl)phenyl)carbamoyl)-7-azaspiro[3.5]nonane-7-carboxylate (**S25**): A 3.0 mL dram vial equipped with a stir bar was charged with 7-(tert-butoxycarbonyl)-7-azaspiro[3.5]nonane-2-carboxylic acid (**S24**) (55 mg, 0.2 mmol), HATU (93 mg, 0.24 mmol), N,N-dimethylformamide (1.0 mL), and N,N-diisopropylethylamine (71 µL, 0.41 mmol). The solution was allowed to stir for 10 minutes before the portion wise addition of 5-(4-aminophenyl)-8-fluoro-2,3,4,6-tetrahydro-1H-azepino[5,4,3-cd]indol-1-one (**S3**) (60 mg, 0.2 mmol). Stirring continued until UPLC analysis demonstrated complete consumption of the starting materials. The reaction reached completion after 20 minutes of stirring at room temperature. The reaction solution was extracted with ethyl acetate (3 x 10 mL), the organic layer was then washed with brine (3 x 1.0 mL) and dried over sodium sulfate. The organic layer was then iteratively removed in a 20 mL dram vial using reduced pressure to yield a brown oil which was used without further purification.

UPLC-MS, ESI$^+$, *m/z* 547.36 $[M+H]^+$

Synthesis of N-(4-(8-fluoro-1-oxo-2,3,4,6-tetrahydro-1H-azepino[5,4,3-cd]indol-5-yl)phenyl)-7-azaspiro[3.5]nonane-2-carboxamide (**S26**): Trifluoroacetic acid (2 mL) and a stir bar was added to the dram vial containing the crude material **S25**. The reaction progression was monitored using UPLC. After one hour the starting material had been completely converted to the corresponding product and the volatiles were removed using reduced pressure to yield a brown oil.

UPLC-MS, $ESI^+$, *m/z* 447.30 $[M+H]^+$

Synthesis of (*S*)-7-(2-(4-(4-chlorophenyl)-2,3,9-trimethyl-6H-thieno[3,2-f][1,2,4]triazolo[4,3-a][1,4]diazepin-6-yl)acetyl)-N-(4-(8-fluoro-1-oxo-2,3,4,6-tetrahydro-1H-azepino[5,4,3-cd]indol-5-yl)phenyl)-7-azaspiro[3.5]nonane-2-carboxamide (**S27,** compound **5**). A 3.0 mL dram vial equipped with a stir bar was charged with (*S*)-4-(4-chlorophenyl)-2,3,9-trimethyl-6H-thieno[3,2-f][1,2,4]triazolo[4,3-a][1,4]diazepine-6-carboxylic acid (**S6**) (80 mg, 0.2 mmol), HATU (90 mg, 0.24 mmol), N,N-dimethylformamide (1.0 mL), and N,N-diisopropylethylamine (200 µL, 1.0 mmol). The solution was allowed to stir for 10 minutes before being syringed into the dram vial containing the crude material containing **S26**. Stirring continued until UPLC analysis demonstrated complete consumption of the starting materials. The reaction stalled at 50% conversion after one hour until addition of more N,N-diisopropylethylamine (200 µL, 1.0 mmol), after stirring for five additional minutes the starting material was completely consumed, as determined by UPLC analysis. Promptly, the crude reaction solution was directly loaded onto a 12-gram RediSep C18 reversed phase column equipped to a Combiflash NextGen 300+ auto column. The crude reaction mixture was purified using an acidic gradient (0.1% trifluoracetic acid) ranging from 10-100% water/acetonitrile. The gradient began during the first fraction and concluded after the 25-minute run. Clean fractions were determined using UPLC technologies and subsequently lyophilized to yield a white powder (10 mg, 6%).

$^1$H NMR (600 MHz, DMSO-$d_6$) δ 11.64 (s, 1H), 9.99 (s, 1H), 8.25 (t, *J* = 5.9 Hz, 1H), 7.78 (d, *J* = 6.4 Hz, 2H), 7.57 (d, *J* = 7.1 Hz, 2H), 7.50 (d, *J* = 6.6 Hz, 2H), 7.45 – 7.40 (m, 3H), 7.31 (dd, *J* = 9.1, 2.5 Hz, 1H), 4.64 – 4.55 (m, 1H), 3.62 (d, *J* = 7.0 Hz, 2H), 3.56 – 3.50 (m, 1H), 3.49 – 3.45 (m, 1H), 3.42 – 3.35 (m, 4H), 3.27 (s, 1H), 3.03 (s, 2H), 2.61 (s, 3H), 2.42 (s, 3H), 2.14 – 1.91 (m, 5H), 1.79 – 1.70 (m, 1H), 1.63 (s, 4H), 1.58 – 1.55 (m, 1H), 1.50 – 1.45 (m, 1H).

$^{13}$C NMR (151 MHz, DMSO-$d_6$) δ 172.84, 167.87, 167.25, 162.33, 158.44, 156.89, 154.68, 149.23, 138.41, 136.11, 134.63, 131.56, 130.18, 129.59, 129.31, 127.89, 127.67, 125.61, 124.97, 122.70, 118.52, 115.95, 114.03, 110.53, 108.88, 99.96, 53.60, 41.66, 41.26, 40.85, 37.77, 37.57, 37.14, 36.03, 35.42, 34.09, 33.55, 33.38, 32.75, 28.18, 13.43, 12.10, 10.68.

$^{19}$F NMR (376 MHz, DMSO-$d_6$, referenced to $C_6F_6$): δ -123.90.

HRMS (ESI-TOF) m/z: $[M+H]^+$ calculated for $C_{45}H_{43}ClFN_8O_3S$: 829.2846, found: 829.2844.

UPLC-MS, ESI$^+$, *m/z* 829.46 $[M+H]^+$

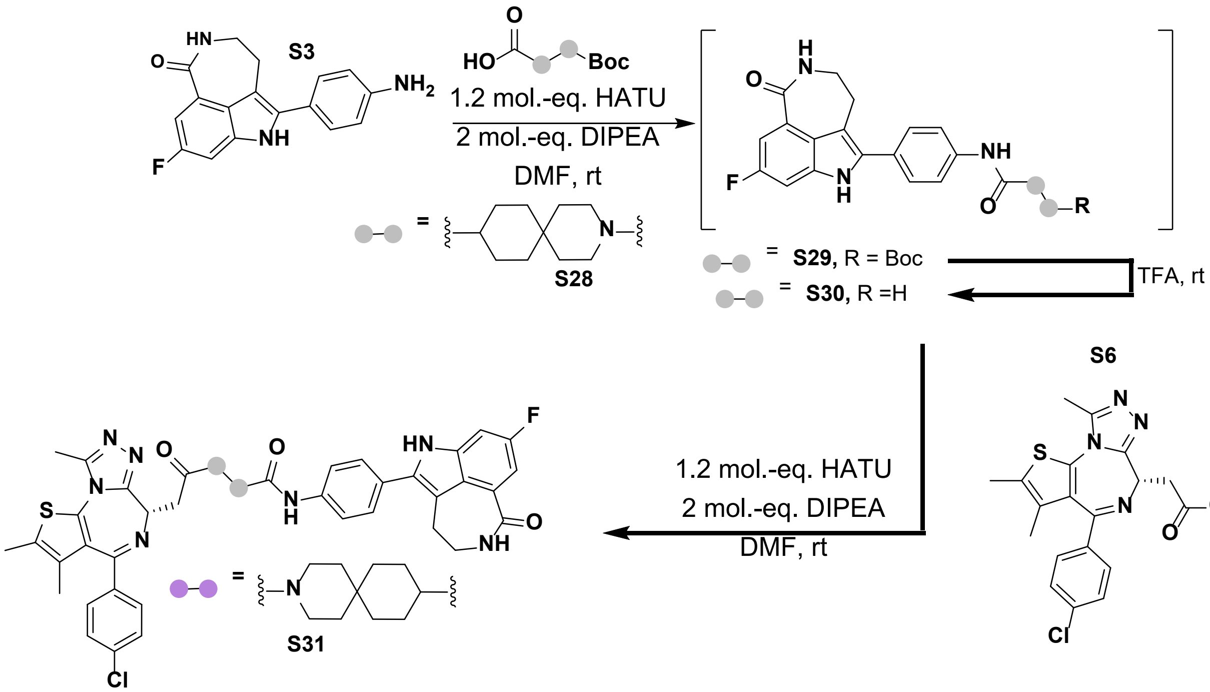

Synthesis of tert-butyl 9-((4-(8-fluoro-1-oxo-2,3,4,6-tetrahydro-1H-azepino[5,4,3-cd]indol-5-yl)phenyl)carbamoyl)-3-azaspiro[5.5]undecane-3-carboxylate (**S29**): A 3.0 mL dram vial equipped with a stir bar was charged with 3-(tert-butoxycarbonyl)-3-azaspiro[5.5]undecane-9-carboxylic acid (**S28**) (60 mg, 0.2 mmol), HATU (93 mg, 0.24 mmol), N,N-dimethylformamide (1.0 mL), and N,N-diisopropylethylamine (71 µL, 0.41 mmol). The solution was allowed to stir for 10 minutes before the portion wise addition of 5-(4-aminophenyl)-8-fluoro-2,3,4,6-tetrahydro-1H-azepino[5,4,3-cd]indol-1-one (**S3**) (60 mg, 0.2 mmol). Stirring continued until UPLC analysis demonstrated complete consumption of the starting materials. The reaction reached completion after 20 minutes of stirring at room temperature. The reaction solution was extracted with ethyl acetate (3 x 10 mL), the organic layer was then washed with brine (3 x 1.0 mL) and dried over sodium sulfate. The organic layer was then iteratively removed in a 20 mL dram vial using reduced pressure to yield a brown oil which was used without further purification.

UPLC-MS, ESI$^+$, *m/z* 575.43 $[M]^+$

Synthesis of N-(4-(8-fluoro-1-oxo-2,3,4,6-tetrahydro-1H-azepino[5,4,3-cd]indol-5-yl)phenyl)-3-azaspiro[5.5]undecane-9-carboxamide (**30**): Trifluoroacetic acid (2 mL) and a stir bar was added to the dram vial containing the crude material **S29**. The reaction progression was monitored using UPLC. After one hour the starting material had been completely converted to the corresponding product and the volatiles were removed using reduced pressure to yield a brown oil.

UPLC-MS, ESI$^+$, *m/z* 475.37 $[M+H]^+$

Synthesis of (*S*)-3-(2-(4-(4-chlorophenyl)-2,3,9-trimethyl-6H-thieno[3,2-f][1,2,4]triazolo[4,3-a][1,4]diazepin-6-yl)acetyl)-N-(4-(8-fluoro-1-oxo-2,3,4,6-tetrahydro-1H-azepino[5,4,3-cd]indol-5-yl)phenyl)-3-azaspiro[5.5]undecane-9-carboxamide (**S31**, compound **6**): A 3.0 mL dram vial equipped with a stir bar was charged with (*S*)-4-(4-chlorophenyl)-2,3,9-trimethyl-6H-thieno[3,2-f][1,2,4]triazolo[4,3-a][1,4]diazepine-6-carboxylic acid (**S6**) (80 mg, 0.2 mmol), HATU (90 mg, 0.24 mmol), N,N-dimethylformamide (1.0 mL), and N,N-diisopropylethylamine (200 µL, 1.0 mmol). The solution was allowed to stir for 10 minutes before being syringed into the dram vial containing the crude material **S30**. Stirring continued until UPLC analysis demonstrated complete consumption of the starting materials. The reaction stalled at 50% conversion after one hour until addition of more N,N-diisopropylethylamine (200 µL, 1.0 mmol), after stirring for five additional minutes the starting material was completely consumed, as determined by UPLC analysis. Promptly, the crude reaction solution was directly loaded onto a 12-gram RediSep C18 reversed phase column equipped to a Combiflash NextGen 300+ auto column. The crude reaction mixture was purified using an acidic gradient (0.1% trifluoracetic acid) ranging from 10-100% water/acetonitrile. The gradient began during the first fraction and concluded after the 25-minute run. Clean fractions were determined using UPLC technologies and subsequently lyophilized to yield a white powder (19 mg, 10%).

$^1$H NMR (600 MHz, DMSO-$d_6$) δ 11.63 (d, *J* = 2.1 Hz, 1H), 10.02 (d, *J* = 5.4 Hz, 1H), 8.25 (t, *J* = 5.7 Hz, 1H), 7.77 (dd, *J* = 8.7, 3.5 Hz, 2H), 7.57 (dd, *J* = 8.8, 2.6 Hz, 2H), 7.50 (d, *J* = 8.9 Hz, 2H), 7.45 (dd, *J* = 8.7, 3.4 Hz, 2H), 7.41 (dd, *J* = 11.0, 3.1 Hz, 1H), 7.30 (dd, *J* = 8.6, 3.0 Hz, 1H), 4.60 (t, *J* = 6.7 Hz, 1H), 3.66 – 3.58 (m, 3H), 3.50 – 3.36 (m, 5H), 3.03 (s, 2H), 2.61 (s, 3H), 2.42 (s, 3H), 2.40 – 2.33 (m, 1H), 1.82 (d, *J* = 12.3 Hz, 2H), 1.71 – 1.60 (m, 8H), 1.51 – 1.40 (m, 2H), 1.28 (t, *J* = 5.5 Hz, 1H), 1.22 – 1.15 (m, 2H).

$^{13}C$ NMR (151 MHz, DMSO-$d_6$) δ 173.79, 167.86, 167.13, 162.32, 158.44, 156.89, 154.69, 149.26, 138.46, 136.10, 134.64, 131.56, 130.20, 129.61, 129.31, 127.89, 127.64, 125.58, 125.02, 122.69, 118.53, 115.83, 113.90, 110.52, 108.88, 99.96, 53.56, 44.55, 41.26, 40.46, 40.33, 36.62, 36.48, 34.16, 34.11, 34.06, 33.90, 31.09, 30.45, 30.06, 28.18, 23.45, 13.43, 12.10, 10.68.

$^{19}F$ NMR (376 MHz, DMSO-$d_6$, referenced to $C_6F_6$): δ -123.94.

HRMS (ESI-TOF) m/z: $[M+H]^+$ calculated for $C_{47}H_{47}ClFN_8O_3S$: 857.3159, found: 857.3157.

UPLC-MS, $ESI^+$, *m/z* 857.53 $[M+H]^+$

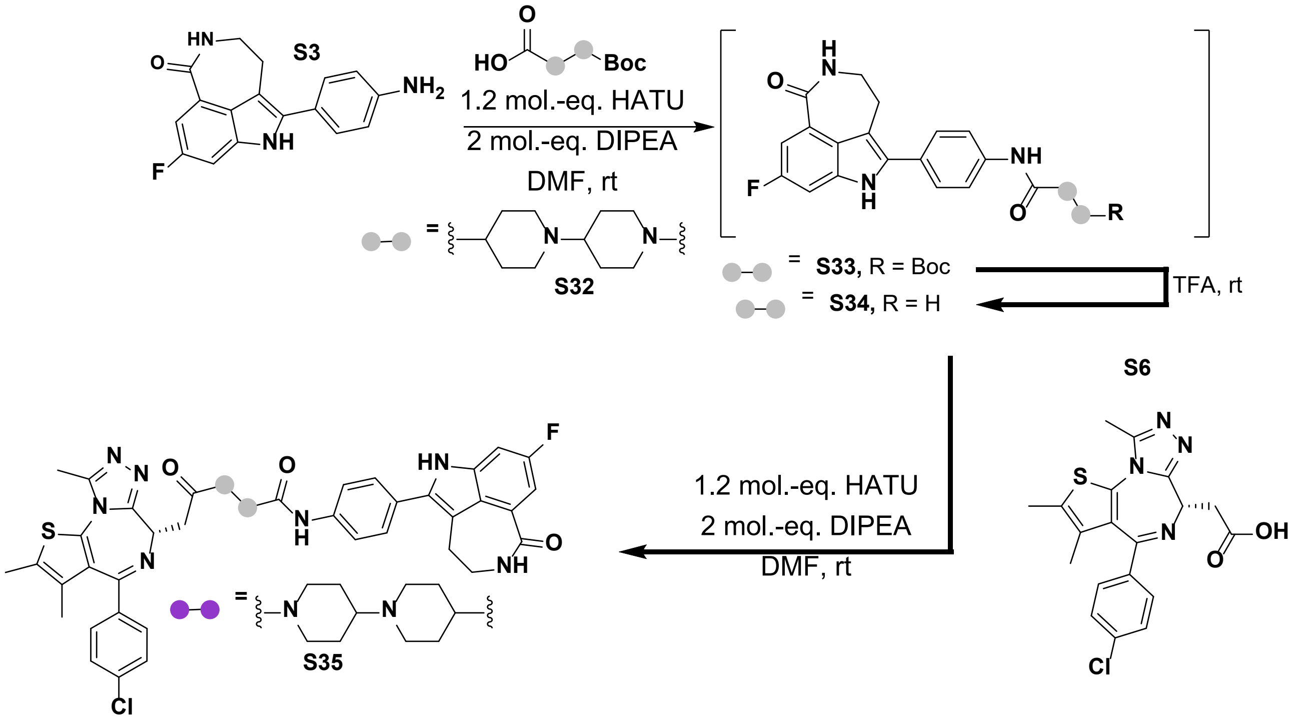

Synthesis of tert-butyl 9-((4-(8-fluoro-1-oxo-2,3,4,6-tetrahydro-1H-azepino[5,4,3-cd]indol-5-yl)phenyl)carbamoyl)-3-azaspiro[5.5]undecane-3-carboxylate (**S33**): A 3.0 mL dram vial equipped with a stir bar was charged with 1'-(tert-butoxycarbonyl)-[1,4'-bipiperidine]-4-carboxylic acid (**S32**) (63 mg, 0.2 mmol), HATU (93 mg, 0.24 mmol), N,N-dimethylformamide (1.0 mL), and N,N-diisopropylethylamine (71 µL, 0.41 mmol). The solution was allowed to stir for 10 minutes before the portion wise addition of 5-(4-aminophenyl)-8-fluoro-2,3,4,6-tetrahydro-1H-azepino[5,4,3-cd]indol-1-one (**S3**) (60 mg, 0.2 mmol). Stirring continued until UPLC analysis demonstrated complete consumption of the starting materials. The reaction reached completion after 20 minutes of stirring at room temperature.

The reaction solution was extracted with ethyl acetate (3 x 10 mL), the organic layer was then washed with brine (3 x 1.0 mL) and dried over sodium sulfate. The organic layer was then iteratively removed in a 20 mL dram vial using reduced pressure to yield a brown oil which was used without further purification.

UPLC-MS, ESI$^+$, *m/z* 595.45 $[M+H]^+$

Synthesis of N-(4-(8-fluoro-1-oxo-2,3,4,6-tetrahydro-1H-azepino[5,4,3-cd]indol-5-yl)phenyl)-[1,4'-bipiperidine]-4-carboxamide (**S34**): Trifluoroacetic acid (2 mL) and a stir bar was added to the dram vial containing the crude product **S33**. The reaction progression was monitored using UPLC. After one hour the starting material had been competely converted to the corresponding product and the volatiles were removed using reduced pressure to yield a brown oil.

UPLC-MS, ESI$^+$, *m/z* 490.35 $[M+H]^+$

Synthesis of ((*S*)-1'-(2-(4-(4-chlorophenyl)-2,3,9-trimethyl-6H-thieno[3,2-f][1,2,4]triazolo[4,3-a][1,4]diazepin-6-yl)acetyl)-N-(4-(8-fluoro-1-oxo-2,3,4,6-tetrahydro-1H-azepino[5,4,3-cd]indol-5-yl)phenyl)-[1,4'-bipiperidine]-4-carboxamide (**S35**, compound **7**): A 3.0 mL dram vial equipped with a stir bar was charged with (*S*)-4-(4-chlorophenyl)-2,3,9-trimethyl-6H-thieno[3,2-f][1,2,4]triazolo[4,3-a][1,4]diazepine-6-carboxylic acid (**S6**) (80 mg, 0.2 mmol), HATU (90 mg, 0.24 mmol), N,N-dimethylformamide (1.0 mL), and N,N-diisopropylethylamine (200 µL, 1.0 mmol). The solution was allowed to stir for 10 minutes before being syringed into the dram vial containing the crude material **S34**. Stirring continued until UPLC analysis demonstrated complete consumption of the starting materials. The reaction stalled at 50% conversion after one hour until addition of more N,N-diisopropylethylamine (200 µL, 1.0 mmol), after stirring for five additional minutes the starting material was completely consumed, as determined by UPLC analysis. Promptly, the crude reaction solution was directly loaded onto a 12-gram RediSep C18 reversed phase column equipped to a Combiflash NextGen 300+ auto column. The crude reaction mixture was purified using an acidic gradient (0.1% trifluoracetic acid) ranging from 10-100% water/acetonitrile. The gradient began during the first fraction and concluded after the 25-minute run. Clean fractions were determined using UPLC technologies and subsequently lyophilized to yield a white powder (10 mg, 6%).

$^1$H NMR (600 MHz, DMSO-$d_6$) δ 11.65 (s, 1H), 10.26 (s, 1H), 9.24 (s, 1H), 8.26 (t, *J* = 5.7 Hz, 1H), 7.77 (d, *J* = 8.4 Hz, 2H), 7.59 (d, *J* = 8.4 Hz, 2H), 7.54 – 7.48 (m, 3H), 7.48 – 7.39 (m, 4H), 7.31 (dd, *J* = 9.1, 2.4 Hz, 1H), 4.59 (q, *J* = 6.2 Hz, 1H), 4.54 (d, *J* = 12.9 Hz, 1H), 4.45 (t, *J* = 7.1 Hz, 1H), 4.37 (t, *J* = 15.0 Hz, 1H), 3.47 – 3.36 (m, 5H), 3.32 (dd, *J* = 16.7, 7.4 Hz, 1H), 3.17 (t, *J* = 12.6 Hz, 1H), 3.05 (br, 4H), 2.62 – 2.60 (m, 4H), 2.42 (d, *J* = 6.2 Hz, 4H), 2.15 (m, 5H), 1.94 (m, 3H), 1.63 (d, *J* = 4.4 Hz, 4H).

$^{13}$C NMR (151 MHz, DMSO-$d_6$) δ 171.26, 167.83, 162.51, 158.44, 156.89, 154.60, 154.20, 149.20, 137.93, 136.11, 134.63, 131.59, 130.14, 129.53, 128.99, 127.88, 127.65, 126.00, 125.04, 122.62, 118.68, 116.71, 114.75, 110.61, 108.91, 99.78, 62.09, 52.93, 51.98, 47.57, 43.00, 42.75, 41.20, 35.88, 34.17, 33.95, 28.15, 25.21, 13.43, 12.06, 10.65.

$^{19}$F NMR (376 MHz, DMSO-$d_6$, referenced to $C_6F_6$): δ -123.78.

HRMS (ESI-TOF) m/z: $[M+H]^+$ calculated for $C_{47}H_{48}ClFN_9O_3S$: 872.3268, found: 872.3266.

UPLC-MS, ESI$^+$, *m/z* 872.52 $[M+H]^+$

**NMR spectra of novel compounds**

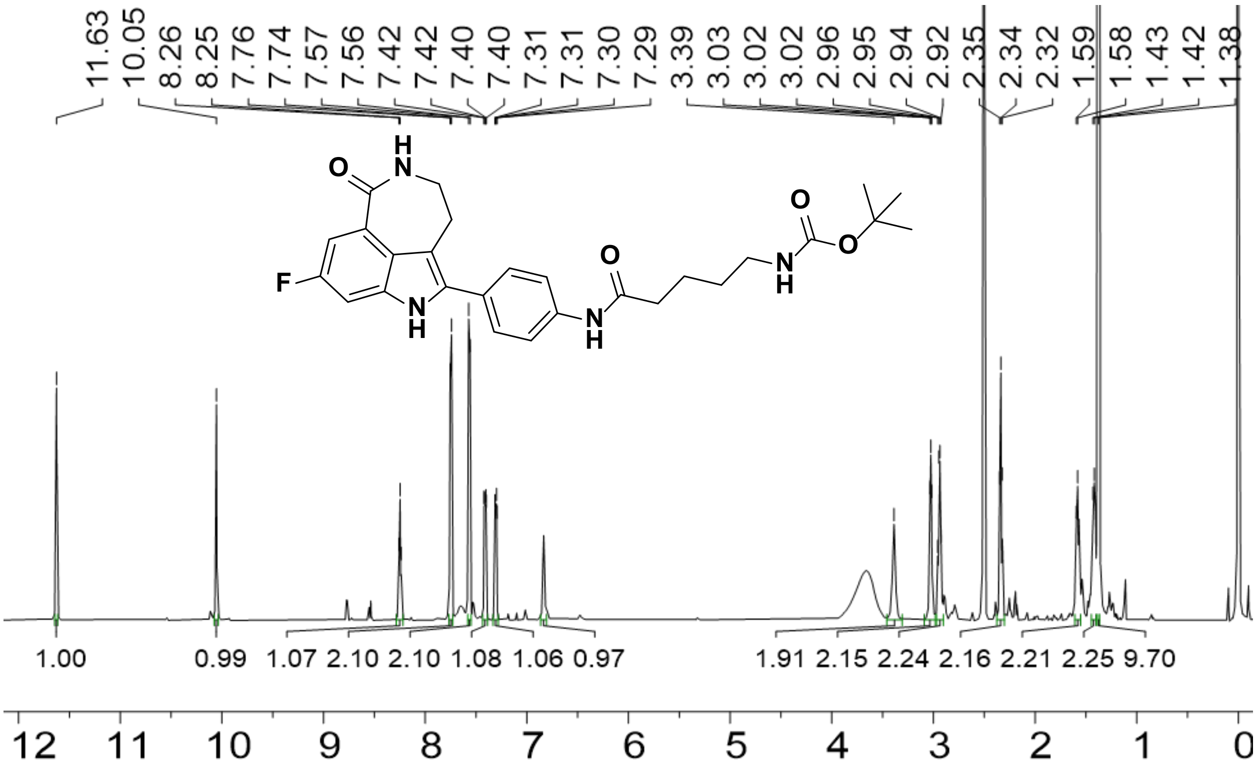

Figure S1. $^{1}H$ NMR spectrum of **S10** (600 MHz, DMSO-$d_6$, 298 K).

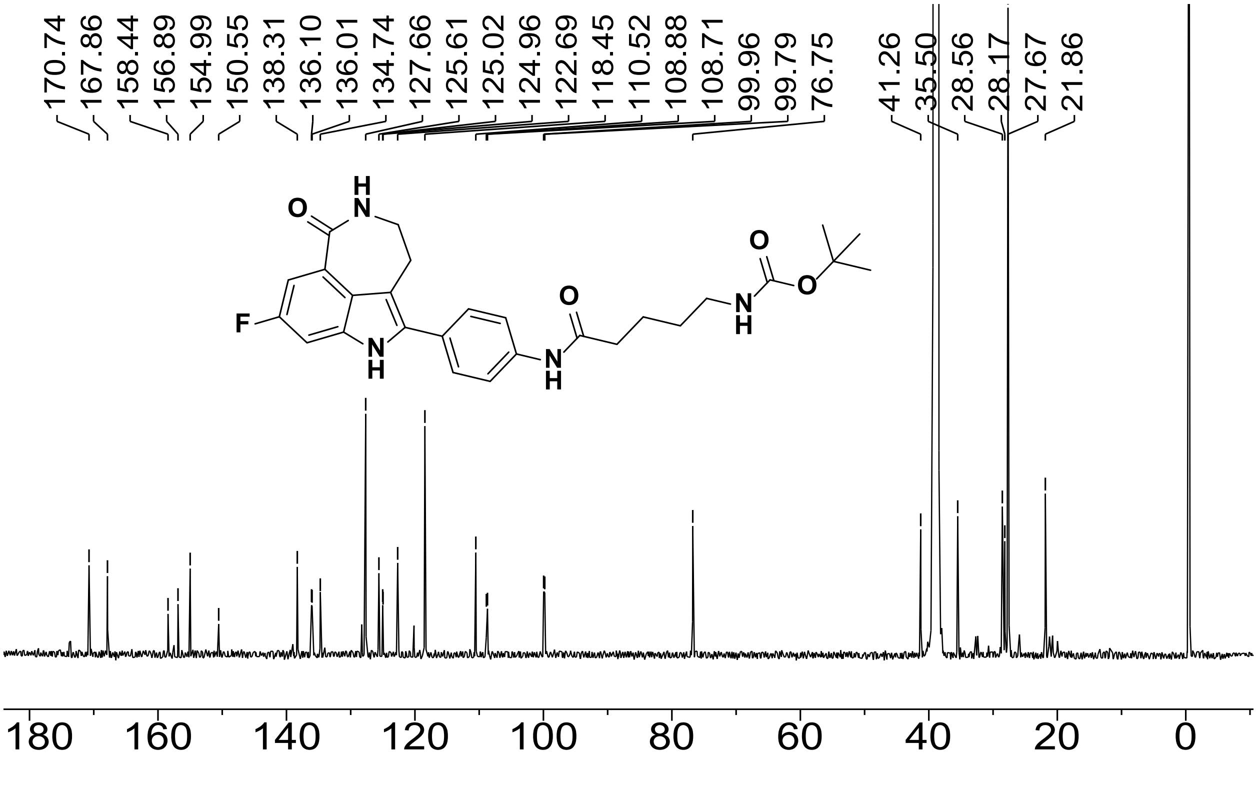

Figure S2. $^{13}C$ NMR spectrum of **S10** (151 MHz, DMSO-$d_6$, 298 K).

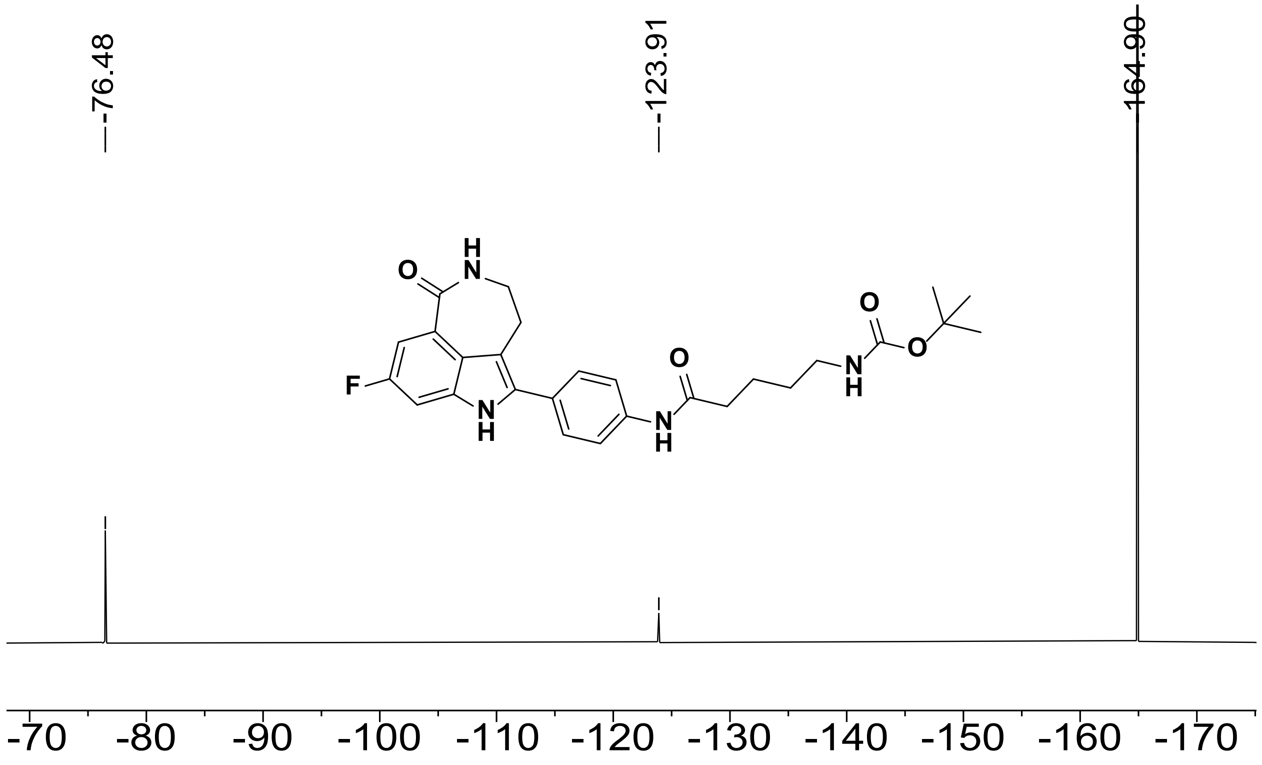

Figure S3. [19]F NMR spectrum of **S10** (376 MHz, DMSO-$d_6$, referenced to $C_6F_6$, 298 K).

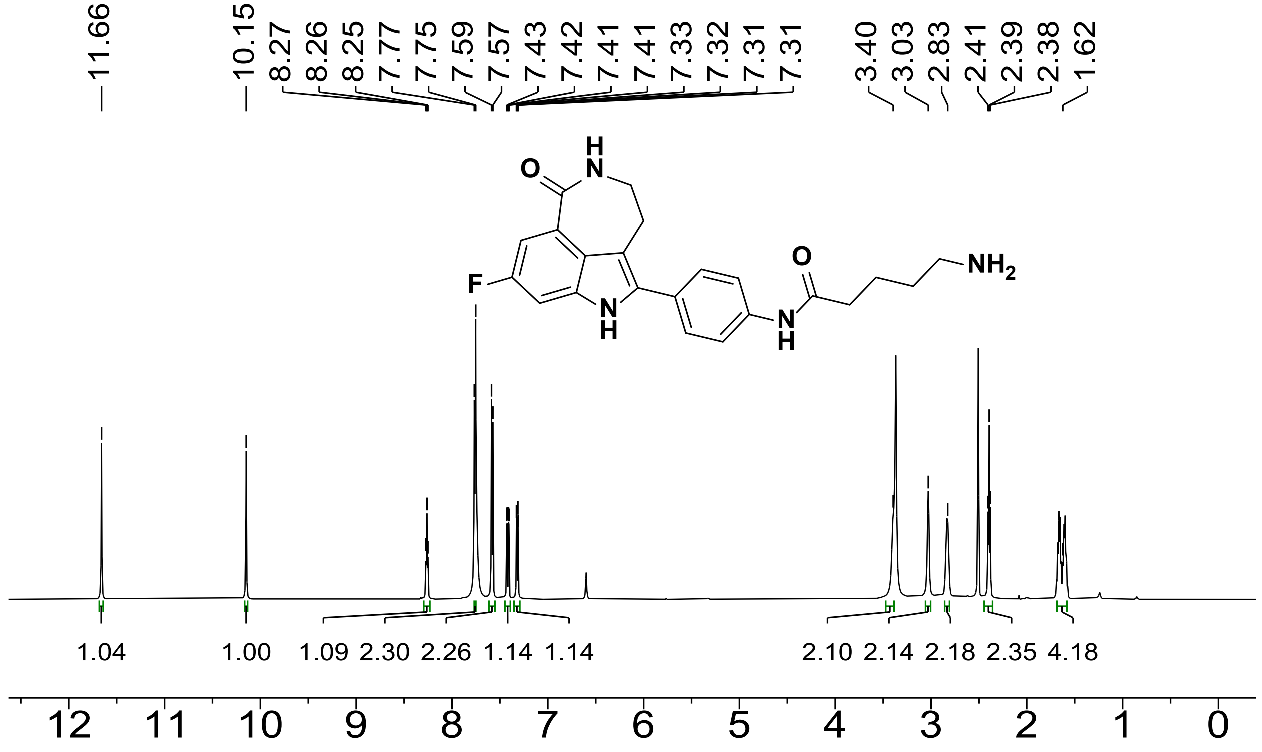

Figure S4. [1]H NMR spectrum of **S12** (600 MHz, DMSO-$d_6$, 298 K).

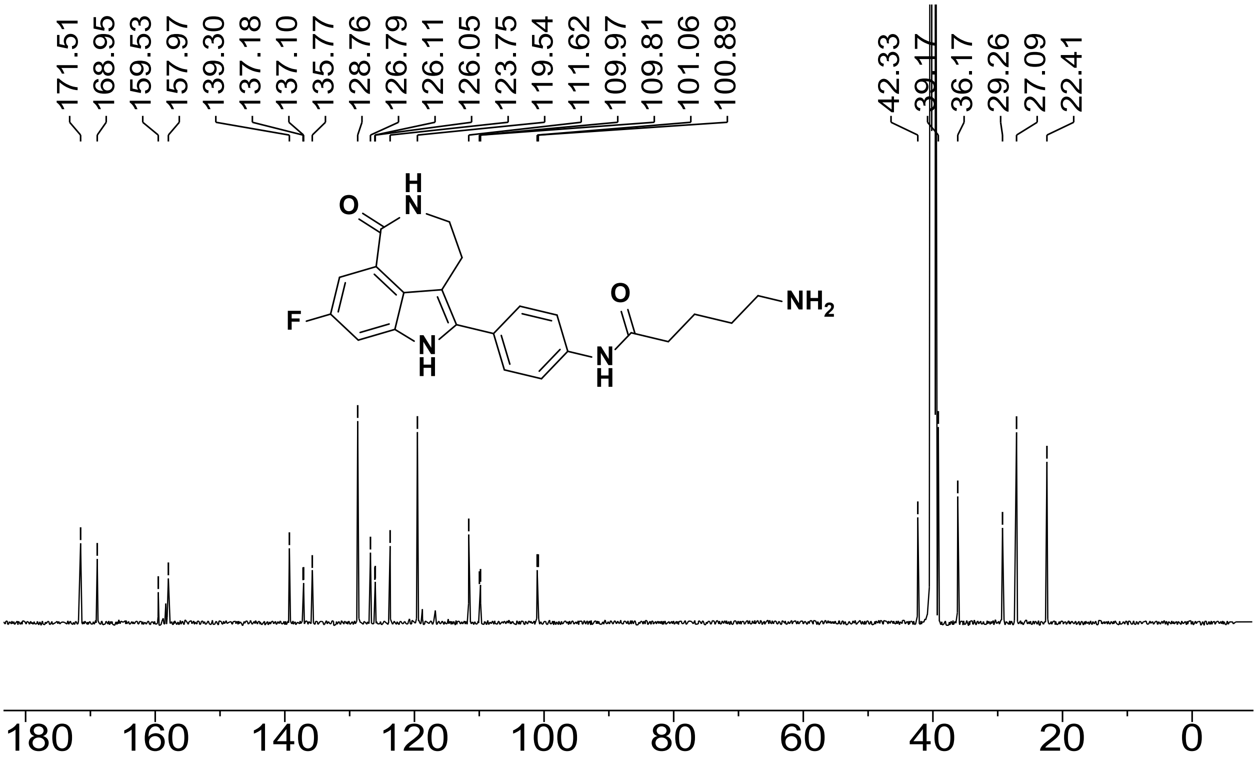

Figure S5. [13]C NMR spectrum of **S12** (151 MHz, DMSO-$d_6$, 298 K).

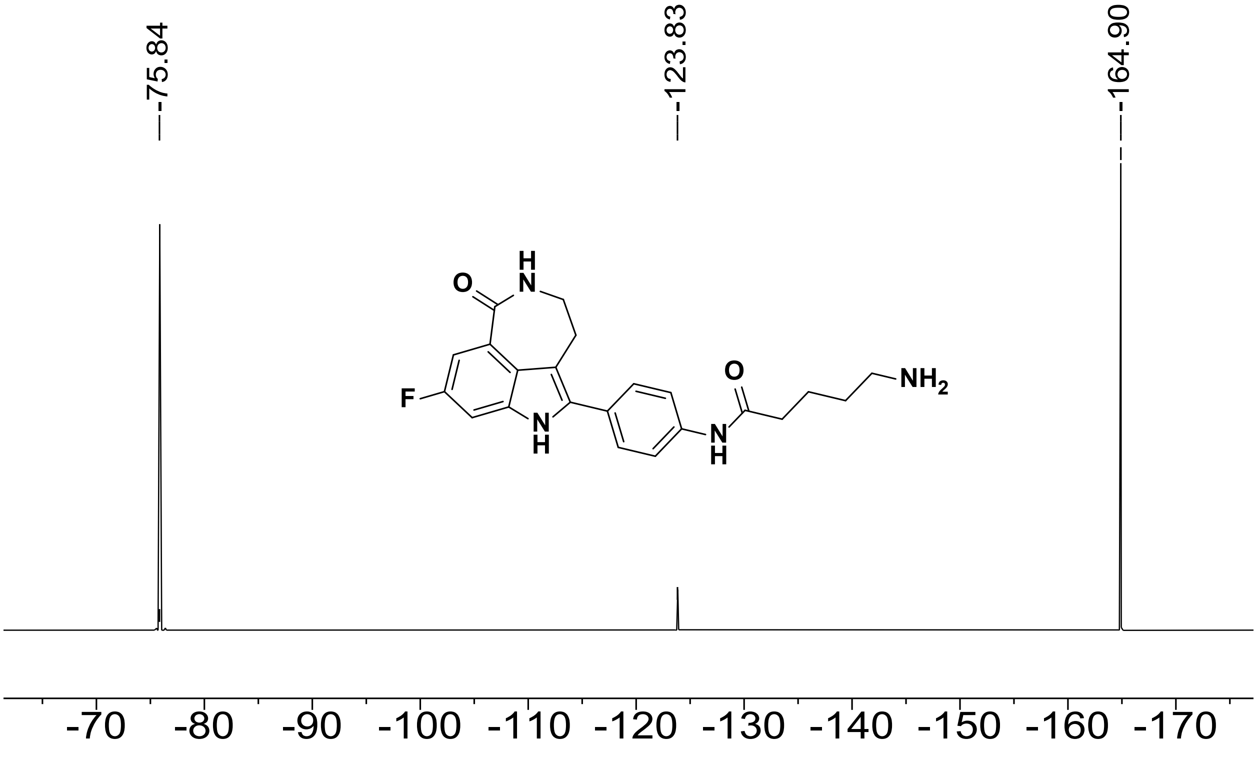

Figure S6. [19]F NMR spectrum of **S12** (376 MHz, DMSO-$d_6$, referenced to $C_6F_6$, 298 K).

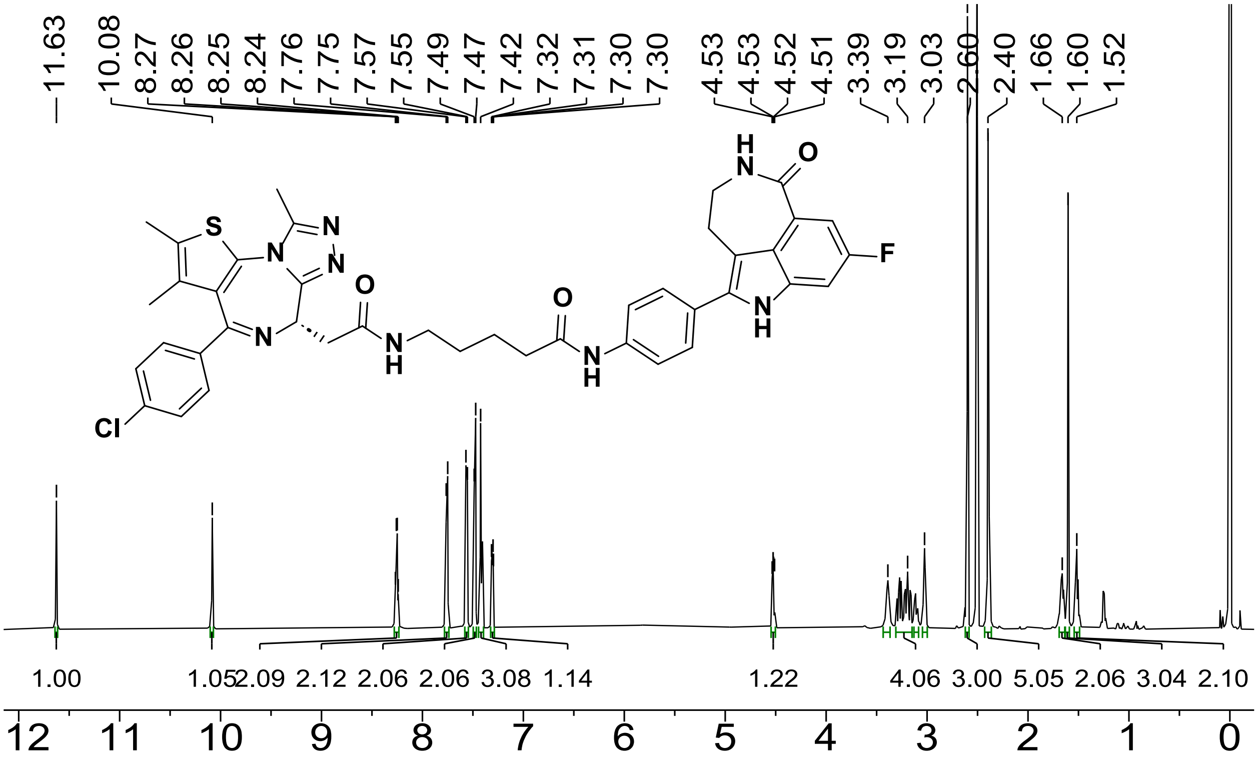

Figure S7. [1]H NMR spectrum of **S14**, compound **1** (600 MHz, DMSO-$d_6$, 298 K).

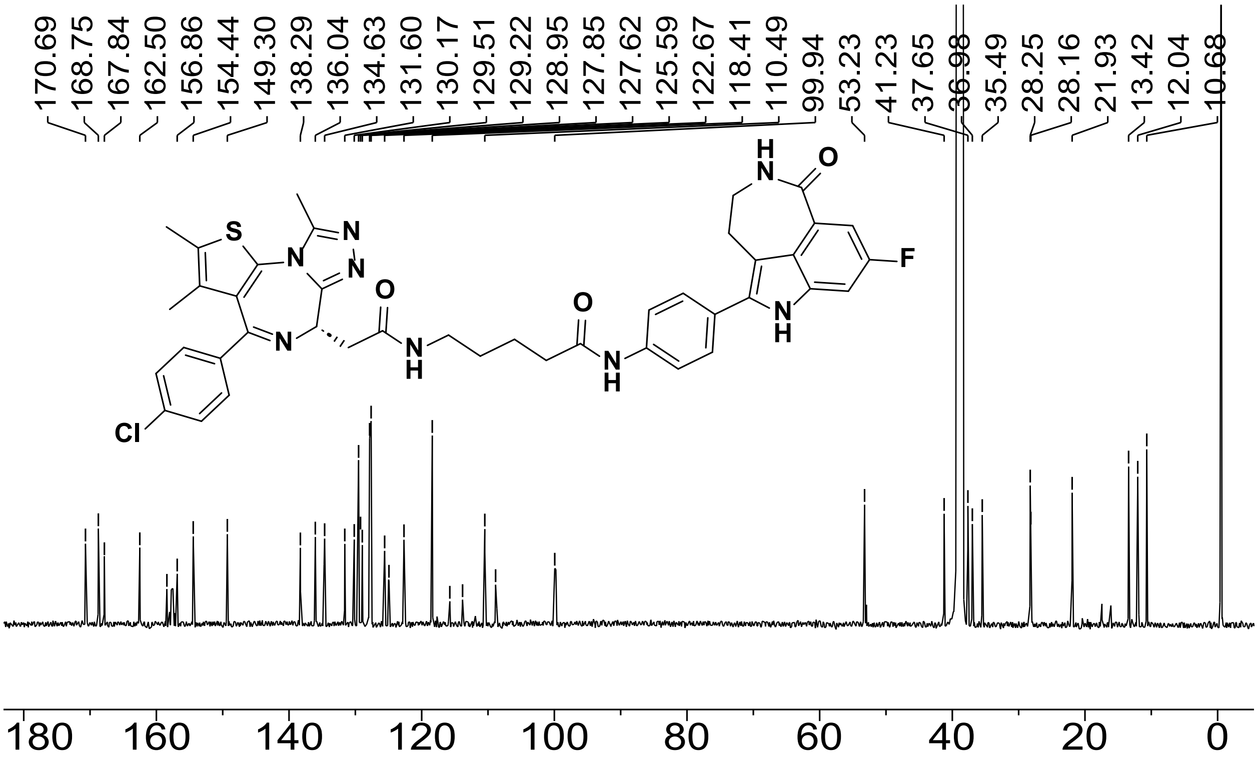

Figure S8. [13]C NMR spectrum of **S14**, compound **1** (151 MHz, DMSO-$d_6$, 298 K).

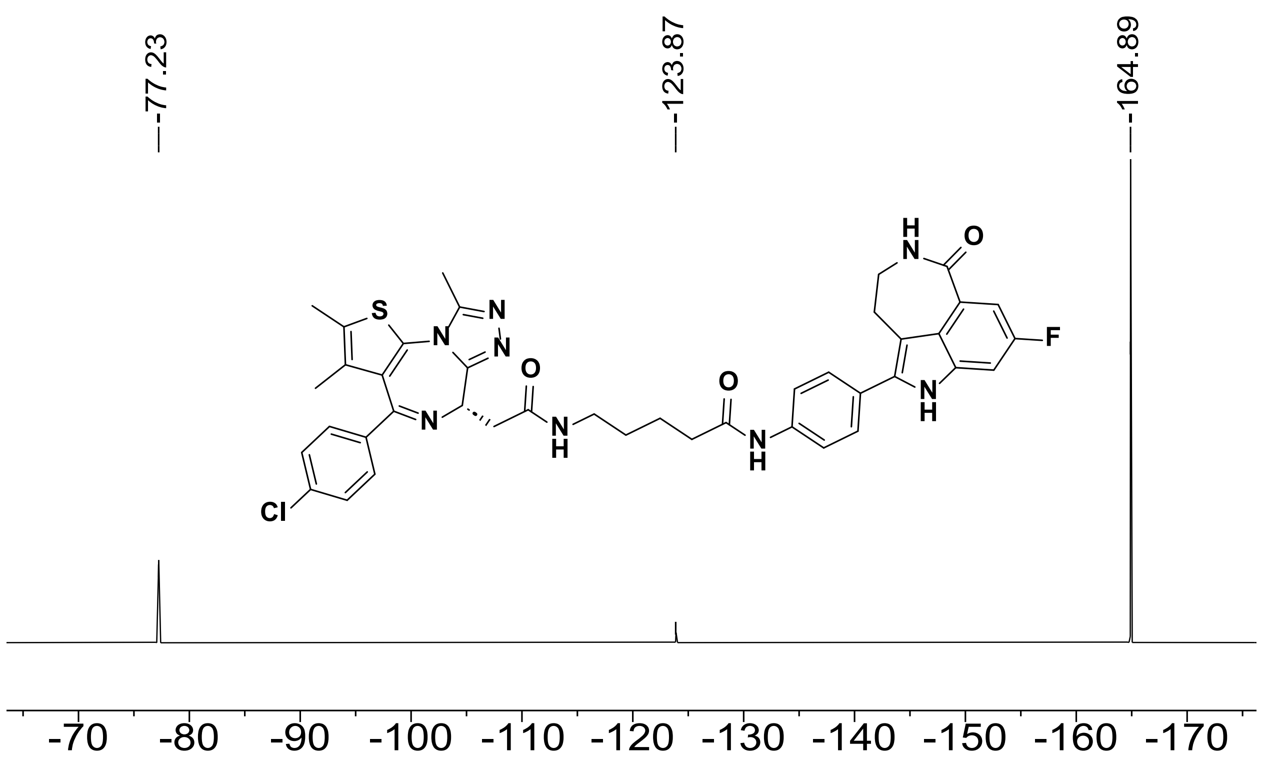

Figure S9. [19]F NMR spectrum of **S14**, compound **1** (376 MHz, DMSO-$d_6$, referenced to $C_6F_6$, 298 K).

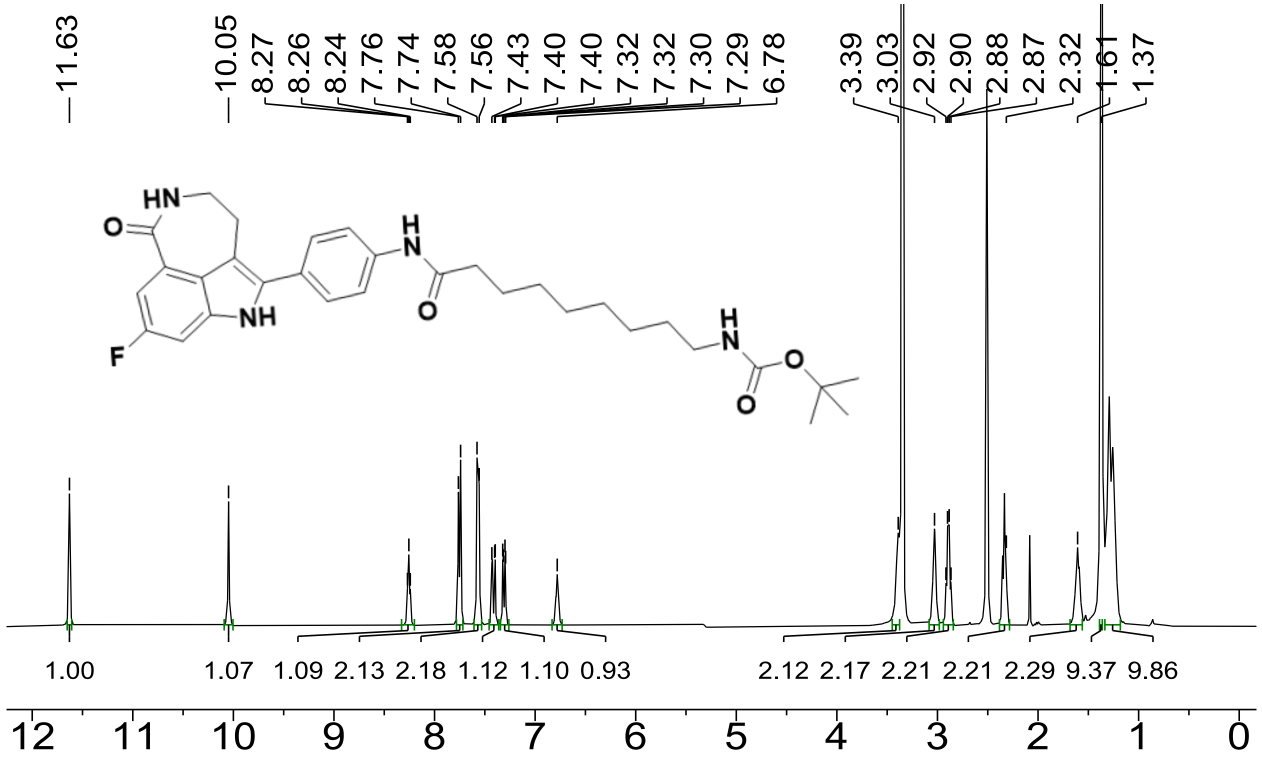

Figure S10. [1]H NMR spectrum of **S11** (600 MHz, DMSO-$d_6$, 298 K).

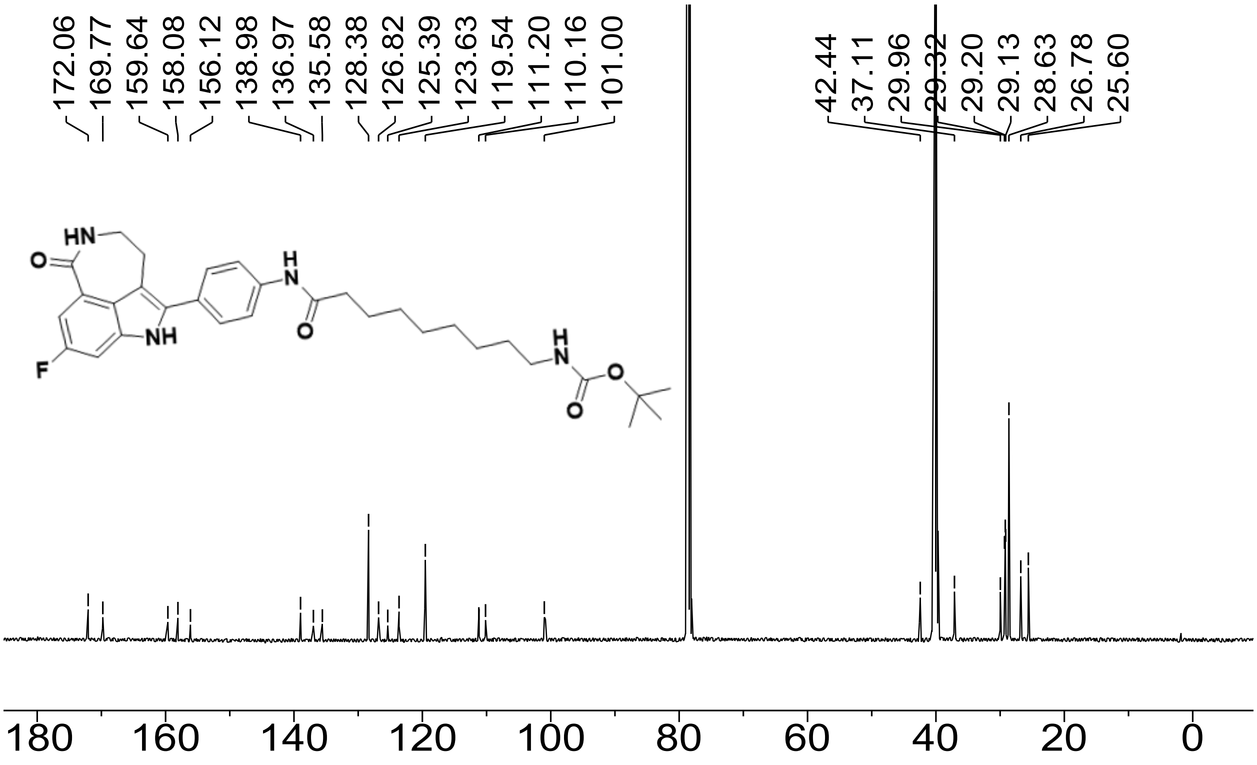

Figure S11. $^{13}C$ NMR spectrum of **S11** (151 MHz, DMSO-*d*$_6$, 298 K).

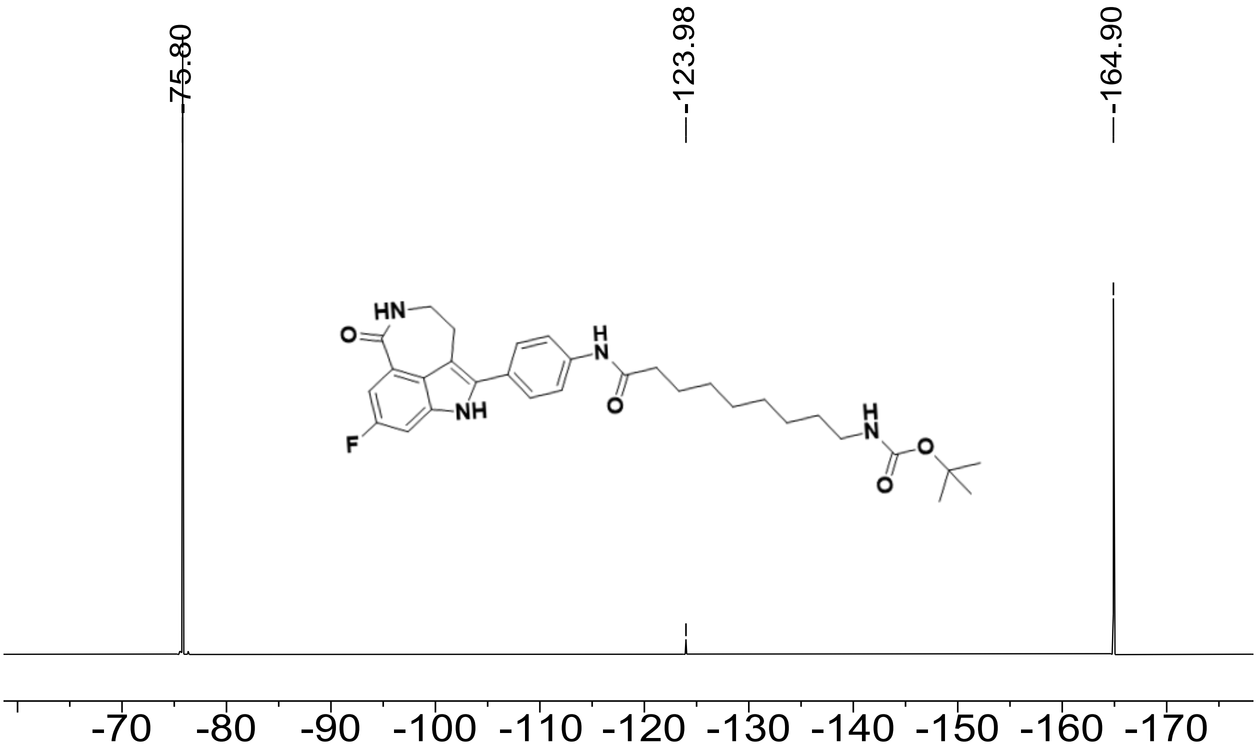

Figure S12. $^{19}F$ NMR spectrum of **S11** (376 MHz, DMSO-*d*$_6$, referenced to $C_6F_6$, 298 K).

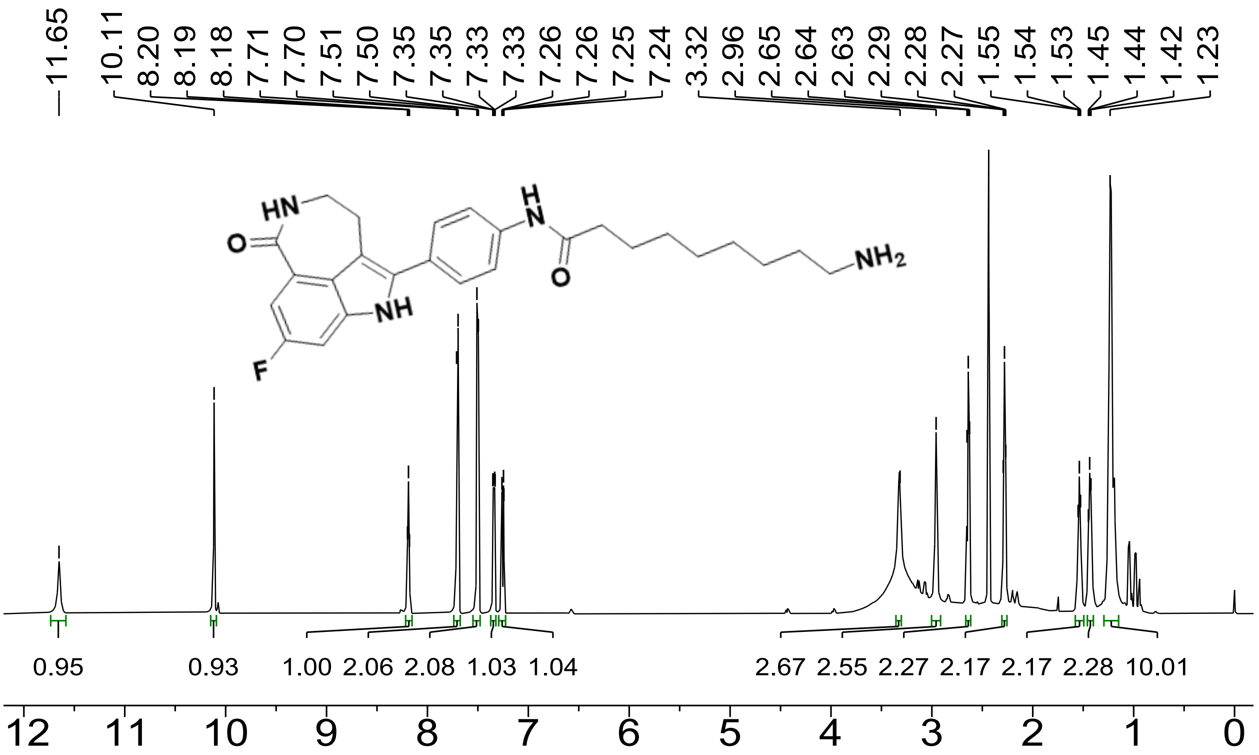

Figure S13. [1]H NMR spectrum of **S13** (600 MHz, DMSO-$d_6$, 298 K).

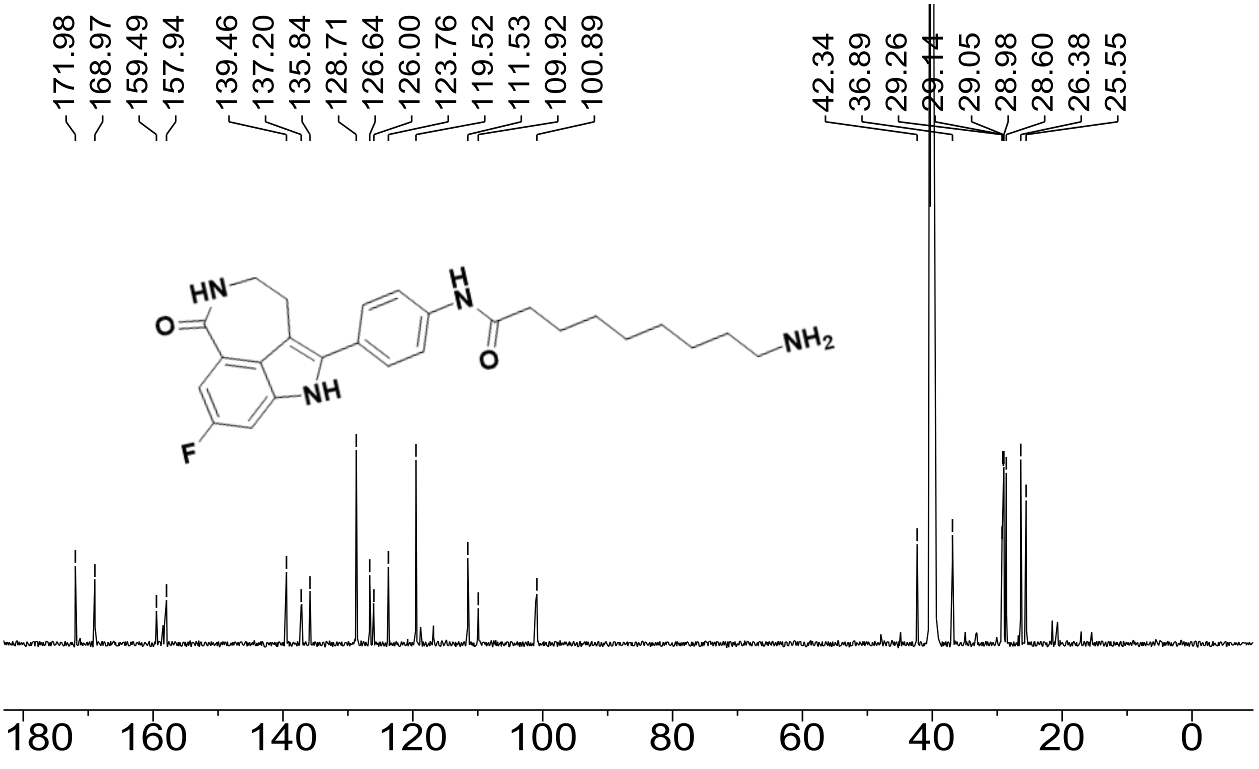

Figure S14. [13]C NMR spectrum of **S13** (151 MHz, DMSO-$d_6$, 298 K).

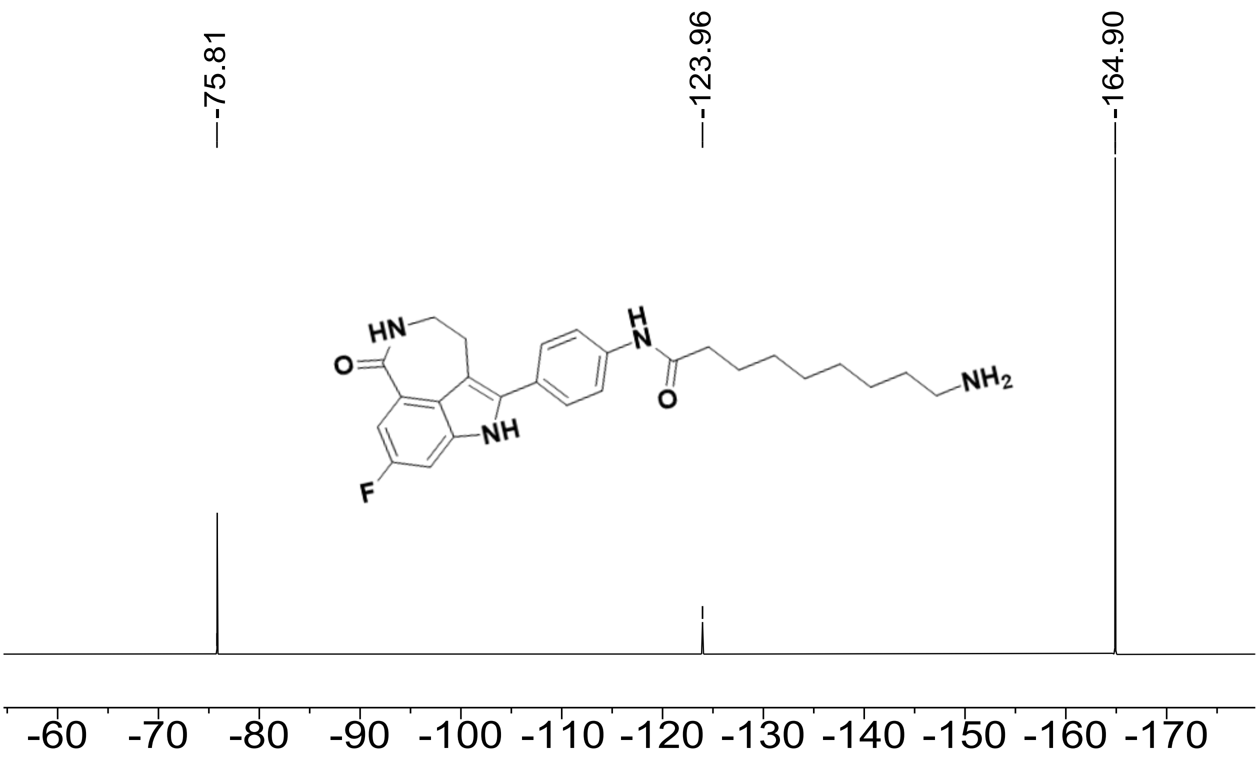

Figure S15. $^{19}$F NMR spectrum of **S13** (376 MHz, DMSO-*d*$_6$, referenced to $C_6F_6$, 298 K).

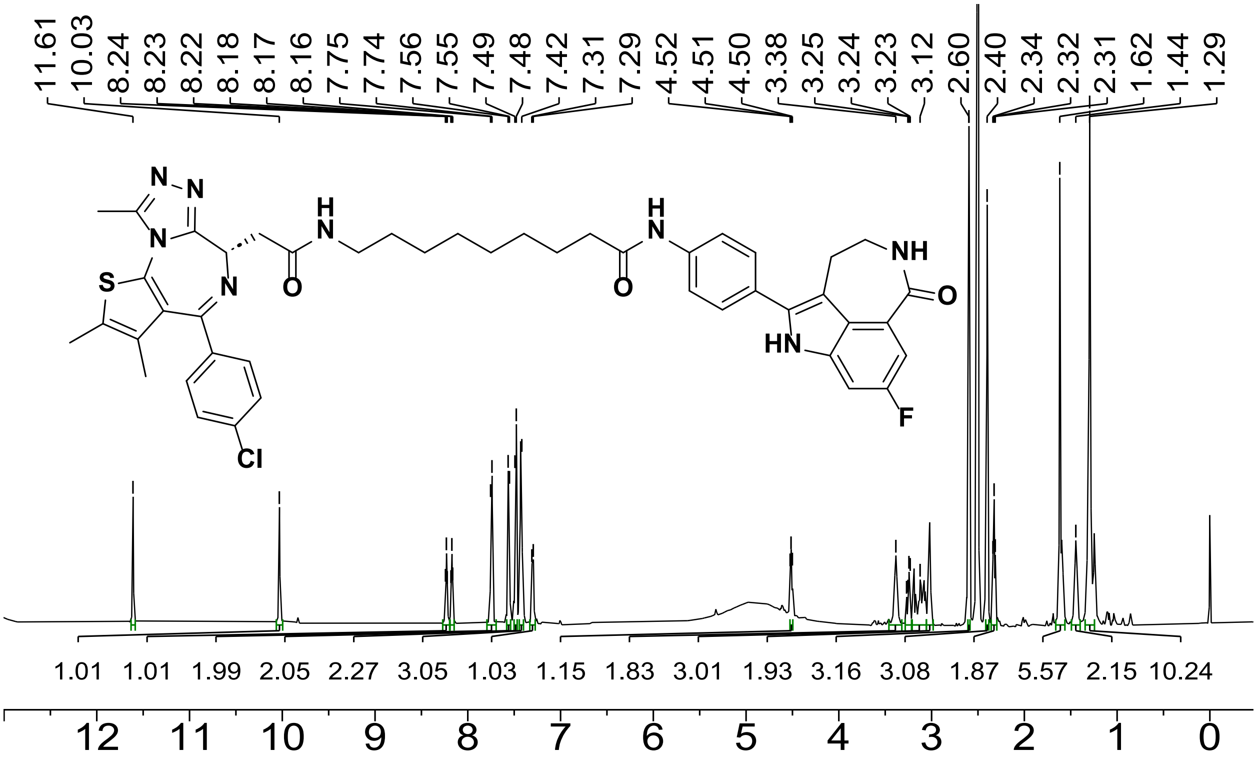

Figure S16. $^{1}$H NMR spectrum of **S15**, compound **2**, PCIP-1 (600 MHz, DMSO-*d*$_6$, 298 K).

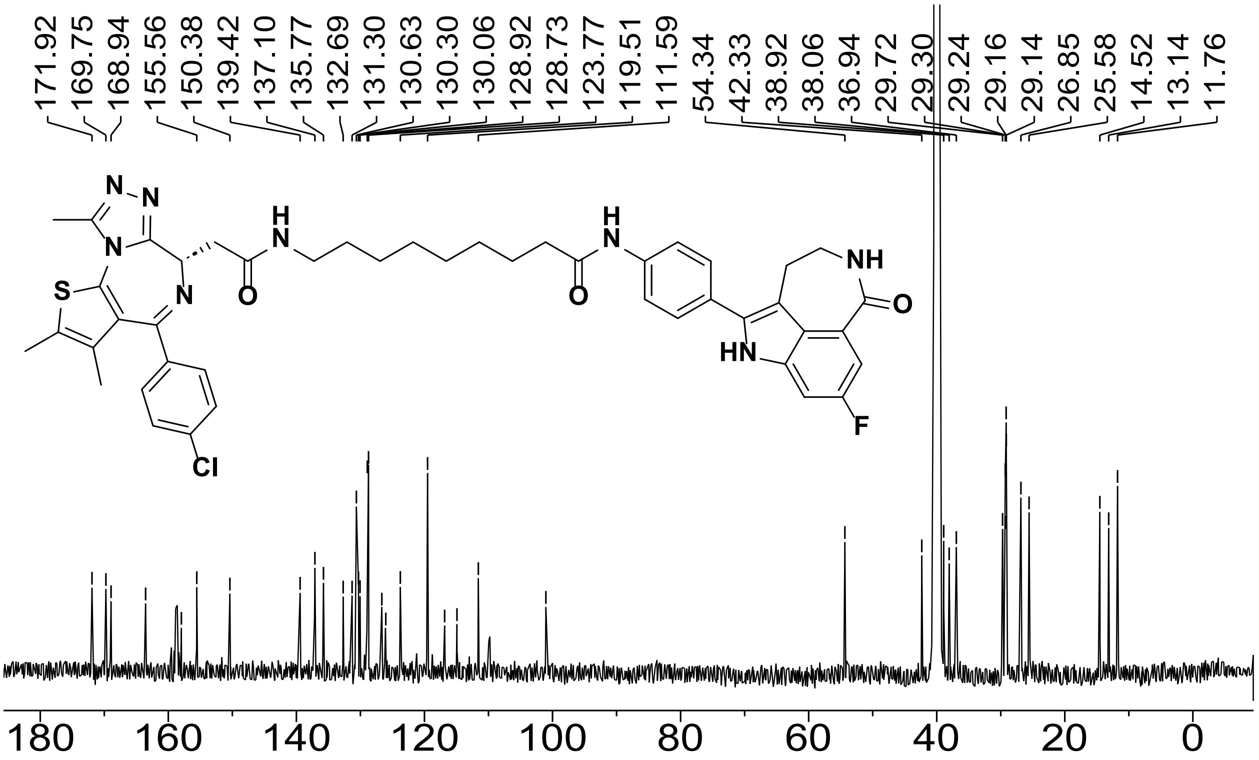

Figure S17. [13]C NMR spectrum of **S15**, compound **2**, PCIP-1 (151 MHz, DMSO-*d*$_6$, 298 K).

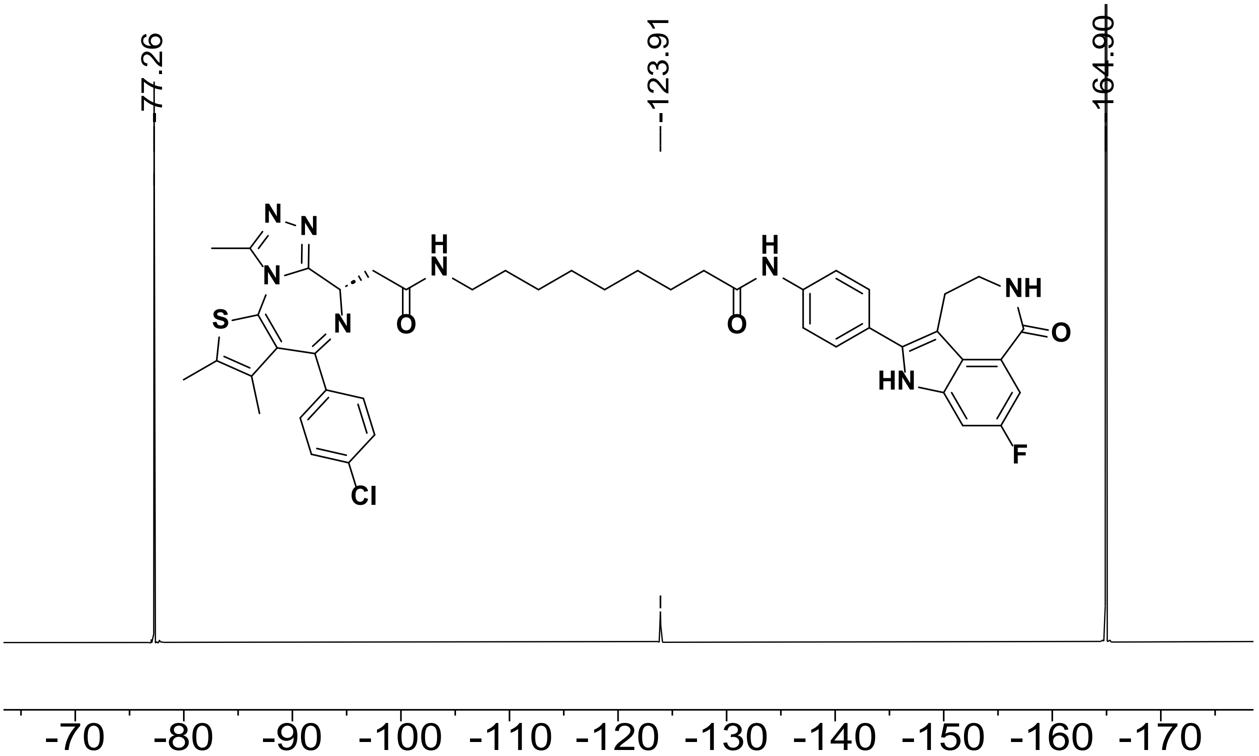

Figure S18. [19]F NMR spectrum of **S15**, compound **2**, PCIP-1 (376 MHz, DMSO-*d*$_6$, referenced to $C_6F_6$, 298 K).

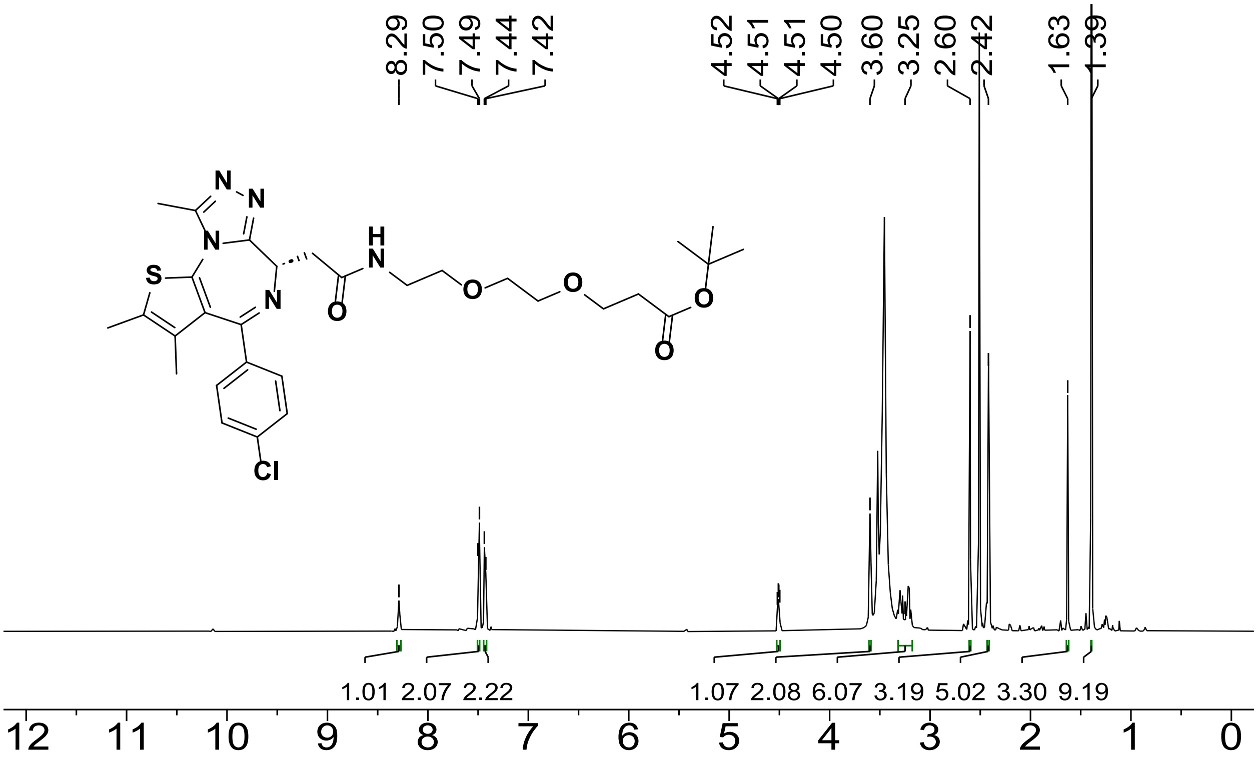

Figure S19. $^{1}$H NMR spectrum of **S18** (600 MHz, DMSO-$d_6$, 298 K).

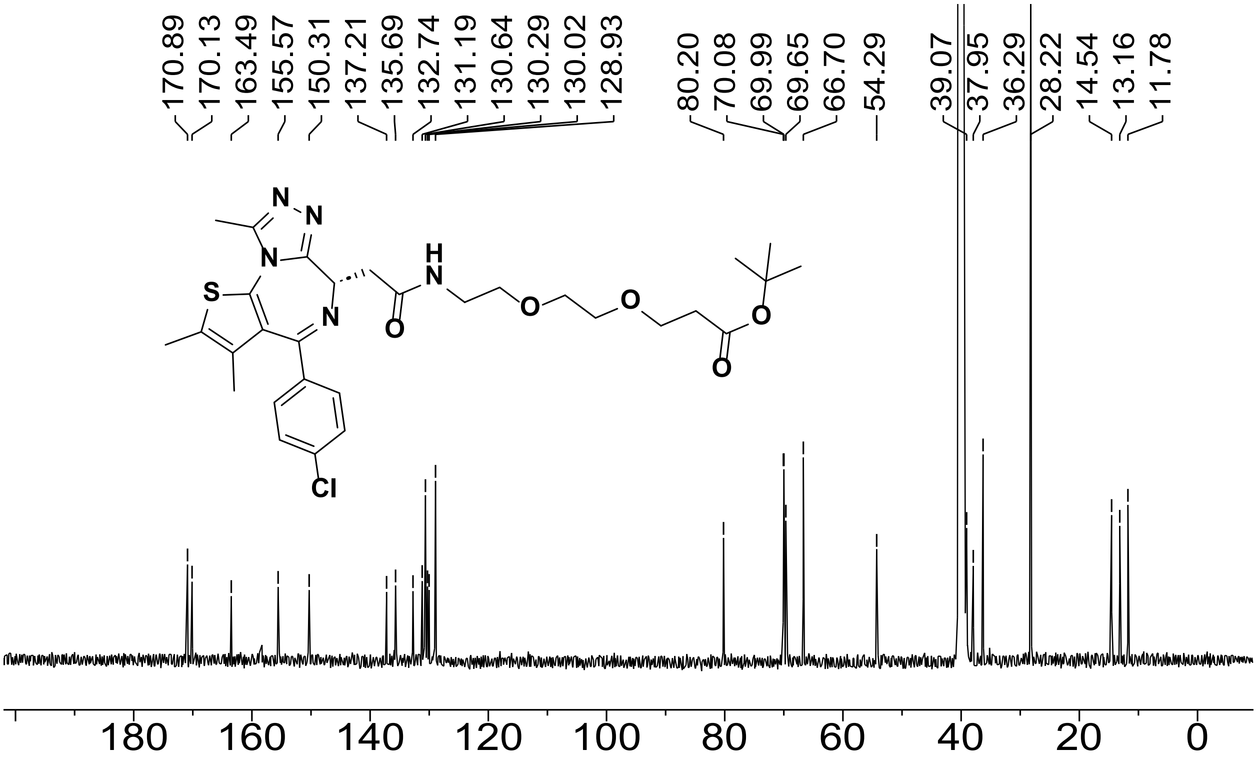

Figure S20. $^{13}$C NMR spectrum of **S18** (151 MHz, DMSO-$d_6$, 298 K).

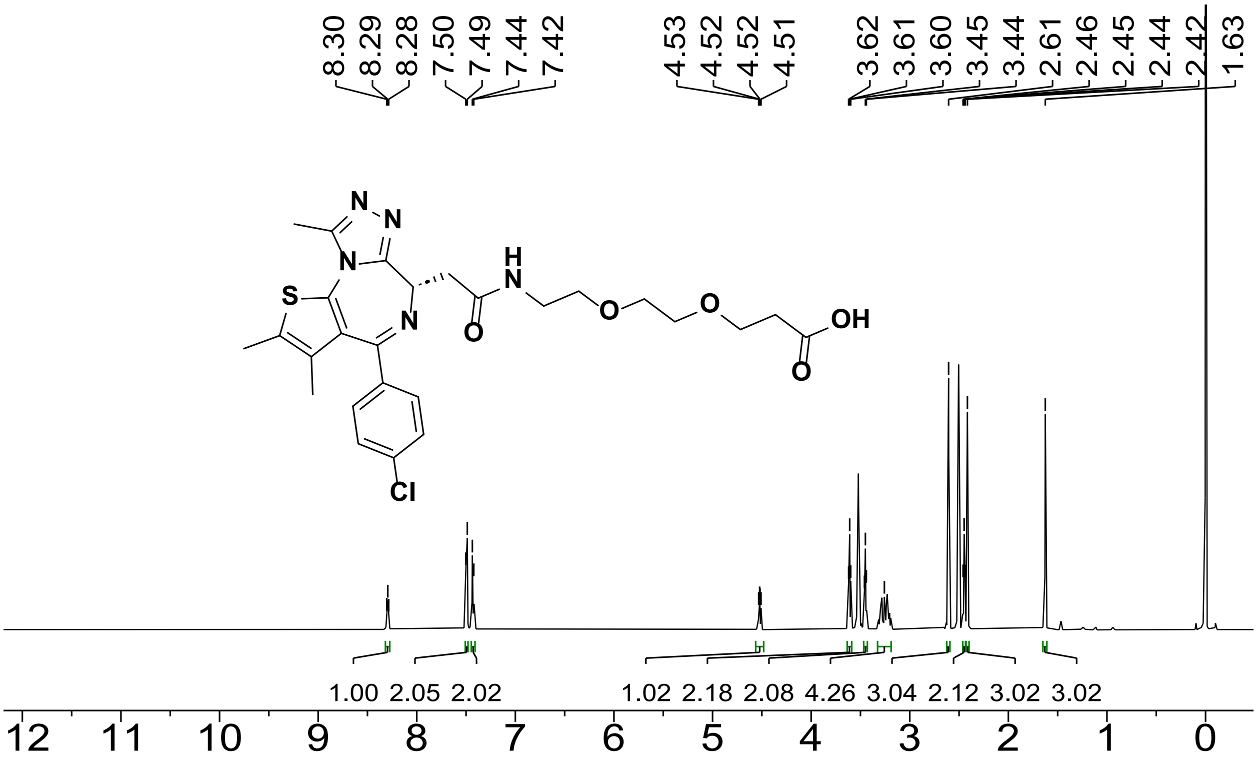

Figure S21. $^{1}$H NMR spectrum of **S20** (600 MHz, DMSO-$d_6$, 298 K).

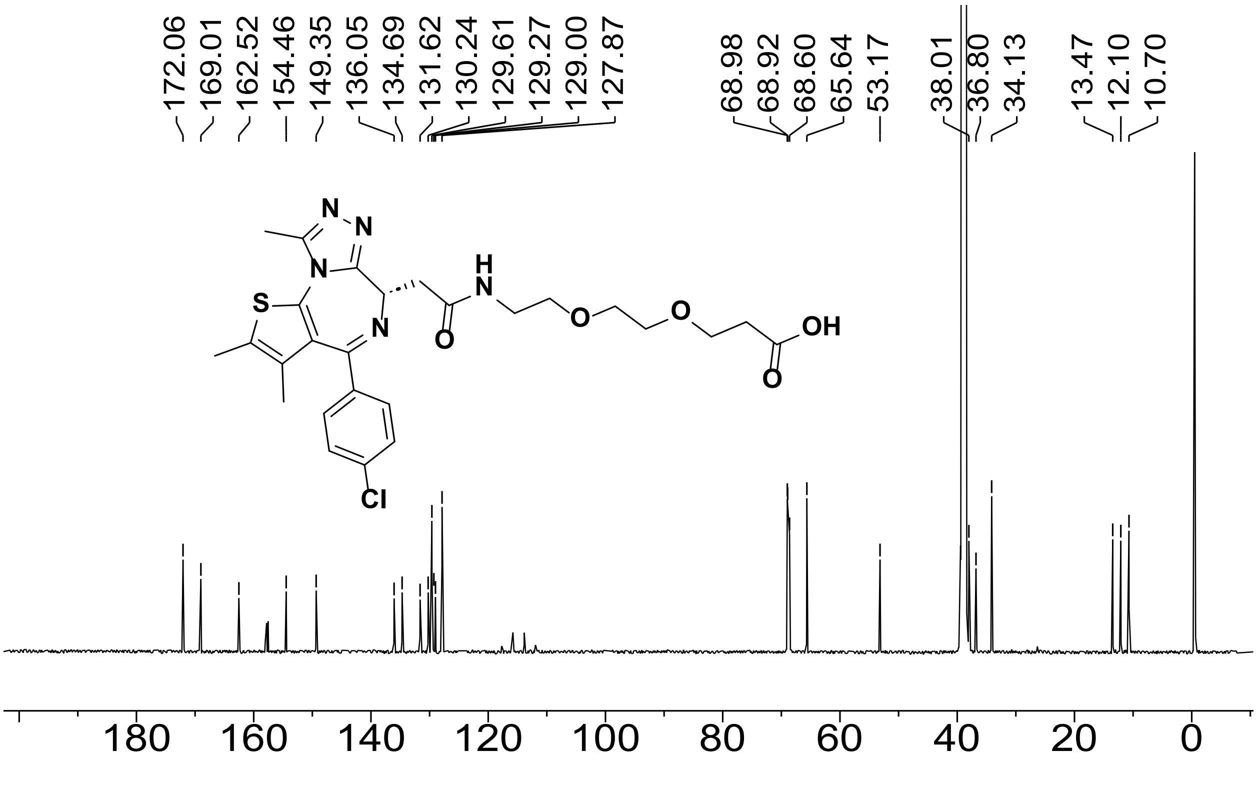

Figure S22. $^{13}$C NMR spectrum of **S20** (151 MHz, DMSO-$d_6$, 298 K).

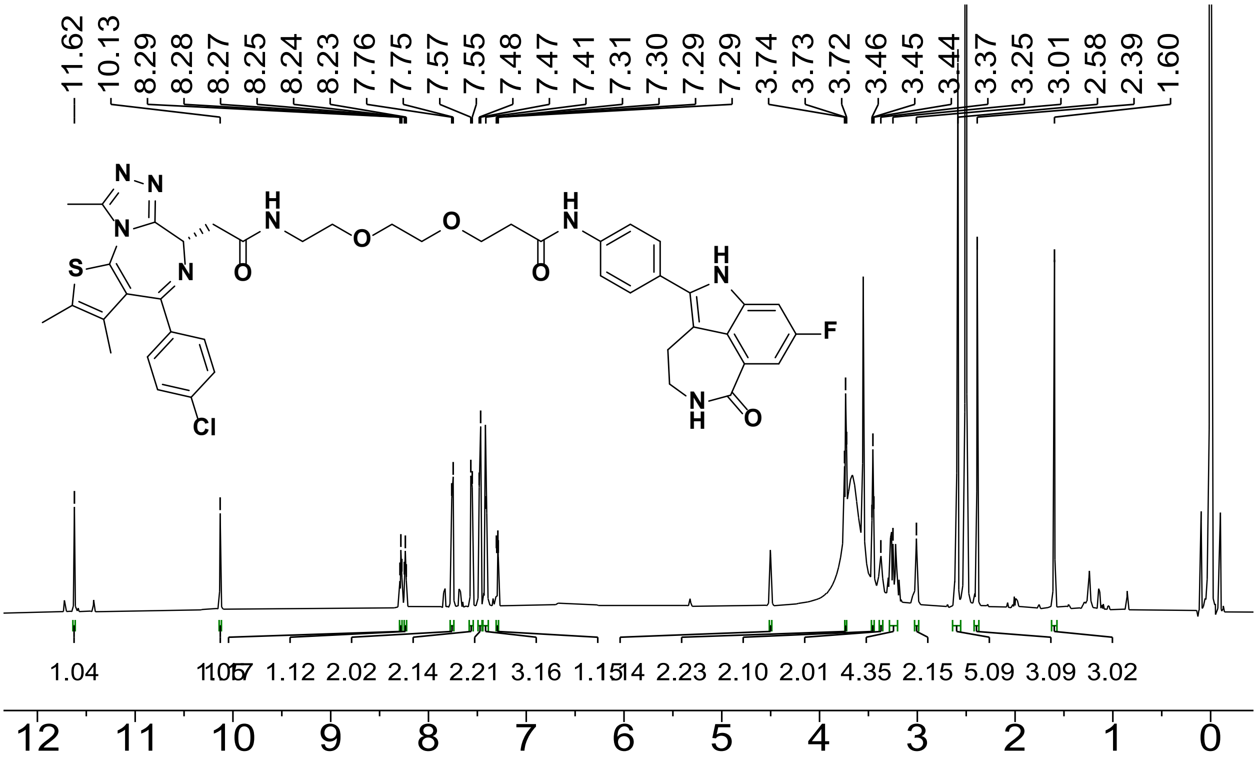

Figure S23. [1]H NMR spectrum of **S22**, compound **3** (600 MHz, DMSO-$d_6$, 298 K).

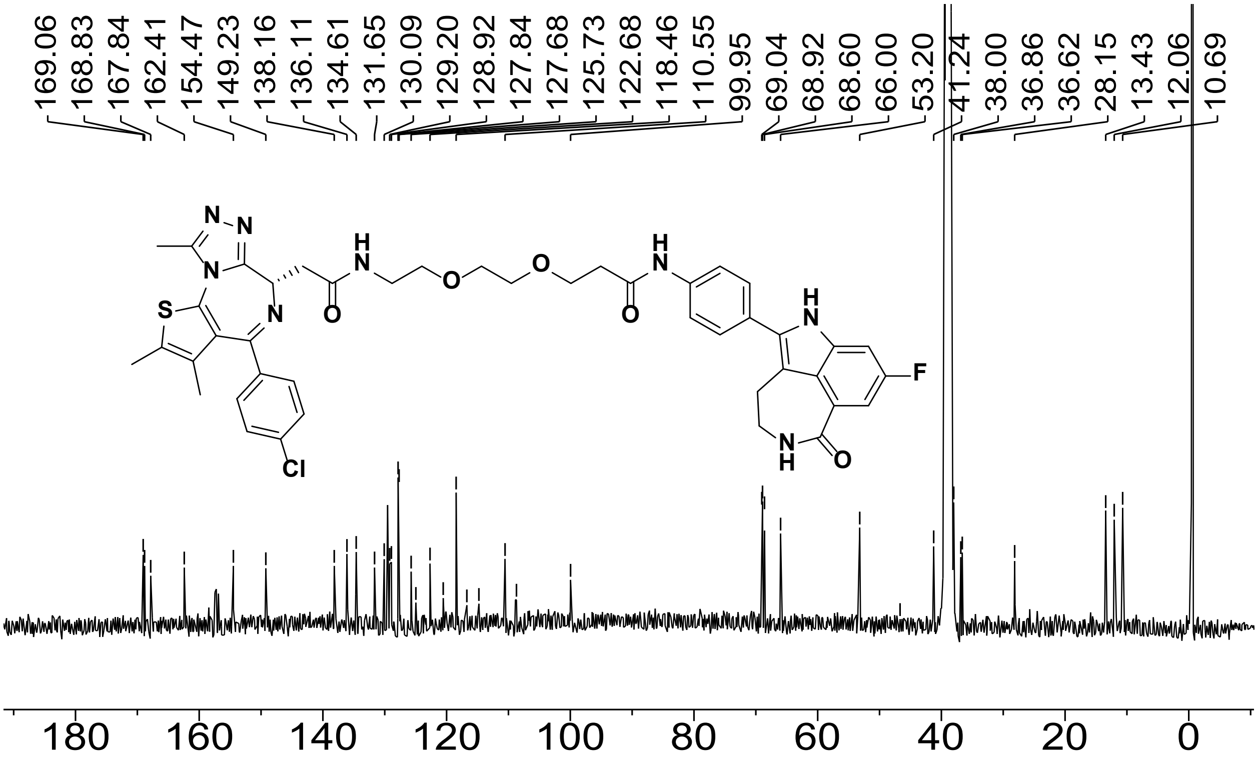

Figure S24. [13]C NMR spectrum of **S22**, compound **3** (151 MHz, DMSO-$d_6$, 298 K).

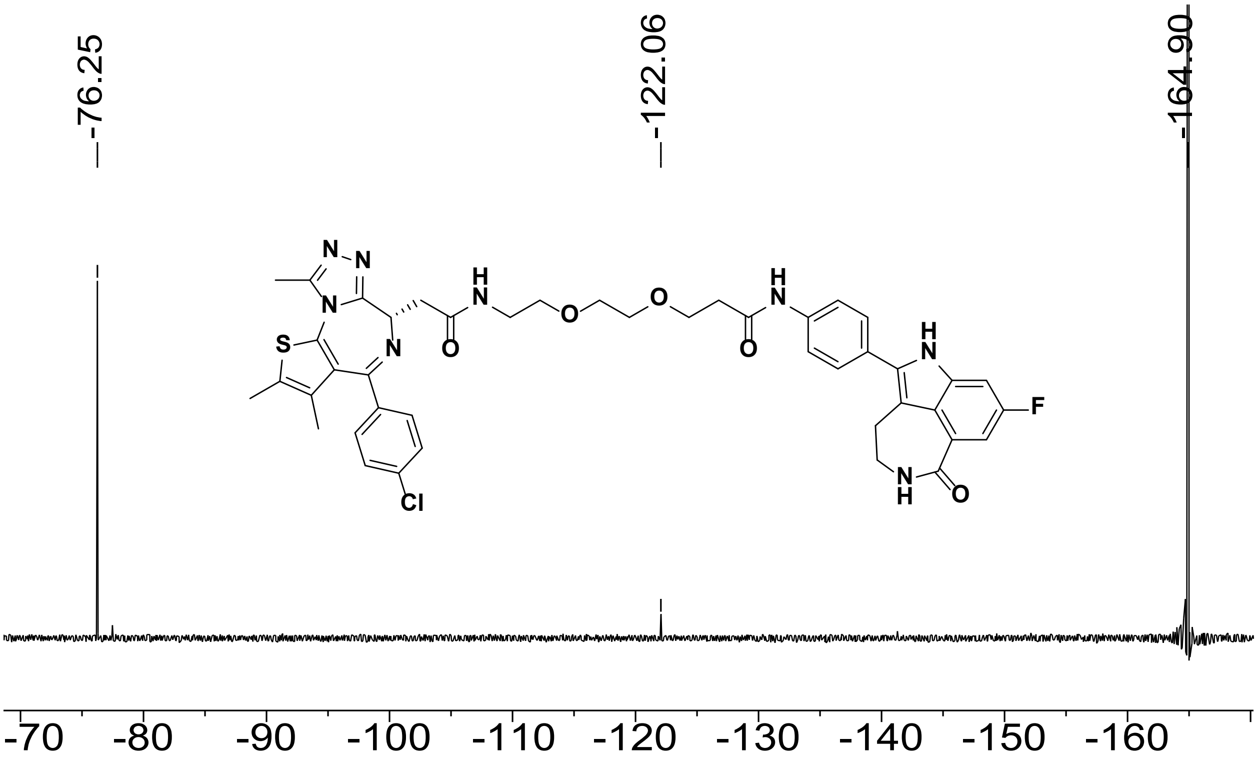

Figure S25. [19]F NMR spectrum of **S22**, compound **3** (376 MHz, DMSO-$d_6$, referenced to $C_6F_6$, 298 K).

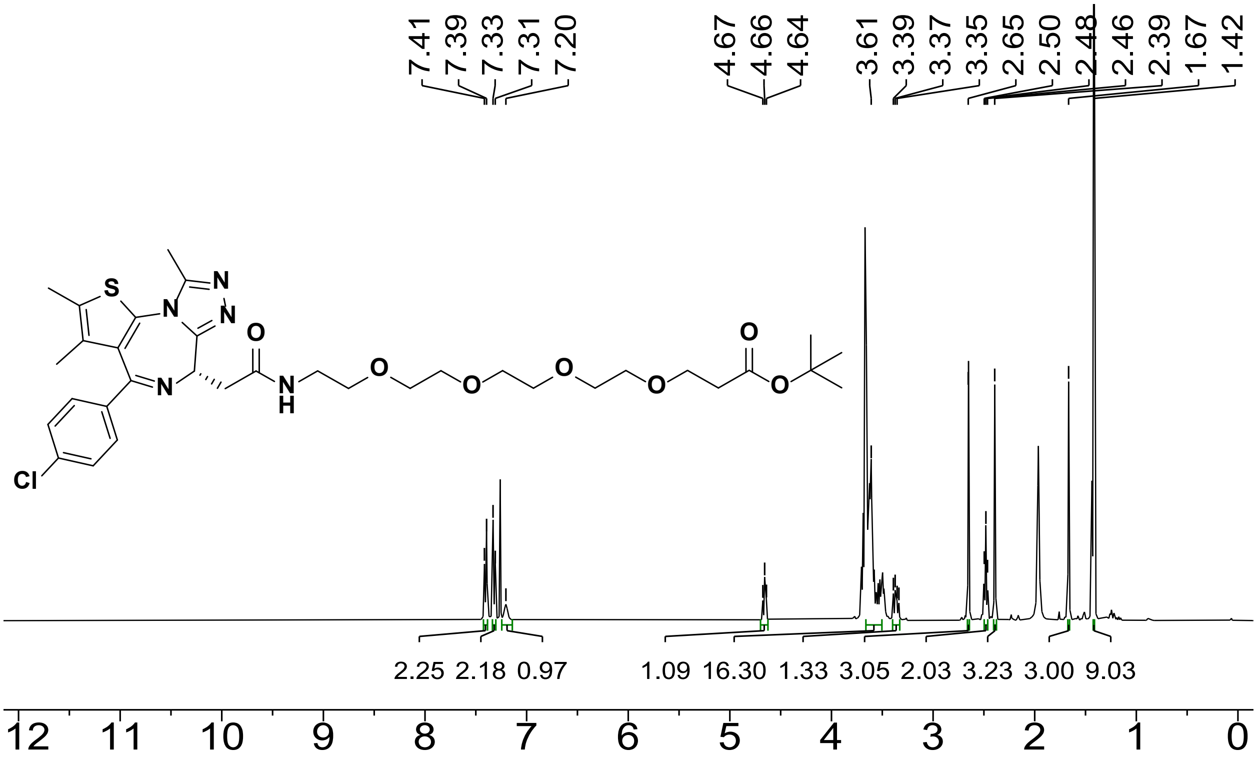

Figure S26. [1]H NMR spectrum of **S19** (600 MHz, $CDCl_3$, 298 K).

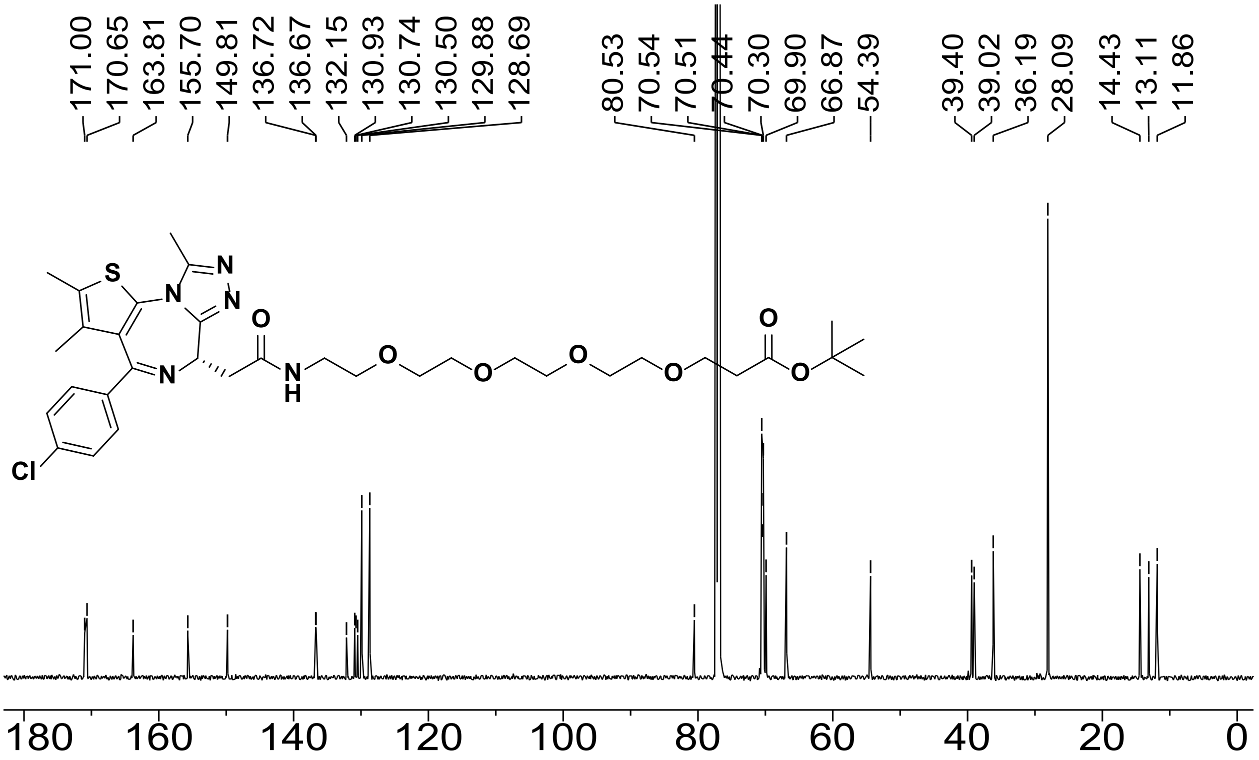

Figure S27. [13]C NMR spectrum of **S19** (151 MHz, $CDCl_3$, 298 K).

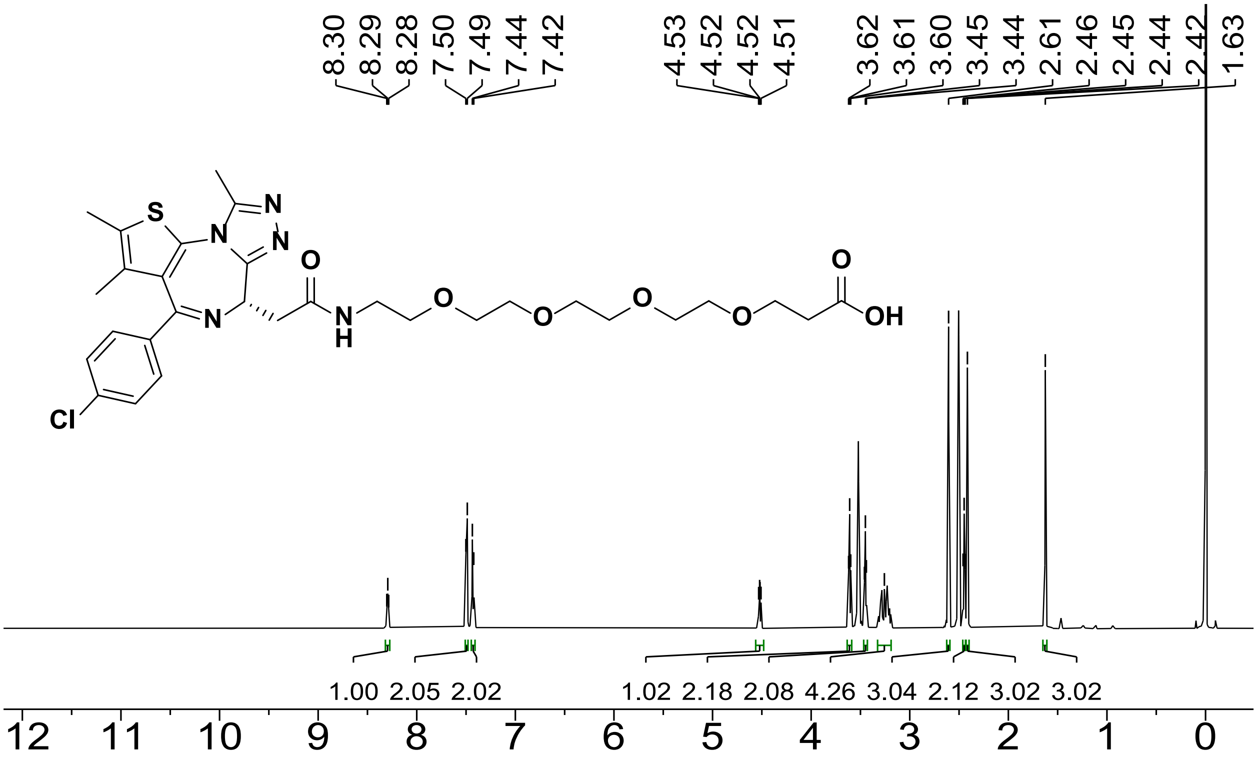

Figure S28. [1]H NMR spectrum of **S21** (600 MHz, DMSO-$d_6$, 298 K).

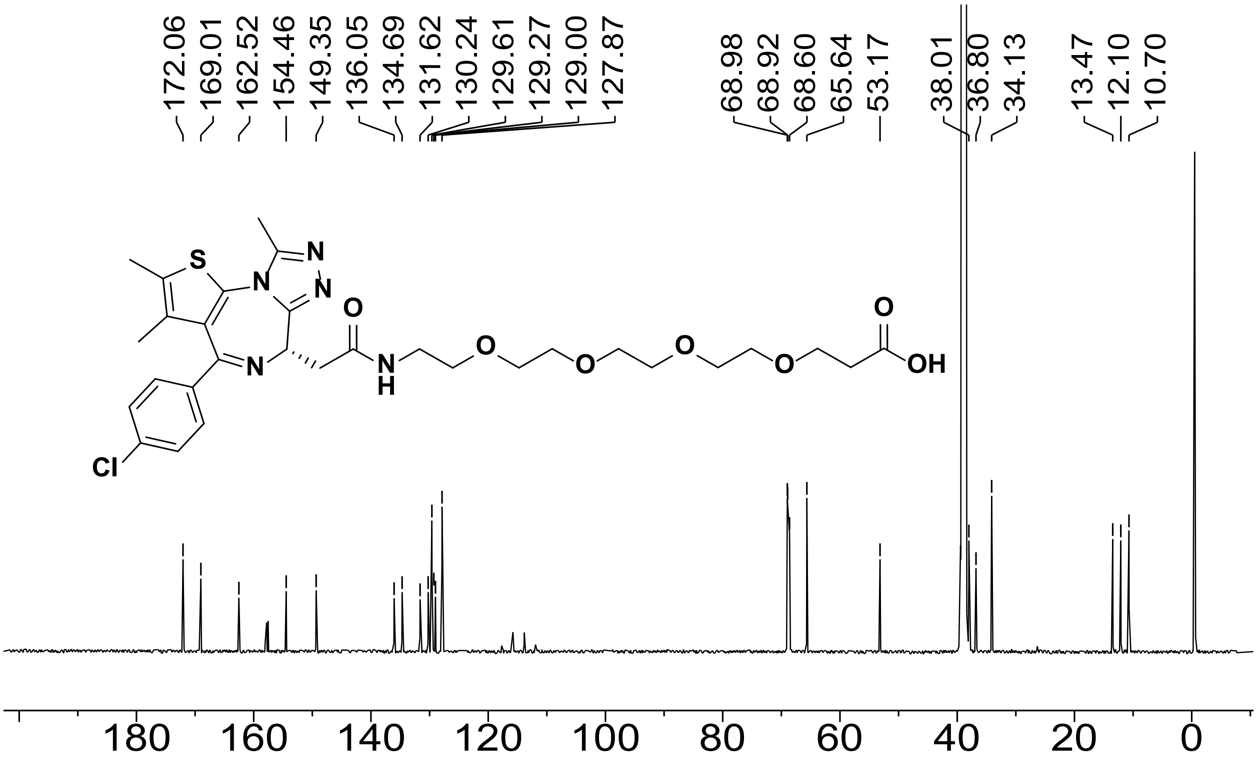

Figure S29. $^{13}$C NMR spectrum of **S21** (151 MHz, DMSO-$d_6$, 298 K).

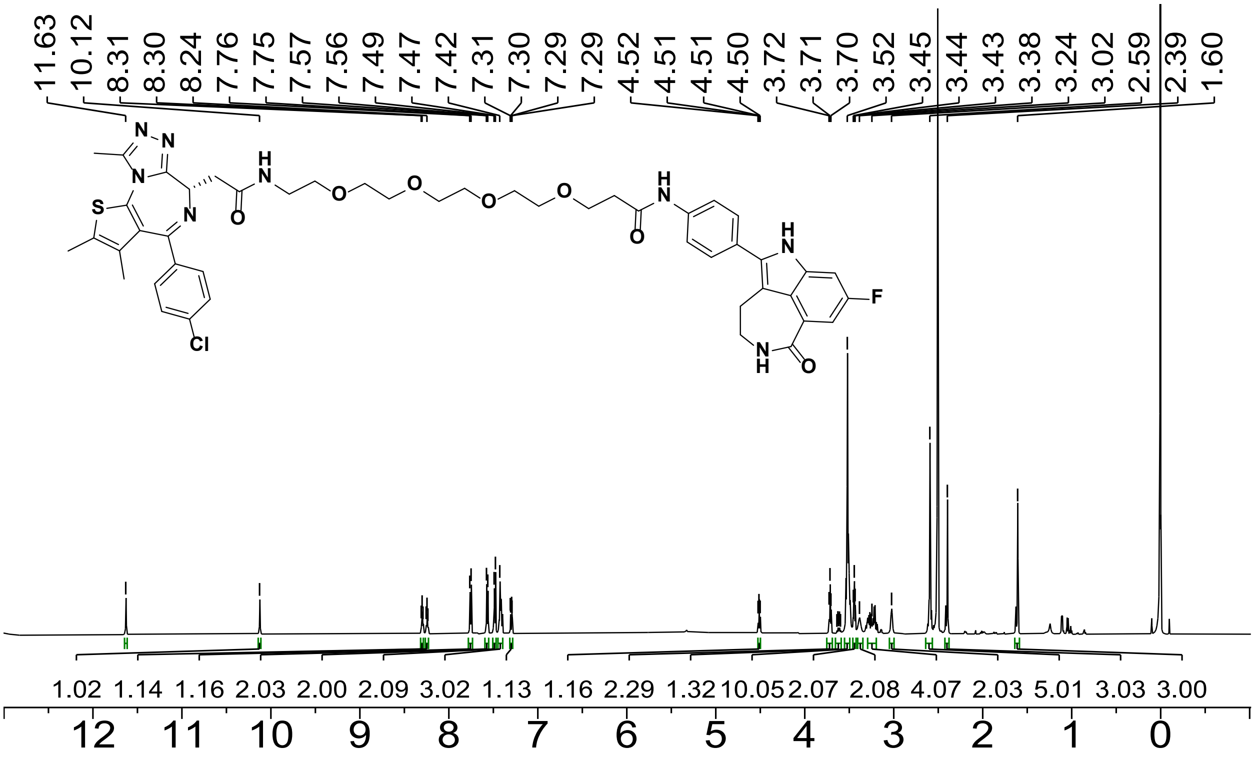

Figure S30. $^{1}$H NMR spectrum of **S23**, compound **4** (600 MHz, DMSO-$d_6$, 298 K).

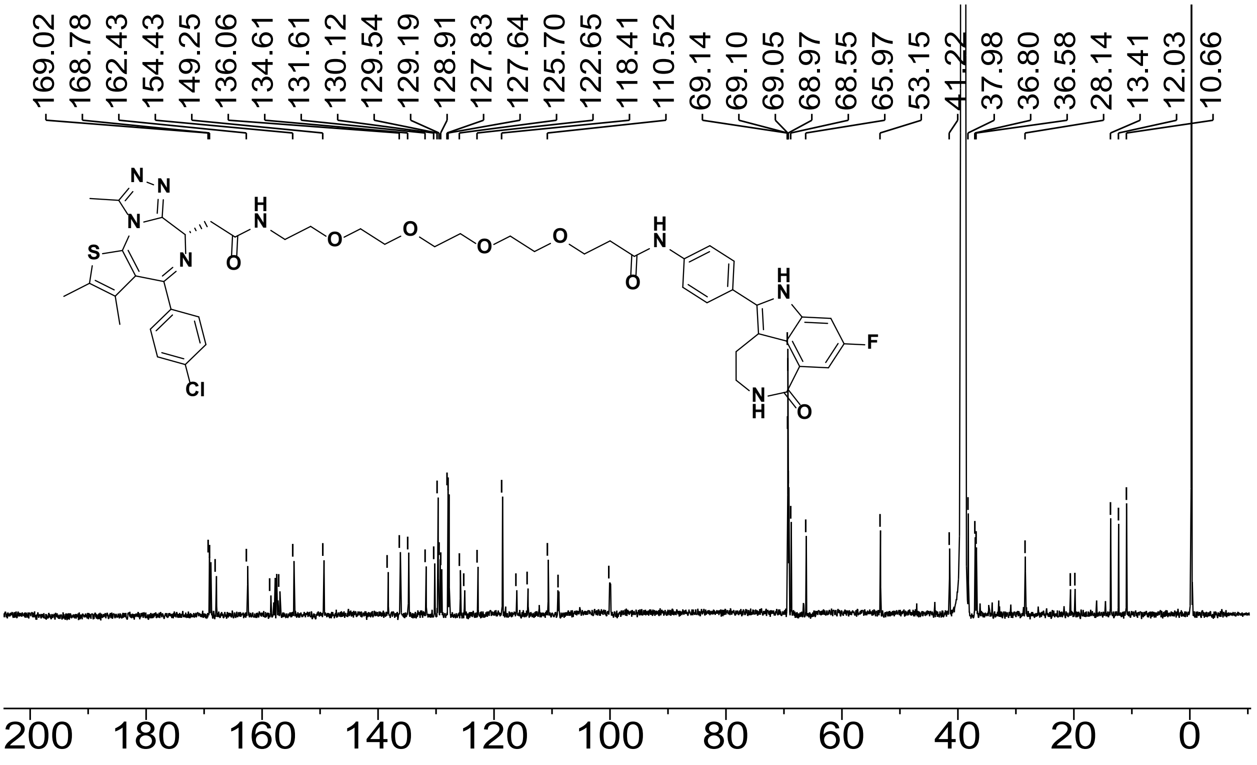

Figure S31. [13]C NMR spectrum of **S23**, compound **4** (151 MHz, DMSO-*d6*, 298 K).

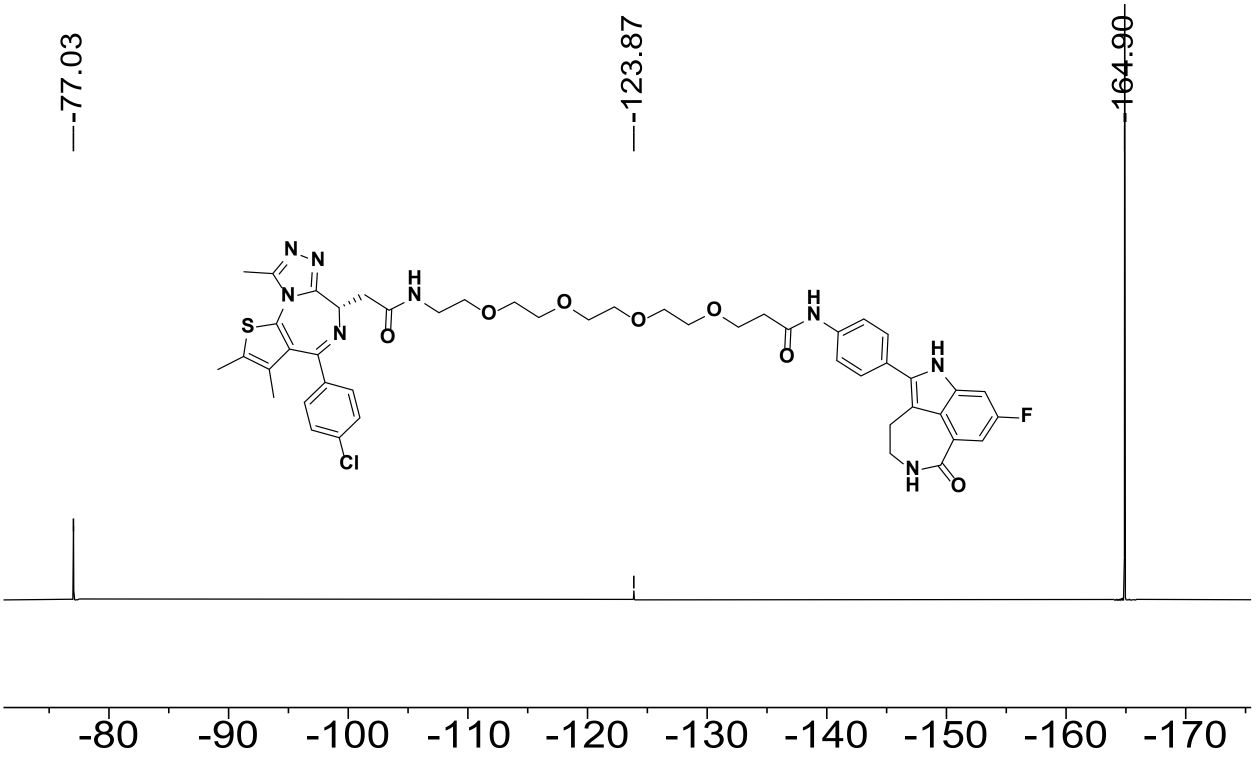

Figure S32. [19]F NMR spectrum of **S23**, compound **4** (376 MHz, DMSO-*d6*, referenced to $C_6F_6$, 298 K).

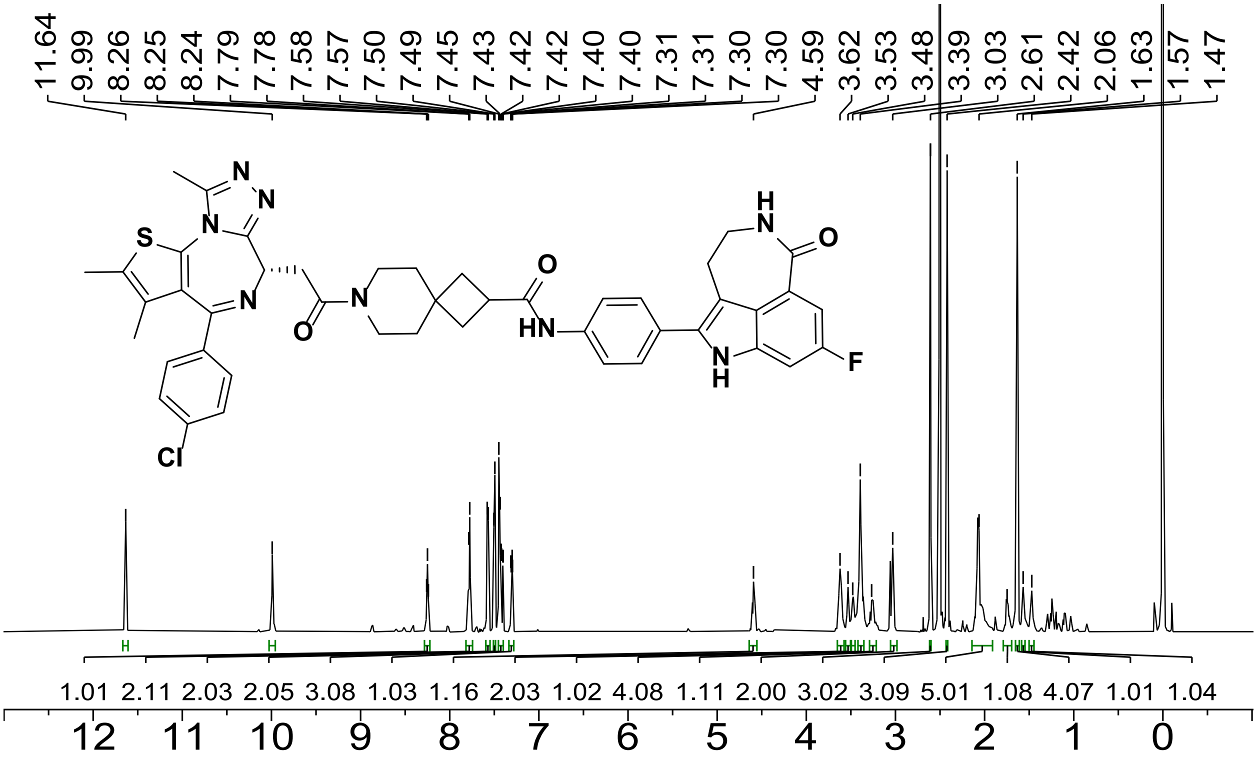

Figure S33. [1]H NMR spectrum of **S27**, compound **5** (600 MHz, DMSO-$d_6$, 298 K).

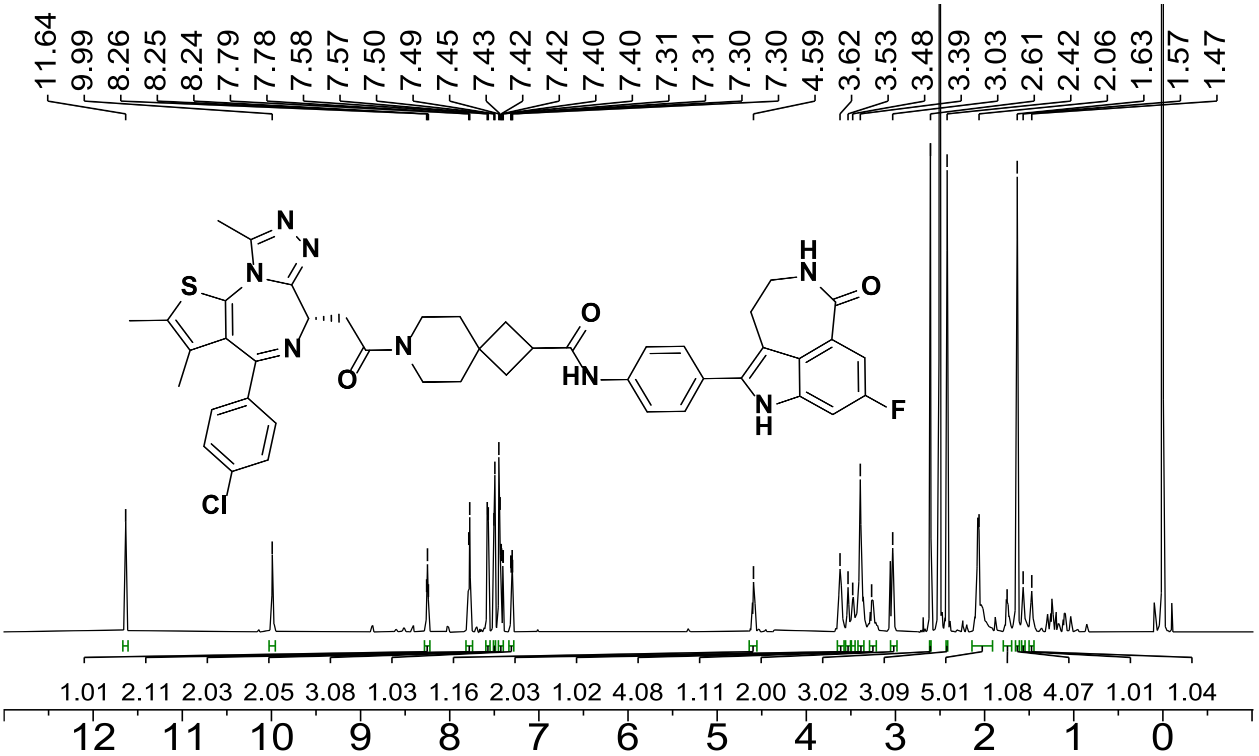

Figure S34. [13]C NMR spectrum of **S27**, compound **5** (151 MHz, DMSO-$d_6$, 298 K).

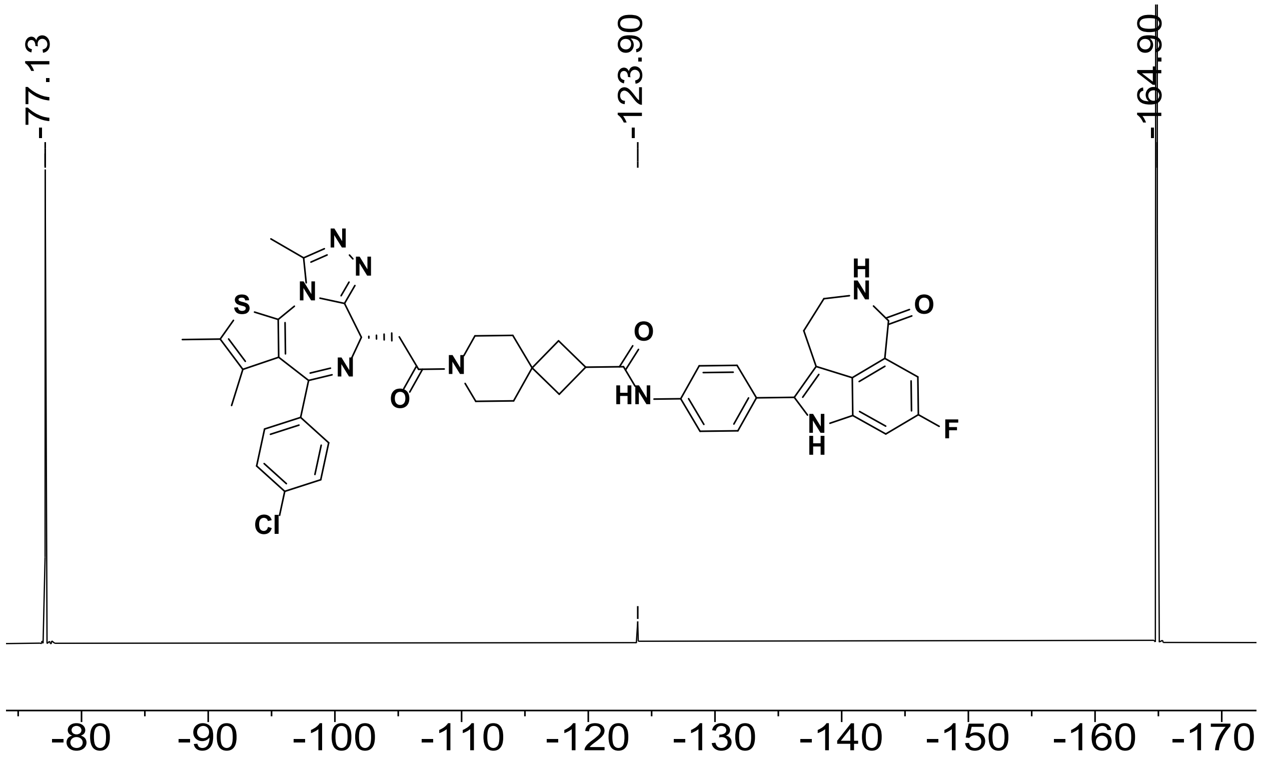

Figure S35. [19]F NMR spectrum of **S27**, compound **5** (376 MHz, DMSO-$d_6$, referenced to $C_6F_6$, 298 K).

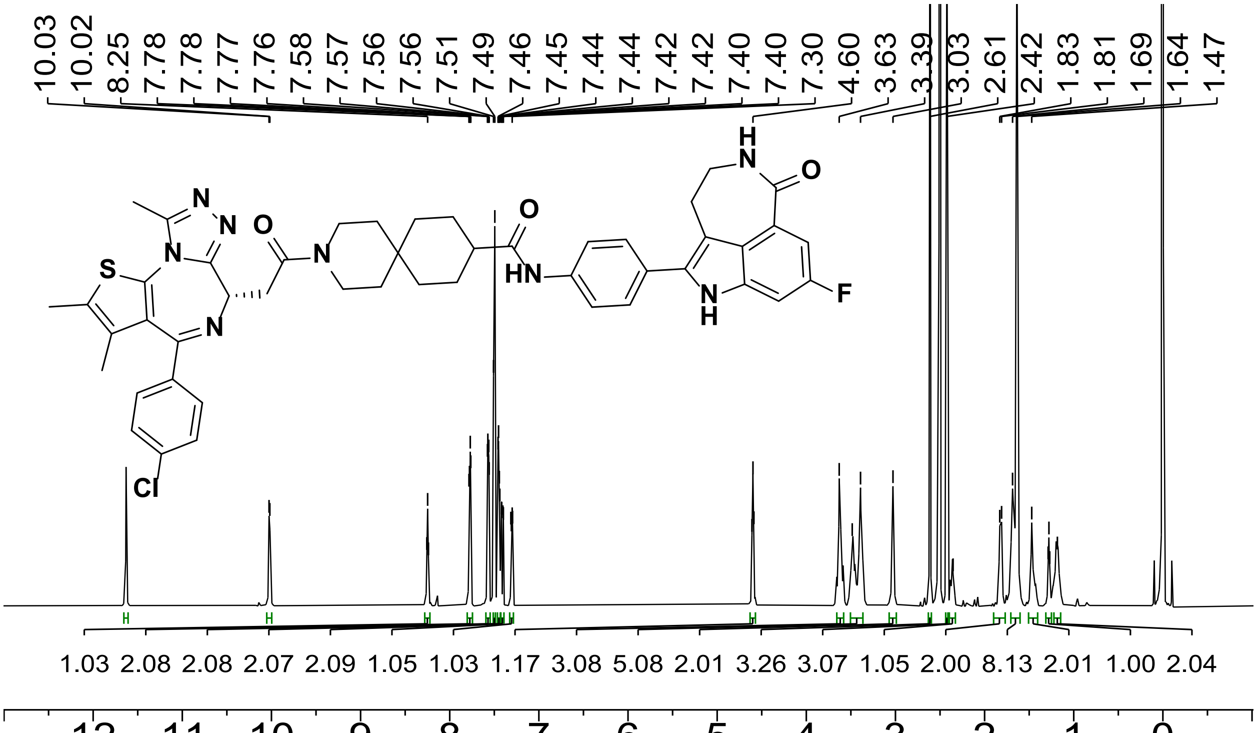

Figure S36. [1]H NMR spectrum of **S31**, compound **6** (600 MHz, DMSO-$d_6$, 298 K).

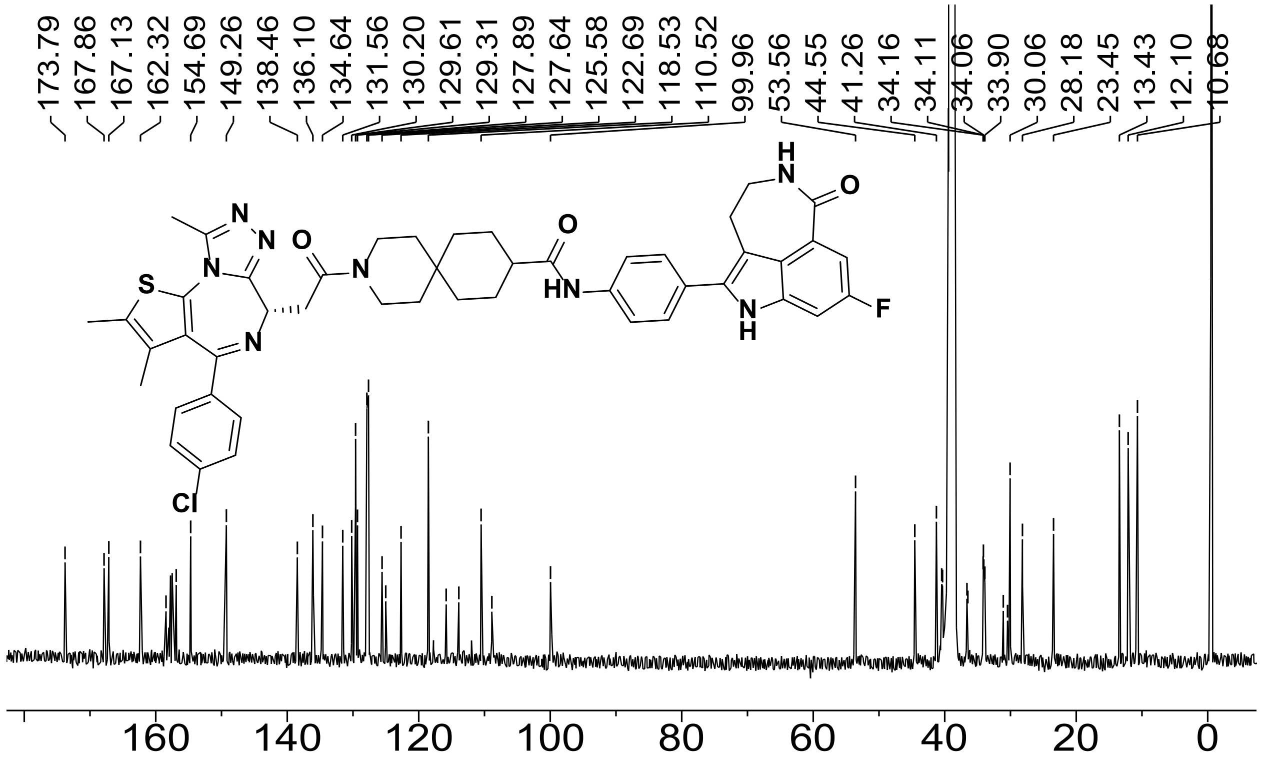

Figure S37. [13]C NMR spectrum of **S31**, compound **6** (151 MHz, DMSO-*d*$_6$, 298 K).

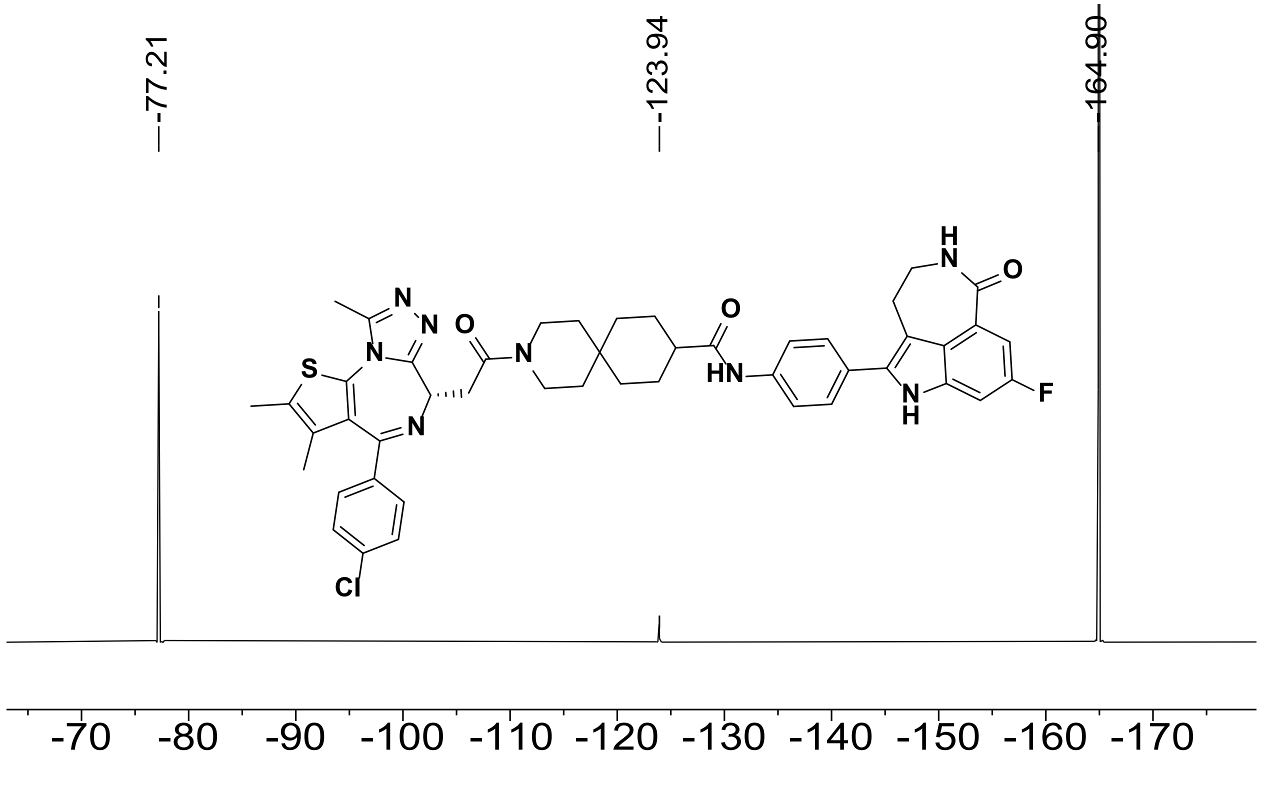

Figure S38. [19]F NMR spectrum of **S31**, compound **6** (376 MHz, DMSO-*d*$_6$, referenced to $C_6F_6$, 298 K).

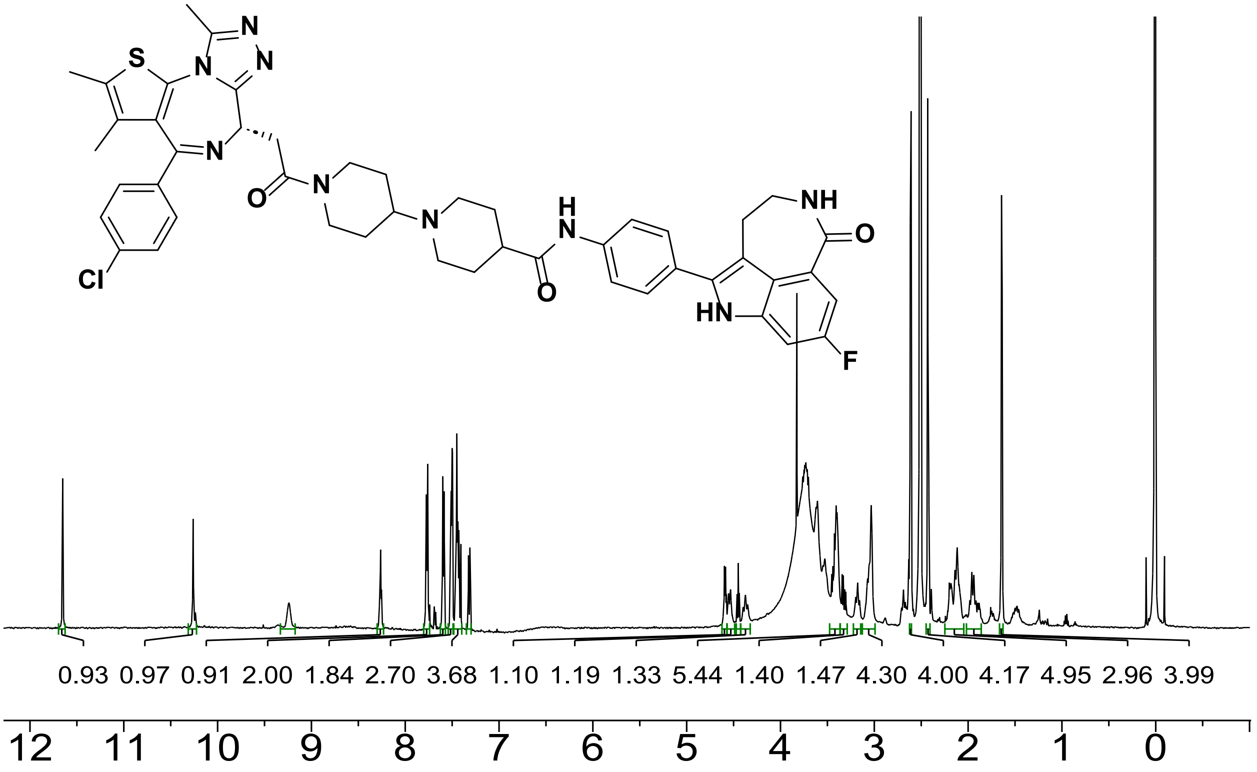

Figure S39. [1]H NMR spectrum of **S35**, compound **7** (600 MHz, DMSO-$d_6$, 298 K).

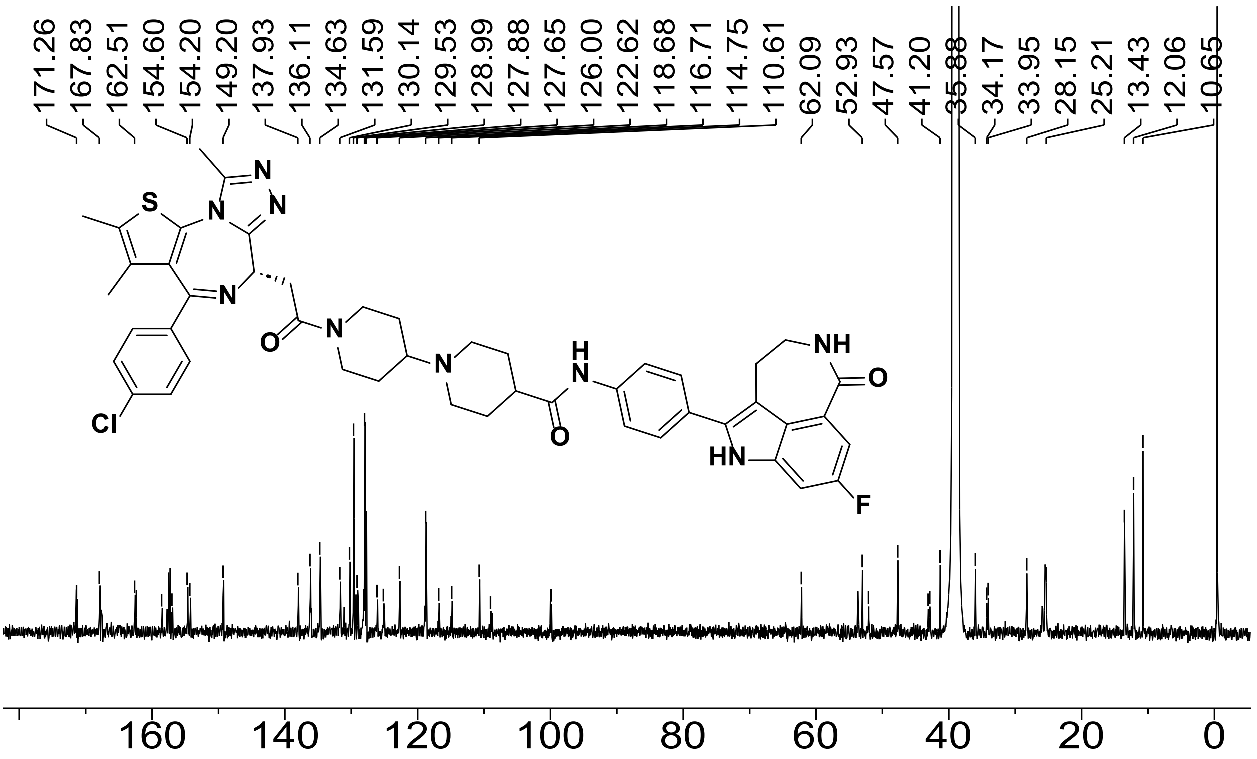

Figure S40. [13]C NMR spectrum of **S35**, compound **7** (151 MHz, DMSO-$d_6$, 298 K).

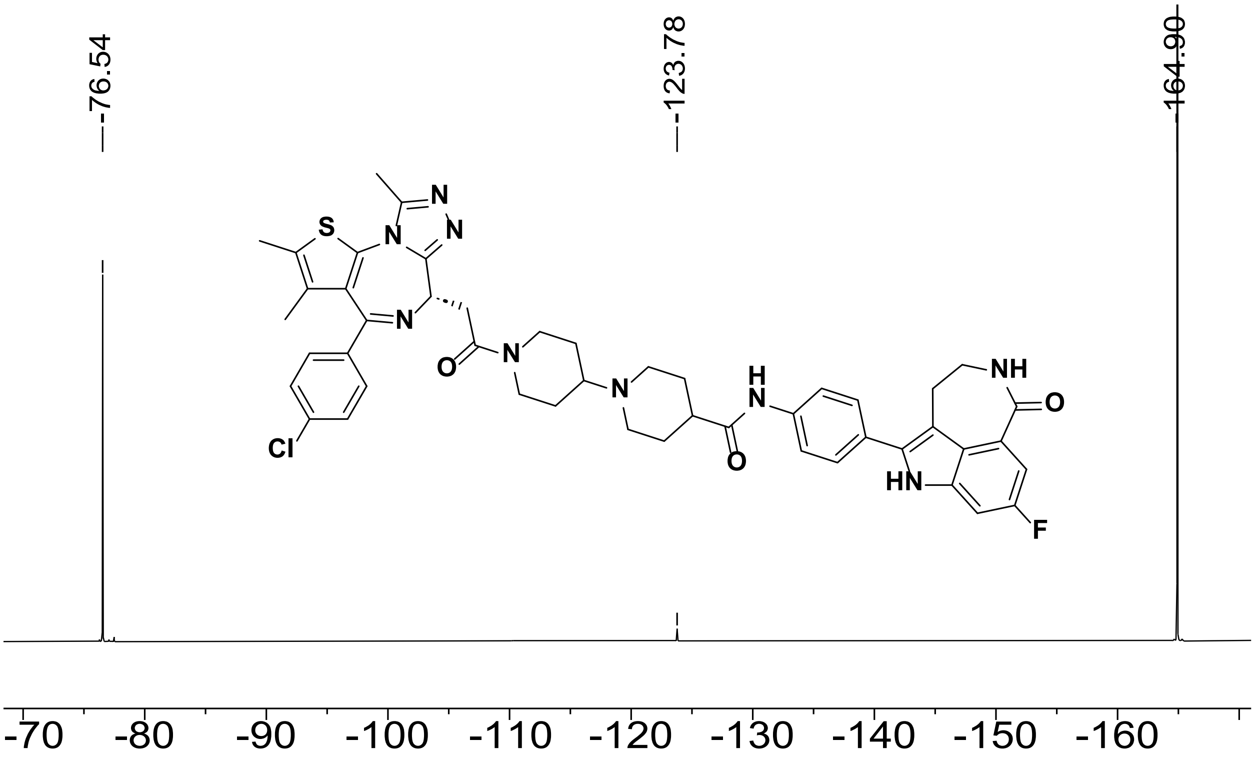

Figure S41. $^{19}$F NMR spectrum of **S35**, compound **7** (376 MHz, DMSO-$d_6$, referenced to $C_6F_6$, 298 K).

**References**

1. Zimmermann, M. *et al.* CRISPR screens identify genomic ribonucleotides as a source of PARP-trapping lesions. *Nature* **559**, 285–289 (2018).

2. Yu, J. *et al.* Structure-based design, synthesis, and evaluation of inhibitors with high selectivity for PARP-1 over PARP-2. *Eur. J. Med. Chem.* **227**, 113898 (2022).
